# CRISPR RiPCA for Investigating eIF4E-m^7^GpppX Capped mRNA Interactions

**DOI:** 10.1101/2025.06.19.660603

**Authors:** Gabriela Vega-Hernández, Jesse Duque, Brandon J. C. Klein, Dalia M. Soueid, Jason C. Rech, Hui Wang, Wenhui Zhou, Amanda L. Garner

**Affiliations:** Program in Chemical Biology, University of Michigan, Ann Arbor, Michigan 48109, United States; Michigan Center for Therapeutic Innovation, Michigan Medicine, University of Michigan, Ann Arbor, Michigan 48109, United States; Department of Medicinal Chemistry, College of Pharmacy, University of Michigan, Ann Arbor, Michigan 48109, United States; Promega Corporation, San Luis Obispo, California 93401, United States

## Abstract

Post-transcriptional modifications expand the information encoded by an mRNA. These dynamic and reversible modifications are specifically recognized by reader RNA-binding proteins (RBPs), which mediate the regulation of gene expression, RNA processing, localization, stability, and translation. Given their crucial functions, any disruptions in the normal activity of these readers can have significant implications for cellular health. Consequently, the dysregulation of these RBPs has been associated with neurodegenerative disorders, cancers, and viral infections. Therefore, there has been growing interest in targeting reader RBPs as a potential therapeutic strategy since developing molecules that restore proper RNA processing and function may offer a promising avenue for treating diseases. In this work, we coupled our previously established live-cell RNA-protein interaction (RPI) assay, RNA interaction with Protein-mediated Complementation Assay (RiPCA), with CRISPR technology to build a new platform, CRISPR RiPCA. As a model for development, we utilized the interaction of eukaryotic translation initiation factor 4E (eIF4E), a reader RBP that binds to the m^7^GpppX cap present at the 5′ terminus of coding mRNAs, with an m^7^G capped RNA substrate. Using eIF4E CRISPR RiPCA, we demonstrate our technology’s potential for measuring on-target activity of inhibitors of the eIF4E RPI of relevance to cancer drug discovery.

## INTRODUCTION

RNAs can undergo >170 post-transcriptional modifications (**Figure 1A**).^1,2^ These chemical modifications, collectively referred to as the epitranscriptome, can change the post-transcriptional fate of a target RNA.^1,2^ The addition and removal of RNA modifications are catalyzed by “writer” and “eraser” RNA-modifying proteins (RMPs), respectively. Key regulators of these modifications include “reader” RNA-binding proteins (RBPs), which specifically recognize and bind to the modified RNA substrates and regulate localization, stabilization, translation, turnover, and silencing (**Figure 1B**).^1,2^ The dysregulation of these biological processes, altered post-transcriptional modifications within messenger RNA (mRNA), and sequence variations that interfere with epitranscriptomic marks have been associated with metabolic and neurological diseases and cancers.^1,3,4^ As a result, therapeutics that manipulate the landscape of RNA modifications, both targeting RMPs and reader RBPs, are actively being explored in the field to rescue pathological phenotypes.^2,5,6^ Excitingly, a small molecule inhibitor of METTL3, an RMP that catalyzes the *N*^6^-methyladenosine (m^6^A) modification of mRNAs, has recently been advanced to clinical trials in patients with advanced solid tumors.^7–9^ This development highlights the growing potential of targeting the epitranscriptome for cancer therapy.^1,3,10^

**Figure 1.**
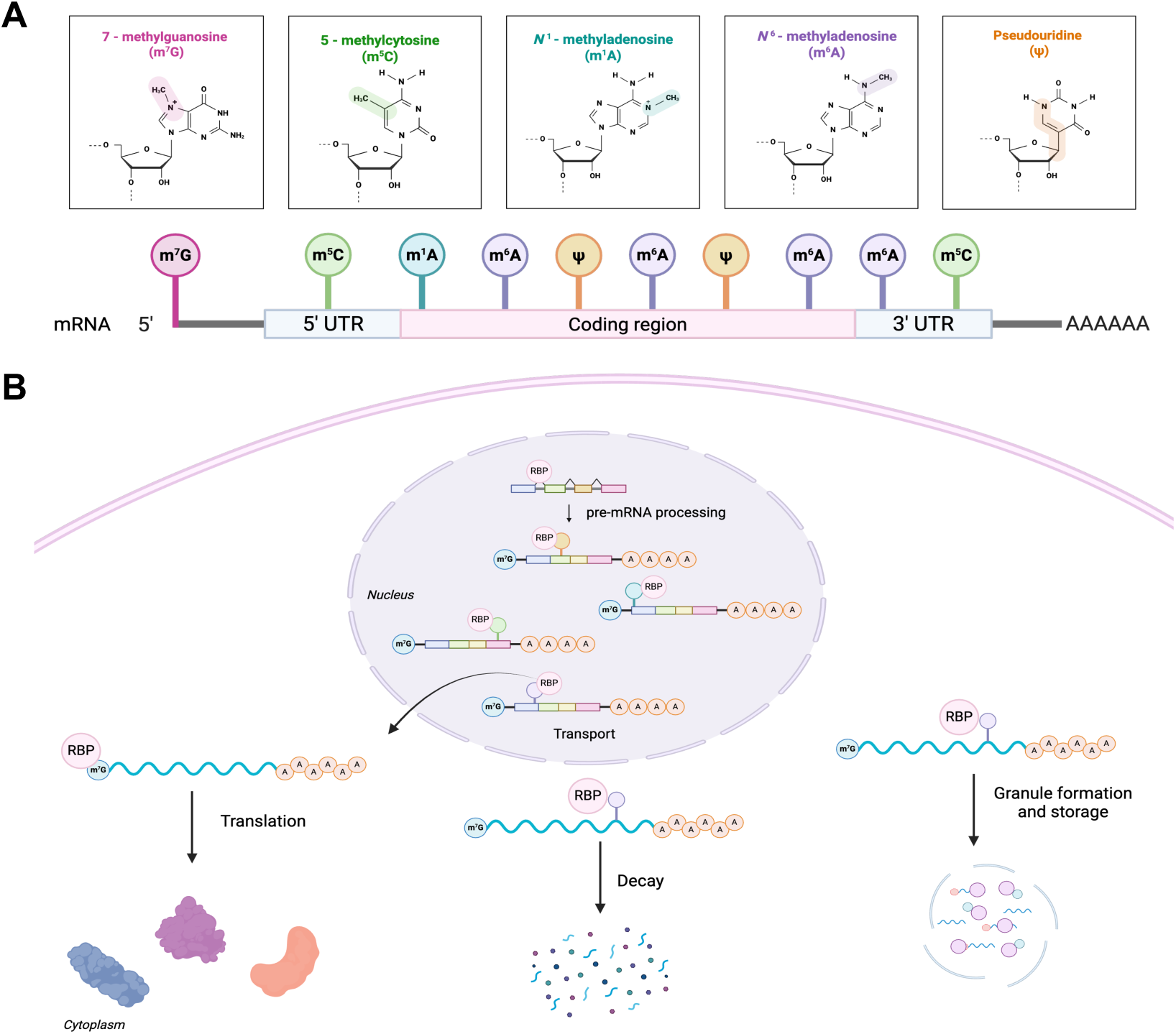
Regulation of the epitranscriptome by RNA-binding proteins (RBPs). (A) Select examples of RNA modifications on an mRNA transcript. (B) Reader RBPs regulate many aspects of mRNA biology.

A reader RBP that plays a global role in protein synthesis is the eukaryotic translation initiation factor 4E (eIF4E), which binds to the 5′ m^7^GpppX cap of mRNAs to initiate cap-dependent translation.^11,12^ eIF4E activity is tightly regulated, primarily by its binding partner 4E-BP1, which inhibits its function. However, when 4E-BP1 is phosphorylated via the mTOR pathway, eIF4E is released and recruits other initiation factors to form the eIF4F complex, facilitating cap-dependent translation. Hyperactivation of eIF4E’s activity promotes the aberrant translation of mRNA transcripts encoding proliferation and survival-promoting proteins such as cyclins D1 and D3, c-Myc, VEGF, MDM2, and Bcl-2, important in driving many cancerous phenotypes. Accordingly, identifying modulators that target eIF4E activity is of high interest for therapeutic development. Characterization of inhibitors of the eIF4E RNA-protein interaction (RPI) has traditionally been carried out using *in vitro* biochemical methodologies such as fluorescence polarization assays and cellular target engagement assessed using methods like cellular thermal shift assays (CETSA).^13–15^ A method that could simultaneously assess binding and cellular target engagement would greatly streamline inhibitor characterization of this and other RPIs to enable the field of RBP-targeted drug discovery. To enable these efforts, herein, we report the adaptation and expansion of our lab’s **R**NA-**i**nteraction with **P**rotein-mediated **C**omplementation **A**ssay (RiPCA) technology, a live-cell assay to detect RPIs (**Figure 2A**), for eIF4E.^16–19^ Leveraging CRISPR/Cas9 gene editing and bioorthogonal chemistry, we describe an optimized RiPCA platform with applicability for detecting and monitoring temporal modulation of eIF4E with drug-like small molecule inhibitors.

**Figure 2.**
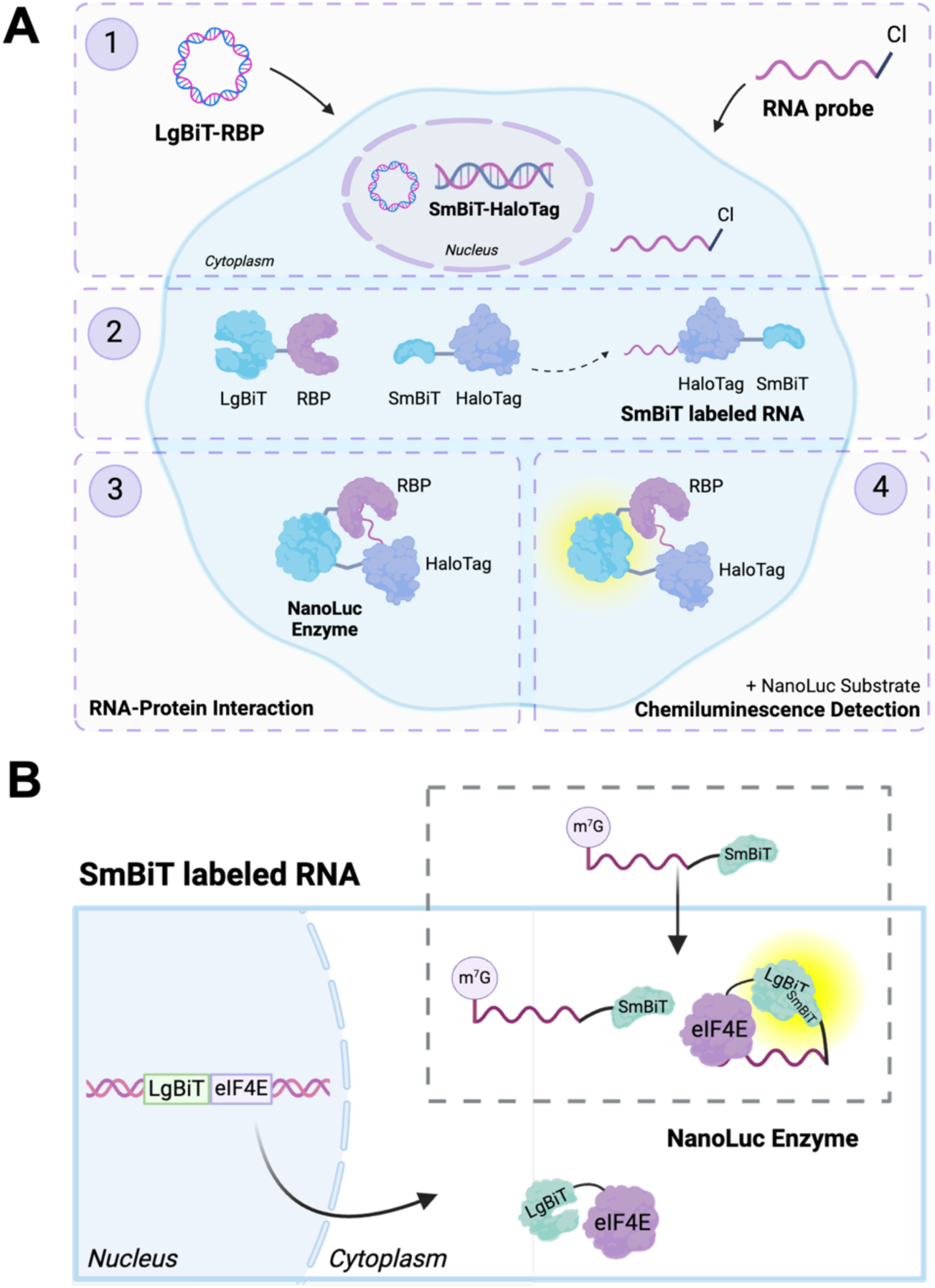
Live-cell detection of RNA-Protein Interactions (RPIs). (A) Previous work: RNA-interaction with Protein-mediated Complementation Assay (RiPCA) utilizing transfection of LgBiT-tagged RBP plasmid and RNA substrate. (B) This work: CRISPR RiPCA using endogenously LgBiT-tagged RBP.

## RESULTS AND DISCUSSION

### RiPCA 2.0 for eIF4E: design and challenges

Existing methods to detect RPIs and assess inhibition often rely on purified systems which lack physiological relevance because they are conducted outside of the cell and cannot replicate its complex environment or native interactions. ^6,19,20^ Current cell-based methods for detecting RPIs generally rely on fluorescent-based reporters^21–24^ or proximity-dependent labeling.^25–30^ While these methods are effective for detecting RPIs in their native cellular context, they are unsuitable for evaluating the inhibitory effects of small molecules due to their irreversible nature of detection. To enable RPI detection in living cells, our lab developed an assay called RNA-interaction with Protein-mediated Complementation Assay (RiPCA) (**Figure 2A**).^16–18^ RiPCA utilizes a split NanoLuciferase (NanoLuc^®^) enzyme,^31^ whose small subunit (SmBiT) and large subunit (LgBiT) have low affinity for each other (K_d_ = 190 μM), which ensures that assembly of the NanoLuc^®^ enzyme is driven by the RPI. This reversibility also allows modulators of the RPI to be evaluated. RiPCA has been a successful system for the detection of RPIs involving microRNAs and mRNA motifs with their respective RBPs.^16,17,19^ To determine RiPCA’s applicability for the detection of the interaction of modified RNAs with reader RBPs, we utilized eIF4E’s interaction with m^7^GpppX capped mRNA, as this interaction is currently being explored by our laboratory as a potential therapeutic target.^12–14^

To initiate the development of eIF4E RiPCA, we identified an RNA substrate previously reported to bind to the RBP with an affinity of ∼39 nM determined by intrinsic fluorescence measurements.^32^ Based on these findings, a 10-nt RNA oligo containing a 5′ 7-methylguanosine-triphosphate (m^7^GTP) cap and 3′ C7 amino linker was designed to test as eIF4E’s RNA substrate in RiPCA (**Figure 3A** and **Table S1**). Additionally, we designed two negative control RNAs with the same sequence as the m^7^GTP-containing RNA, which either lacked 7-methylation of the guanine or the entire 5′ to 5′ guanine modification (**Figure 3A** and **Table S1**). The RNA probes were subsequently conjugated to HaloTag^®^ ligands containing 2, 4, or 6 PEG linker lengths (**Figure S1A**). To measure the binding affinity of the designed RNA substrates, a fluorescence polarization (FP) assay was employed.^13,14,33^ From this analysis, we demonstrated that the added linker and the modification on the RNA probe did not significantly disrupt interaction with eIF4E, with measured EC_50_ values of 35.5 (NH_2_), 131.5 (PEG_2_), 140.4 (PEG_4_), and 137.7 (PEG_6_) nM, respectively (**Figures 3B** and **S2**). Additionally, binding was highly selective as both negative control RNAs lacked activity in this assay (**Figures 3B** and **S2**). Subsequently, we confirmed our RNA substrate’s ability to be modified by the SmBiT-HaloTag protein and detected characteristic bands of the SmBiT-HaloTag-RNA complex (**Figure S3)**.

**Figure 3.**
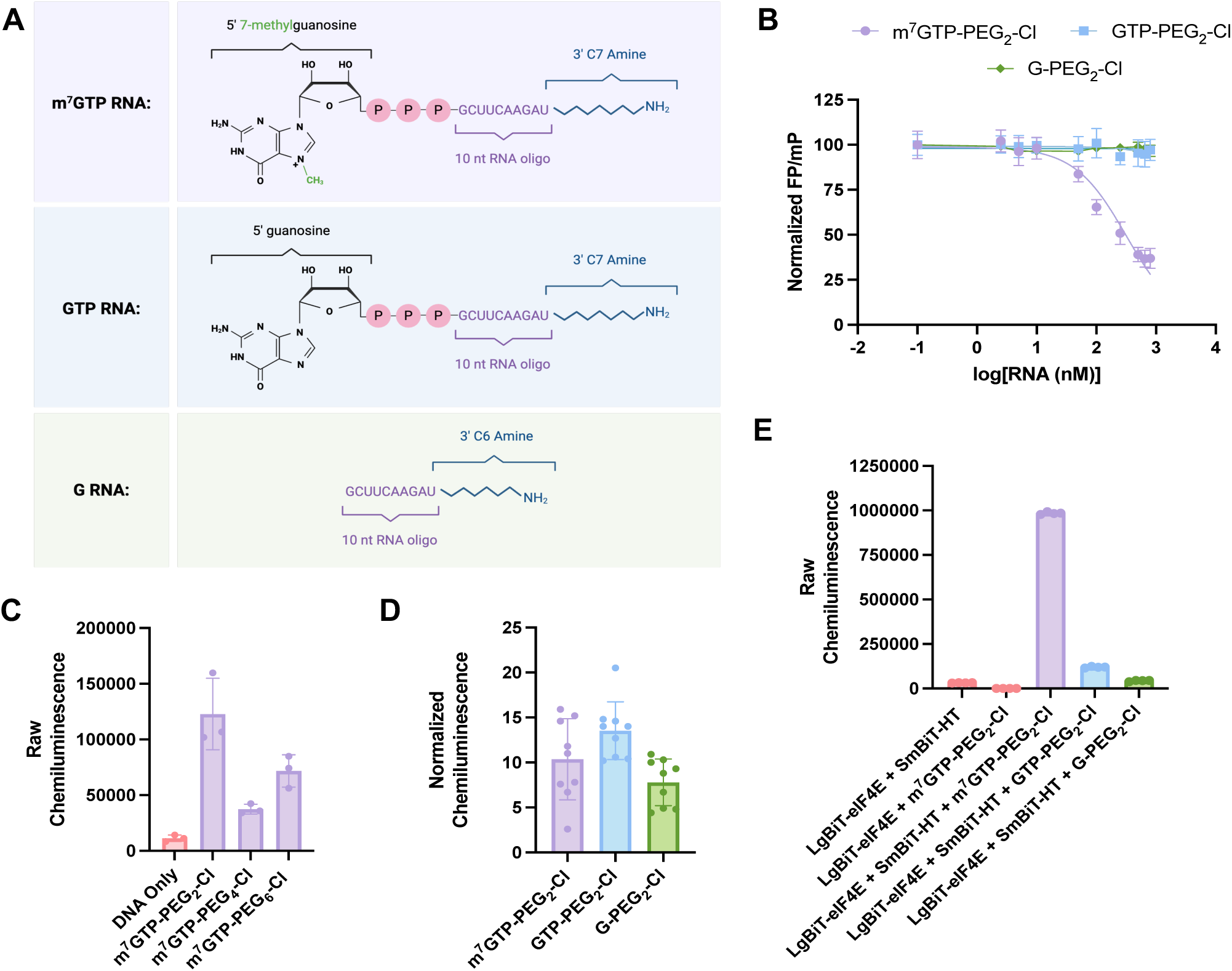
RiPCA 2.0 for eIF4E. (A) RNA sequences used for eIF4E RiPCA. (B) Characterization of RNA binding using an eIF4E fluorescence polarization assay. (C) Chemiluminescence signal generated in SmBiT-HaloTag cells transfected with LgBiT-eIF4E (2 ng/well) and m^7^GTP RNA substrates (100 nM). (D) Lack of specificity as determined by performing RiPCA using negative control RNAs. (E) Biochemical confirmation of the specificity of LgBiT-eIF4E binding to a m^7^GTP RNA RiPCA substrate.

After identifying an appropriate eIF4E-binding RNA substrate, we adapted the RiPCA 2.0 system for this RPI. eIF4E was first cloned into *N*- and *C*-terminal LgBiT plasmids. Test expression experiments demonstrated successful production of the *N*-terminally tagged LgBiT-eIF4E protein (**Figure S4**). Negligible expression of the *C*-terminal construct was not surprising, as the *C* terminus lies within a structured domain of eIF4E while the *N*-terminus is disordered.^34^ Upon testing LgBiT-eIF4E in RiPCA with our chloroalkane-labeled m^7^GTP RNA substrates, we observed a signal increase, suggesting successful RPI detection (**Figure 3C**). Preference for the shorter linker length was observed likely due to favorable proximity; thus, we conducted DNA and RNA titrations with this RNA and observed dose-dependent signal increases as expected (**Figure S5**). However, a critical issue arose when the negative control RNAs were tested at the same concentration range, wherein each produced similar chemiluminescence signal as the positive control RNA, thus raising concern that the signal detected was due to non-specific LgBiT/SmBiT assembly rather than being driven by the RPI (**Figure 3D**).

To test the ability of LgBiT-eIF4E to form a specific ternary complex with SmBiT-HaloTag-labeled m^7^GTP RNA, we developed a biochemical version of RiPCA (**Figure S6A**).^35^ Using this assay, we were excited to observe a 30-fold enhancement in chemiluminescence signal for the m^7^GTP RNA over negative control RNAs and samples lacking SmBiT-HaloTag or the RNA (**Figure 3E**). Based on these results, which indicate the specificity of ternary complex formation with the m^7^GTP RNA, we posited that overexpression of LgBiT-eIF4E in SmBiT-HaloTag expressing cells may cause non-specific assembly of NanoLuc^®^ leading to signal with non-substrate RNAs. Of note, we previously observed a similar result during the development of a RiPCA 2.0 for AUF1.^19^ To overcome this specificity challenge, we chose to develop a new RiPCA system that would allow us to test RNA binding at physiological levels of RBP by endogenous LgBiT tagging using CRISPR/Cas9 engineering and simultaneously control the amount of SmBiT in cells through transfection to enable selective RPI detection.

### Development of eIF4E CRISPR RiPCA

To address the limitations associated with RiPCA 2.0 for eIF4E, we first engineered a LgBiT-eIF4E HEK293T cell line using CRISPR/Cas9. A guide RNA was designed to target upstream of the start codon of eIF4E along with a DNA donor template encoding LgBiT, intended to be integrated at the *N*-terminus (**Figure S7A** and **Table S2**). The ribonucleotide protein (RNP) complex and donor DNA were introduced to cells via electroporation. After 72 h, fluorescence-activated cell sorting (FACS) was conducted to isolate single cells expressing GFP-fused Cas9 (GFP-positive), which were then plated into 96-well plates (**Figure S7B**). After individual clones were grown, incorporation of the LgBiT tag was assessed by lytic detection using its high affinity 11-amino acid HiBiT binding partner (K_d_ = 0.7 nM) (**Figure S8A**).^36^ From this analysis, a single clone was identified (**Figure S8B**), and its genomic DNA was subsequently analyzed for on-target insertion into the *N*-terminus of eIF4E and homozygosity (**Figure S8C**), as well as off-target effects via whole-genome sequencing (**Figures S9−S11** and **Table S3**). Expression of LgBiT-eIF4E protein was confirmed via Western blot (**Figure S8D**) and confocal microscopy (**Figure S8E**). To functionally characterize the cells, we performed a cap pull-down assay to assess LgBiT-eIF4E binding to its protein binding partners, eIF4G and 4E-BP1, and determined its phosphorylation status. Importantly, LgBiT-eIF4E was found to maintain canonical regulation, giving confidence that these cells would be able to accurately measure eIF4E-mRNA interactions (**Figures S8F** and **S8G**).

We next proceeded to develop the eIF4E CRISPR RiPCA system. Unlike RiPCA 2.0, our CRISPR-engineered cells do not stably express SmBiT-HaloTag.^16–19^ As such, transfection of the DNA encoding this fusion protein is required for RNA labeling (**Figure 2A**). Unfortunately, this approach failed to detect the RPI in living cells (data not shown). This observation likely stems from a misalignment between SmBiT-HaloTag protein synthesis, RNA labeling, and RNA substrate stability. As an alternative strategy, we attempted to deliver the SmBiT-HaloTag-RNA RNP complex into our LgBiT-eIF4E HEK293T cells using the TransIT-X2^®^ RNP delivery system and electroporation; however, these approaches also failed (data not shown). To confirm that transfection efficiency was the limiting factor, we tested the lytic delivery of the SmBiT-HaloTag-RNA RNP conjugate. Before cell lysis, SmBiT-HaloTag-RNA complexes were formed by incubating PEG_2_-chloroalkane-modified m^7^GTP RNA, identified as optimal from our previous development efforts (**Figures 3C** and **S6B**), with purified SmBiT-HaloTag protein. RNPs were delivered at cell lysis, and chemiluminescence generated from the RPI was detected (**Figure S12A**). Excitingly, titration of the SmBiT-HaloTag protein and the m^7^GTP-PEG_2_-Cl RNA resulted in a significant increase in RPI signal over SmBiT-HaloTag negative control samples (**Figure S12B**). Moreover, we observed a “hook effect” indicative of saturation of the individual components,^37^ which disrupts the formation of the ternary complex. Combined, these results demonstrate the feasibility of RiPCA-mediated detection of the eIF4E RPIs and the potential of the lytic CRISPR RiPCA approach.

Success with lytic CRISPR RiPCA inspired us to reevaluate our approach for performing the assay in live cells. As transfection of the RNA-protein conjugate proved problematic, we hypothesized that conjugation of the SmBiT peptide to the RNA substrate, rather than conjugation using HaloTag^®^, would overcome this challenge, as this direct approach would abrogate the need for cellular SmBiT-HaloTag synthesis and RNA labeling. As an additional benefit, this strategy would allow us to control RNA and SmBiT levels more precisely in the assay to improve assay performance. Previous efforts demonstrated the utility of PEG-labeled SmBiT peptides in biological assay development;^38^ as such, we adapted our labeling chemistry to this approach. SmBiT peptide containing a PEG_8_ linker and a succinimidyl ester was synthesized and used to label our binding and non-binding RNA substrates (**Figure S1B**). Confirmation of RNA labeling efficiency was determined via gel electrophoresis (**Figure S13**). Binding affinity for eIF4E was subsequently determined using a fluorescence polarization assay, demonstrating specificity of binding and maintained affinity (EC_50_ = 37 nM) (**Figure S14A**). Activity of the SmBiT-labeled RNAs was also confirmed using lytic CRISPR RiPCA, which showed specific signal generation with the m^7^GTP-labeled RNA and successful evidence of interaction with LgBiT-eIF4E (**Figure S14B**).

Key to the success of live-cell CRISPR RiPCA is the ability to deliver SmBiT-conjugated RNA into cells. Several commercial transfection reagents were tested for delivery into our LgBiT-eIF4E cells, including TransIT-X2^®^, used in RiPCA 2.0^17,19^, TransIT^®^-mRNA, TransIT-siQUEST^®^, FuGENE^®^, and Lipofectamine^™^ 3000 (**Figure 4A**). Of the tested reagents, Lipofectamine^™^ 3000 proved the most effective (**Figure 4A**), which we hypothesized was due to its superior ability to transfect small RNA species.^39^ As confirmation of our results in CRISPR RiPCA, we performed live-cell confocal microscopy, which demonstrated enhanced cytoplasmic delivery of an AlexaFluor-conjugated m^7^GTP RNA with Lipofectamine^™^ 3000 (**Figure S15**). With TransIT-X2^®^, RNAs appeared to be predominantly trapped in endosomes, which likely explains the lower signal observed in the assay (**Figures 4A** and **S15**).

**Figure 4.**
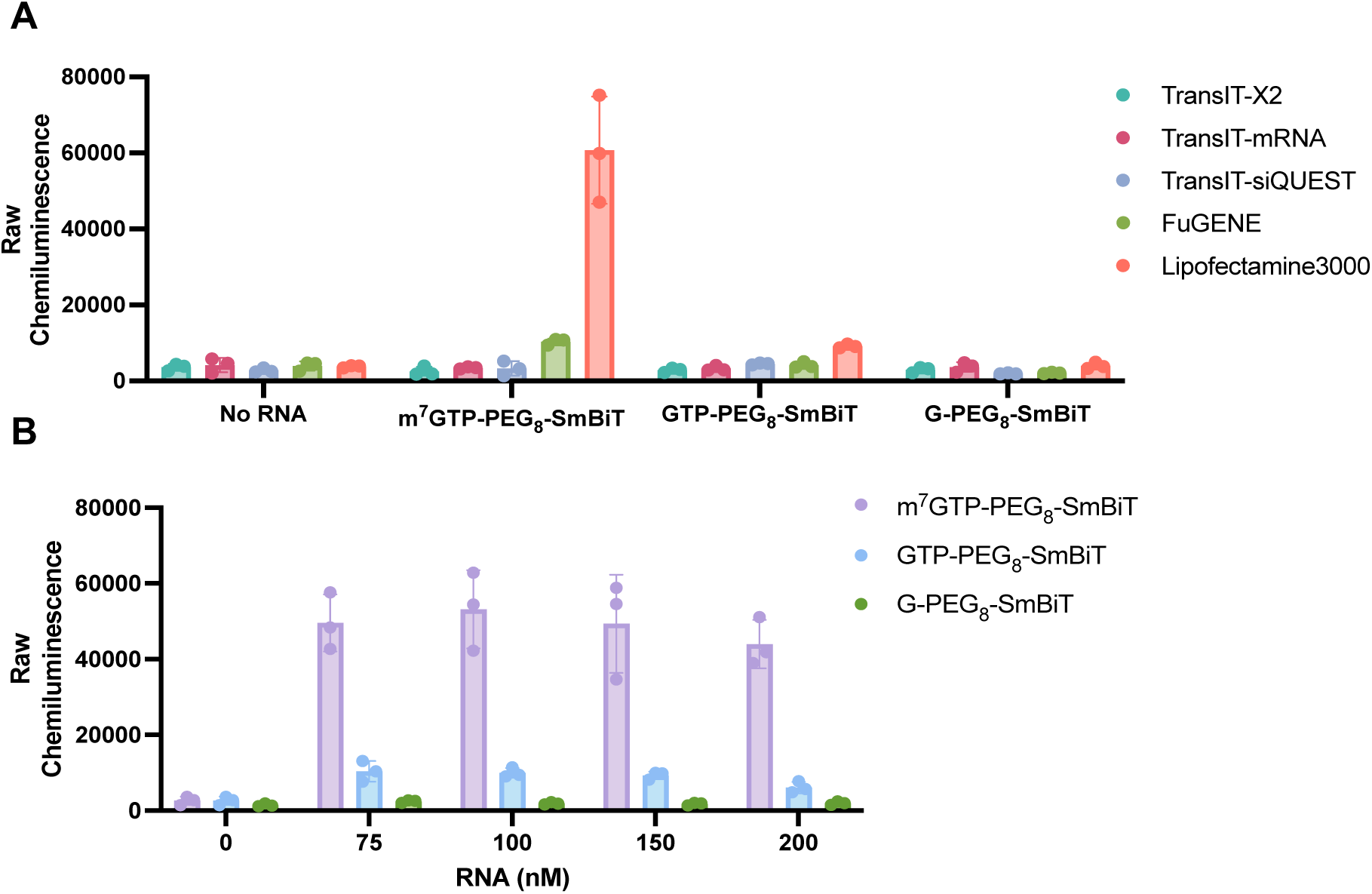
CRISPR RiPCA for eIF4E. (A) Assay signal following delivery of SmBiT-labeled RNA by several commercial transfection reagents. (B) Titration of SmBiT-labeled m^7^GTP RNA and negative control RNAs.

Encouraged by these results, we moved to analyze the specificity and dose-dependence of the assay signal. We performed titrations with both m^7^GTP-SmBiT RNA and negative control SmBiT-labeled RNAs. Excitingly, we observed a strong and dose-dependent increase in chemiluminescence signal for the m^7^GTP-SmBiT RNA and no signal increase for negative control RNAs, an outcome not previously achieved with RiPCA 2.0 (**Figure 4B**). To assess on-mechanism activity, we performed a competition experiment with m^7^GTP RNA lacking SmBiT conjugation, notably observing concentration-dependent decrease in signal in line with SmBiT-m^7^GTP RNA-dependent signal generation (**Figure S16**). In sum, based on these promising results, we were confident of CRISPR RiPCA’s ability to detect eIF4E m^7^GpppX capped mRNA interactions and were eager to determine if we would also be able to use this assay platform for inhibitor testing.

### Application to testing small molecule inhibitors of eIF4E and cap-dependent translation

As the rate-limiting translation initiation factor, eIF4E plays a major role in controlling cap-dependent translation, and many cancers become reprogrammed towards addiction to enhanced eIF4E-driven protein synthesis.^12^ As such, academic and industrial researchers have worked to design small molecule inhibitors of eIF4E to test the therapeutic hypothesis that modulation of eIF4E function would have promising anti-cancer activity.^12^ Based on its known binding pocket for the m^7^GpppX cap, analogues of this modified nucleotide have been designed, including cell-permeable prodrugs by our group.^13,14^ While these compounds have shown promising *in vitro* binding to eIF4E and phenotypic cell activity,^13,14^ confirmation of on-target activity in a live cell-based assay was not possible. With eIF4E CRISPR RiPCA in hand, we sought to use this assay to evaluate the on-target inhibitory activity of a representative from our second-generation cap analogue prodrug series (**Figure 5A**).^14^ As an additional set of compounds to include in this analysis, we synthesized representative eIF4E inhibitors reported from eFFECTOR Therapeutics (eFT) as examples of highly potent non-prodrug-based, cap-competitive inhibitors (**Figure 5A** and **Table S4**).^40^

**Figure 5.**
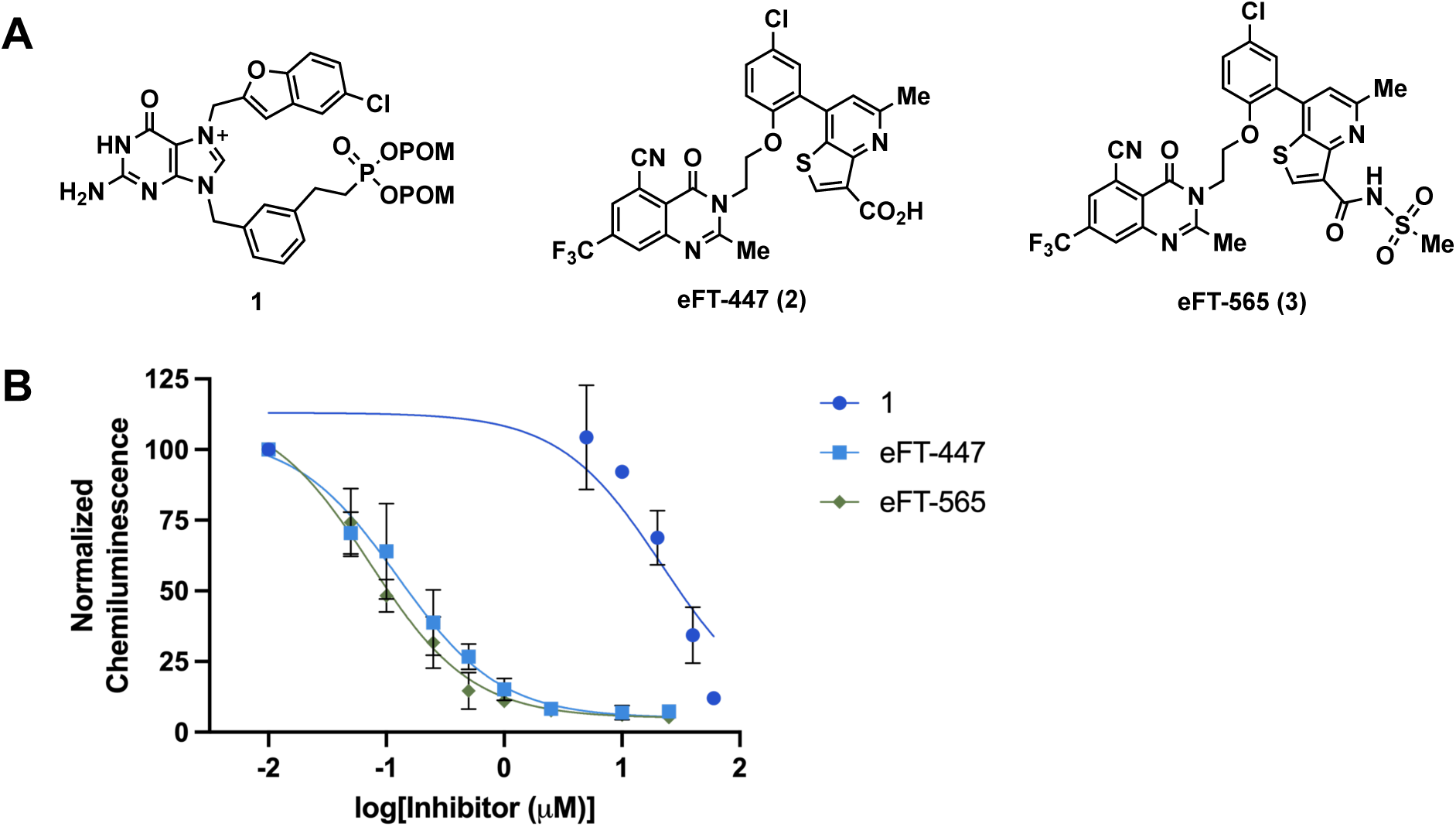
eIF4E CRISPR RiPCA with cap-competitive inhibitors. (A) Structures of compounds tested. (B) CRISPR RiPCA data after 6-hour treatment. IC_50_ values as 95% confidence intervals: **1** (14−36 μM), **2** (91−171 nM), and **3** (57−92 nM).

To assess inhibitory activity, eIF4E CRISPR RiPCA was plated for 18 hours and treated with compounds for 6 hours before reading the assay at 24 hours. In-line with reported binding affinities (**Table S4**), eFT compounds **2** and **3** exhibited nanomolar inhibitory activity in CRISPR RiPCA (IC_50_ values of 124 nM and 73 nM, respectively) (**Figure 5B**). As these analogues differ only by the substitution of the methylthienopyridine ring, we hypothesize that the enhanced activity of compound **3** is due to both improved binding affinity and cellular penetrability by replacing the carboxylic acid with a methylsulfonyl carboxamide group. Prodrug **1**, on the other hand, exhibited reduced activity in comparison (IC_50_ value of 22 μM), but inhibition correlative with previous measures of cellular phenotypic activities for this compound.^14^ Notably, cell death was not observed with any of these compounds at the concentrations tested (**Figure S17**). These results excitingly demonstrate that eIF4E CRISPR RiPCA can be used to assess cap-competitive inhibitors across a range of potencies.

While inhibition of cap binding is the most direct mechanism to inhibit eIF4E’s activity, regulation of eIF4E is complex and mediated through both protein-protein interactions (PPIs) and post-translational modifications (**Figure 6A**).^12^ Importantly, small molecule inhibitors of each of these regulatory mechanisms are known; however, discrepancies in how these molecules affect eIF4E-m^7^GpppX capped mRNA binding have been reported.^41–45^ Thus, we sought to use our assay to shed additional light on how these inhibitors affect eIF4E’s cap-binding activity leading to observed phenotypic decreases in cap-dependent translation inside living cells.

**Figure 6.**
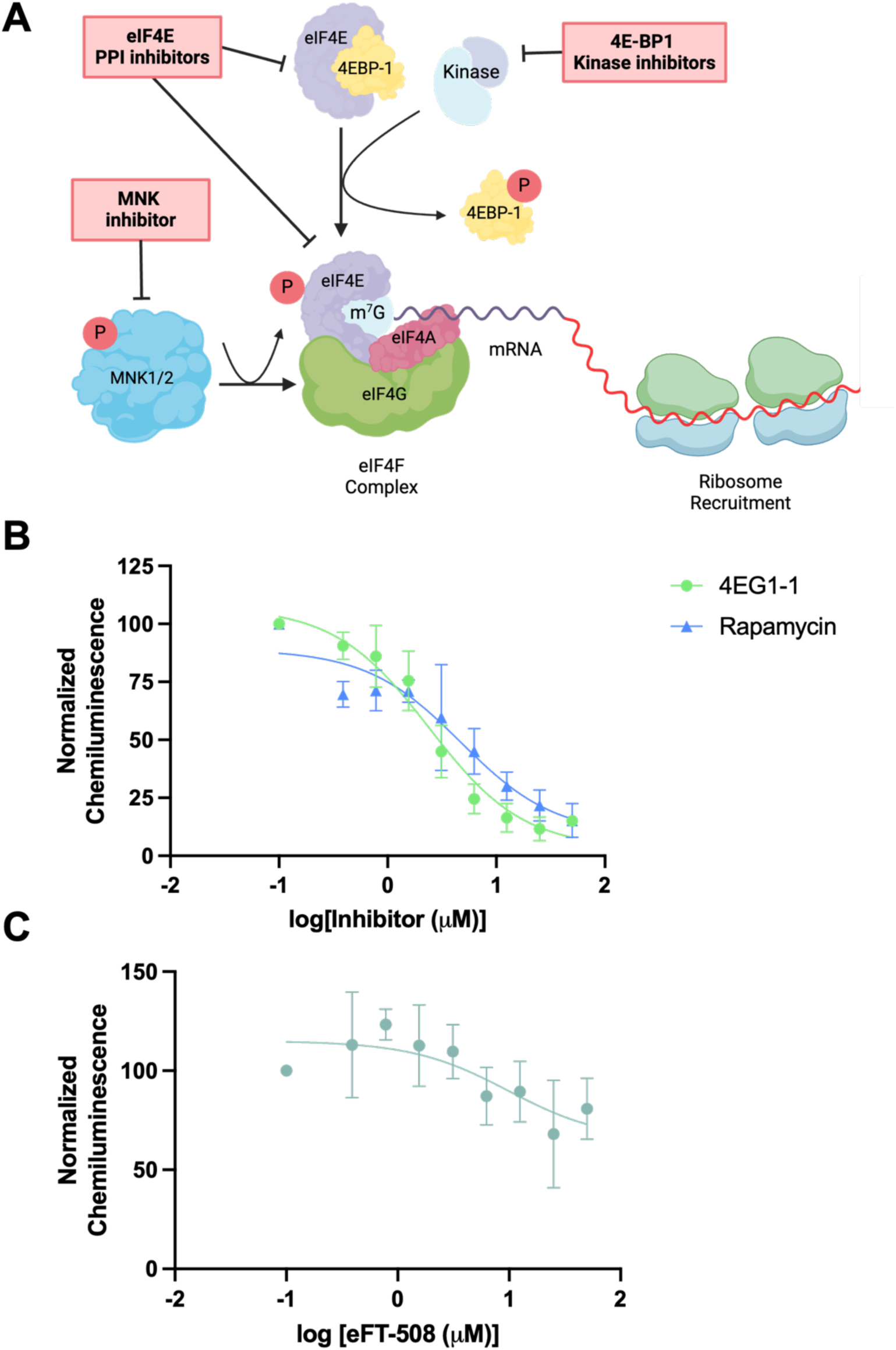
eIF4E CRISPR RiPCA activity by cap-dependent pathway inhibitory compounds. (A) Mechanisms of the regulation of eIF4E and cap-dependent translation initiation. CRISPR RiPCA data after 6-hour treatment with: (B) eIF4E PPI modulators; IC_50_ values as 95% confidence intervals: rapamycin (1.9−11 μM) and 4EGI-1 (1.6−3.6 μM); or (C) inhibitor of eIF4E phosphorylation.

Activation of eIF4E and cap-dependent translation is canonically controlled through regulation of eIF4E PPIs. The primary negative regulator of eIF4E is eIF4E-binding protein 1 (4E-BP1), which sequesters eIF4E from eIF4G, a scaffolding translation initiation factor,^46^ and the eIF4F translation initiation complex by forming an inhibitory PPI.^41,47,48^ This 4E-BP1 activity is canonically regulated by mTORC1-mediated phosphorylation:^49,50^ hypophosphorylated 4E-BP1 binds strongly to eIF4E to inhibit translation, while hyperphosphorylation of 4E-BP1 releases eIF4E to initiate cap-dependent translation.^51–54^ Based on the significance of eIF4E PPIs in mediating translation initiation, reagents have been discovered and developed for their manipulation. Rapamycin is an allosteric inhibitor of mTORC1 which induces 4E-BP1 binding to eIF4E through 4E-BP1 dephosphorylation (**Table S4**).^55^ Small molecule inhibitors of the eIF4E/eIF4G PPI (e.g., 4EGI-1) (**Table S4**),^56,57^ have also been disclosed and found to stimulate 4E-BP1 binding to eIF4E in cells, albeit through an unknown mechanism.

Although enhanced 4E-BP1 binding to eIF4E has not been shown to affect cap binding in biochemical and biophysical assays, recent studies using live-cell fluorescence correlation spectroscopy demonstrated release of the cap from eIF4E following treatment with a mTOR inhibitor.^42^ To better understand how these inhibitors affect binding of eIF4E to m^7^GpppX capped mRNA, LgBiT-eIF4E cells were treated with either rapamycin or 4EGI-1 for 6 h for detection using CRISPR RiPCA. Absence of viability defects was confirmed using the CellTiter-Glo^®^ cell viability assay (**Figure S18**). Excitingly, like previous live-cell studies, with each inhibitor, we observed a dose-dependent decrease in eIF4E binding to our m^7^GTP RNA substrate (IC_50_ values of 4.7 and 2.4 μM for rapamycin and 4EGI-1, respectively) (**Figure 6B**). These results imply that enhanced binding of 4E-BP1 to eIF4E leads to reduced affinity of inhibition of binding to the cap. However, based on the time of our treatment, we cannot rule out the possibility that the decreases in signal observed in CRISPR RiPCA are due to the localization of eIF4E/4E-BP1 complexes to the nucleus.^42,58^ Based on confocal imaging studies, our m^7^GTP RNA substrate does not traverse into the nucleus (**Figure S15**), so we would be unable to detect interactions in this compartment, unlike previous RiPCA systems.^16–19^ Nonetheless, as cap affinity pulldown assays from lysed cells are commonly used to assess inhibitory activity against eIF4E PPIs,^15,56,57,59^ these results raise questions regarding the usage of this assay and call into question the population of eIF4E whose activity is measured in this assay.^60^

In addition to PPIs, eIF4E is also regulated via phosphorylation by MNK1/2 kinases at serine 209.^61^ While the exact function of this post-translational modification is unknown, several studies have indicated that phosphorylation of eIF4E may play a role in initiation factor recycling, affecting the translation of select transcripts, particularly those encoding for survival- and invasion-promoting proteins and cytokines.^43,62–66^ Mechanistically, conflicting reports have found eIF4E phosphorylation to be both inhibitory and stimulatory for cap binding.^43,45^ In addition to the development of cap-competitive eIF4E inhibitors, eFFECTOR Therapeutics also developed the selective MNK1/2 kinase inhibitor tomivosertib (eFT-508).^67^ As we previously showed that LgBiT-eIF4E can be phosphorylated and modulated by this agent (**Figure S8G**), we wanted to further explore how this modulation would affect cap binding in CRISPR RiPCA. Interestingly, treatment with eFT-508 yielded negligible effects on eIF4E’s ability to bind to our m^7^GTP RNA substrate (**Figure 6C**). Although we cannot rule out the possibility that our RNA substrate does not contain elements that are important for inducing functional consequences of eIF4E phosphorylation (e.g., 5′ UTR motifs) or cell type context-dependent effects of eIF4E phosphorylation, inhibition of this post-translational modification does not appear to impact eIF4E binding to capped RNA in HEK293 cells.

### CONCLUSIONS

In conclusion, using the interaction of reader RBP, eIF4E, with m^7^GpppX capped RNA as a model, we demonstrate both challenges and optimizations of our RiPCA live-cell assay technology for detecting and screening RPIs. While RiPCA 2.0 failed for this RPI system, further engineering of our platform using CRISPR/Cas9 gene editing and the development of RNA-SmBiT peptide conjugates led to a physiologically relevant, robust, sensitive, and highly selective assay for monitoring cap binding to eIF4E and inhibition of this disease-relevant RBP. As context-dependent regulation of RPIs is likely for many RBPs, we envision the CRISPR RiPCA platform being enacted on cellular models of disease to enable screening to identify chemical probes and drug-like leads for the therapeutic targeting of RPIs, as well as screening of RNA-binding preferences of RBPs in live cells. Indeed, we envision CRISPR RiPCA as a targeted phenotypic approach, as we have demonstrated how disruption of mechanistically distinct molecular events using diverse small molecule modulators can impact RBP affinity for an RNA substrate. In this regard, screening using CRISPR RiPCA may afford an opportunity to evaluate promising ligandable pathways for manipulating cellular RPIs. Efforts towards these applications will be reported in due course.

## ASSOCIATED CONTENT

## Supporting Information

The Supporting Information is available free of charge at…

Methods and supplemental figures and tables (PDF)

### Accession Codes

eIF4E, P06730

## AUTHOR INFORMATION

### Notes

The authors declare no competing financial interest.

## ACKNOWLEDGMENTS

This work was supported by the NIH (R01 GM135252 and R35 GM153185 to A.L.G.), the Michigan Center for Therapeutic Innovation (A.L.G. and J.C.R.), and the University of Michigan Rackham Merit Fellowship (G.V.H.). We thank the University of Michigan Advanced Genomics Core for whole genome sequencing of LgBiT-eIF4E cells. Figures 1, 2, 3A, 6A, S6A, S7, S8A, S8C, and S9A were created using Biorender.com

## ABBREVIATIONS

eIF4E: Eukaryotic Initiation Factor 4E
RBP: RNA-Binding Protein
RMP: RNA-Modifying Protein
METTL3: Methyltransferase-Like Protein 3
m^6^A: *N^6^*-methyladenosine
m^7^G: *N*^7^-methylguanosine
RPI: RNA-Protein Interaction
PPI: Protein-Protein Interaction
4E-BP1: eIF4E Binding Protein 1
mTOR: Mammalian Target of Rapamycin
VEGF: Vascular Endothelial Growth Factor
MDM2: Murine Double Minute 2
Bcl-2: B-cell leukemia/lymphoma 2
eIF4G: Eukaryotic Initiation Factor 4G
eIF4F: Eukaryotic Initiation Factor 4F
CETSA: Cellular Thermal Shift Assay
RiPCA: RNA-interaction with Protein-mediated Complementation Assay
SmBiT: Small subunit of NanoBiT
LgBiT: Large subunit of NanoBiT
CRISPR: Clustered Regularly Interspaced Short Palindromic Repeats Cas9
CRISPR: associated Protein 9
PEG: Polyethylene Glycol
FP: Fluorescence Polarization
AUF1: AU-rich element RNA-Binding Protein 1
RNP: Ribonucleotide Protein
FACS: Fluorescence-Activated Cell Sorting
GFP: Green Fluorescent Protein
HiBiT: High Affinity binding subunit of NanoBiT
mTORC1: Mammalian Target of Rapamycin Complex 1
MNK1/2: MAPK-interacting Protein Kinase 1 and 2

## Table of contents graphic

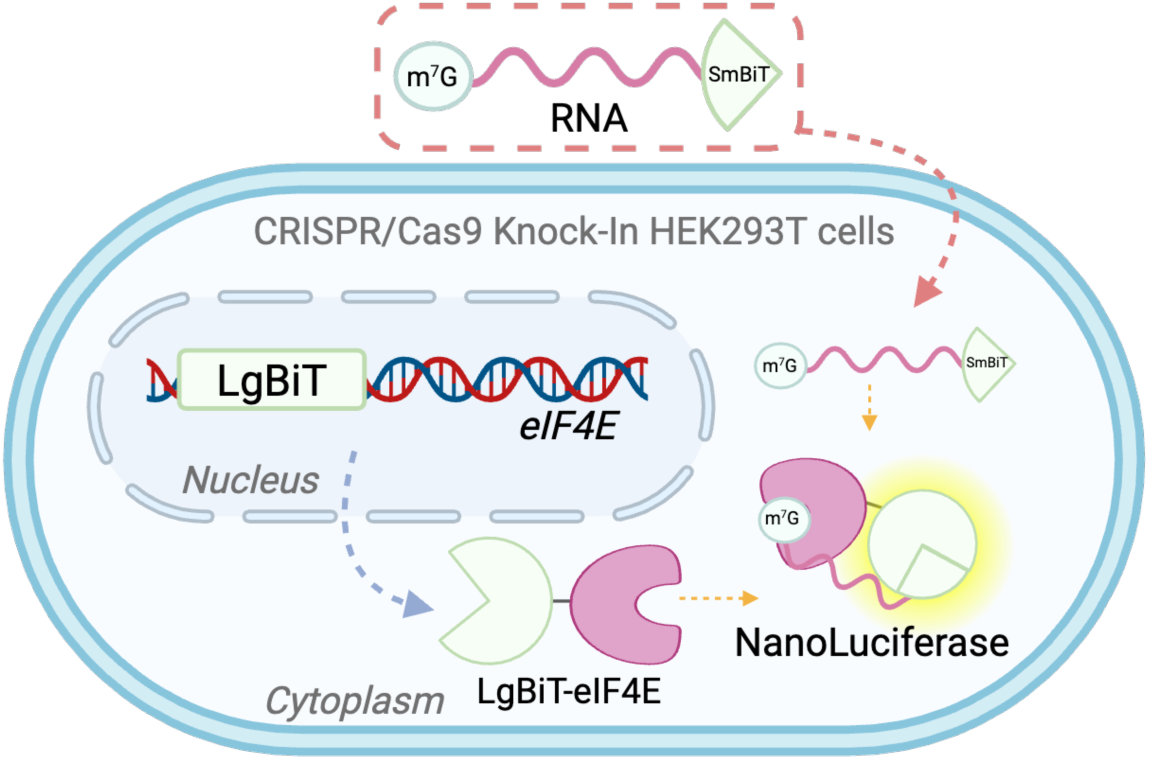

