## Supplementary Information for "CRISPR RiPCA for Investigating eIF4E-m^7^GpppX Capped mRNA Interactions"

| | |
|---|---|
| **A. Materials and Methods** | Pages S2−S8 |
| **B. Supplemental Figures and Tables** | Pages S9−S19 |
| **C. References** | Page S20 |
| **D. Characterization Data for 2 and 3** | Pages S21−S25 |
| **E. Synthesis Details** | Pages S26−S37 |

#### A. MATERIALS AND METHODS

**Materials.** Chemically synthesized m$^7$GTP containing RNA (deprotected, desalted, and HPLC purified) containing a 3′ uridine 7-carbon amine, as well as an unmethylated (GTP) version of this RNA, were purchased from TriLink Biotechnologies and used as received for labeling reactions. The chemically synthesized 10-nucleotide RNA sequence (G) containing a 3′ uridine 6-carbon amine (deprotected, desalted, and HPLC purified) was purchased from Dharmacon and used as received for labeling reactions. HaloTag® Succinimidyl Ester O2 and O4 ligands were purchased from Promega and used as received (Cat. No. 1691 and P6751). HaloTag® Succinimidyl Ester containing a PEG$_6$ linker was synthesized following the previously published protocol.[1] The HaloTag® Succinimidyl Ester Ligands were dissolved in DMSO, and single-use aliquots were stored at -80 °C to avoid degradation. Flp-In™-293 cells and associated vectors were purchased from Thermo Fisher Scientific (Invitrogen Cat. No. 75007 and 601001, respectively). The Nano-Glo® Live Cell Assay System was purchased from Promega and used as received (Cat. No. N2012). The TransIT-X2® transfection reagent was purchased from Mirus (Cat. No. 6000) and used as received. The Lipofectamine™ 300 transfection reagent was purchased from Thermo Fisher Scientific (Cat. No. L30000008) and used as received. Samples for TransIT-mRNA (Cat. No. 2251), and TransIT-siQUEST (Cat. No. 2111) were obtained through Mirus and used as received. The FuGENE SI transfection reagent (Cat. No. E9311) was obtained from Promega and used as received. SmBiT Succinimidyl Ester containing a PEG$_8$ linker was provided by Promega, dissolved in DMSO, and single-use aliquots were stored at -80 °C to avoid degradation.

**General assay and data analysis methods.** Chemiluminescence data was collected on a BioTek Cytation3 plate reader. All data was analyzed using GraphPad Prism version 10.2.3 for macOS. Normalized FP/mP values were obtained by dividing the average of no RNA-containing samples and multiplying by 100. For RiPCA 2.0 normalized chemiluminescence, values were obtained by dividing the average of triplicate wells containing LgBiT-eIF4E DNA only. CRISPR RiPCA normalized chemiluminescence values were obtained by dividing the average signal of DMSO-treated triplicate wells transfected with m$^7$GTP-PEG$_8$-SmBiT RNA and multiplying by 100.

**General cell culture methods.** Flp-In™-293 cells stably expressing SmBiT-HaloTag were grown in DMEM (Corning Cat. No. 10-017-CV) supplemented with 10% FBS (Atlanta Biologicals S11550), L-glutamine (Gibco Cat. No. 25030081), and hygromycin B (100 μg/mL) (Gibco Cat. No. 10687010) at 37 °C with 5% $CO_2$ in a humidifier incubator and passaged at least once before using for an experiment. HEK293T and LgBiT-eIF4E HEK293T cells were grown in DMEM supplemented with 10% FBS and L-glutamine. To passage cells, Trypsin-EDTA (0.25%) (Gibco Cat. No. 25300054) was used and cells were passaged approximately 15–20 times before returning to low passage stocks.

**Molecular cloning.**

*pcDNA3 LgBiT-eIF4E*. eIF4E was amplified from a pFN29K construct containing an *N*-terminal HaloTag fusion protein and inserted into a pcDNA3 vector containing an *N*-terminal LgBiT using standard PCR cloning techniques with XhoI and XbaI restriction enzymes.

Primers:

LgBiT-eIF4E

5′ ATTACTCGAGATGGCGACTGTCGAACCGG
5′ CCTCTAGATTAAACAACAAACCTATTTT

*pcDNA3 eIF4E-LgBiT.* eIF4E was amplified from a pFN29K construct containing an *N*-terminal HaloTag fusion protein and inserted into a pcDNA3 vector containing a *C*-terminal LgBiT using standard PCR cloning techniques with KpnI and AsiSI restriction enzymes. Primers insert a Kozak sequence on the *N*-terminus.

Primers:

eIF4E-LgBiT
5′ AAGGTACCGCCACCATGGCGACTGTCGAACCGG
5′ ATCGGCGATCGCAACAACAAACCTATTTTTAGTG

*peT19bpp LgBiT-eIF4E.* LgBiT-eIF4E was amplified from a pcDNA3 construct and inserted into a peT19bpp vector containing a $His_{10}$ tag using standard cloning techniques with NdeI and BamHI restriction enzymes. Primers insert a Kozak sequence on the *N*-terminus.

Primers:

LgBiT-eIF4E
5′ ATATCATATGGCCACCATGGTCTTCACACTC
5′ GACGGATCCATTAAACAACAAACCTATTTTTAGTGGTGG

**Bioconjugation methods.** Amine-modified RNA substrate (1.0 mM in 100 mM Phosphate buffer, pH 8.0) was mixed with an equivalent volume of HaloTag® ligand (10 mM in DMSO for HaloTag® Succinimidyl Ester O2 and O4, and 20 mM in DMSO for HaloTag Succinimidyl Ester O6) or the SmBiT Succinimidyl Ester reagent (10 mM in DMSO) (**Figure S1**). The reaction was incubated at 25 °C for 2 h before precipitating by adding 0.11× volume of 3.0 M sodium acetate (pH 5.2) and 4 volume equivalents of cold ethanol. The RNA was pelleted at 15,000 rpm for 45 min at 4 °C. The pellet was then re-suspended in 100 mM Phosphate buffer (pH 8.0) at a concentration of 1.0 mM and stored at -80 °C. For RiPCA, the RNAs were diluted to 25 μM for the working concentration in 1.5 mL LoBind® microcentrifuge tubes. The concentration of each RNA was verified using a nanodrop.

**Protein Expression and Purification.**
*eIF4E.* pET19bpp-eIF4E plasmid was transformed into BL21(DE3) cells. Cells were grown at 37 °C to an $OD_{600}$ of 0.6–0.8, induced with 1 mM IPTG, and grown for 16 h at 18 °C. The cells were pelleted and resuspended in lysis buffer (50 mM Tris-HCl, pH 8, 500 mM NaCl, 50 mM imidazole, 20 mM β-mercaptoethanol, 5 mM DTT, and 1% Tween-20 with protease inhibitors) and lysed by sonication. The cell lysate was ultracentrifuged at 15,000 rpm for 2 h and the supernatant was incubated with Ni-NTA resin for 1 h at 4 °C. The resin was washed 3× with wash buffer (50 mM Tris-HCl, pH 8, 500 mM NaCl, 25 mM imidazole, and 5 mM DTT) with 15 min incubations. eIF4E was eluted 2× with 10 mL of elution buffer (10 mM Tris-HCl pH 8, 500 mM NaCl, 100 mM imidazole, and 5 mM DTT). Protein was dialyzed overnight at 4 °C in dialysis buffer (20 mM Tris-HCl pH 7.4, 100 mM NaCl, 2 mM DTT). After dialysis, the protein was concentrated and purified using an FPLC Superdex 75 16/10 column. Pure protein was quantified via the Bradford assay, aliquoted, and stored at -80 °C.

*LgBiT-eIF4E.* pET19bpp-LgBiT-eIF4E plasmid was transformed into BL21(DE3) cells. Cells were grown at 37 °C to an $OD_{600}$ of 0.6–0.8, induced with 1 mM IPTG, and grown for 16 h at 18 °C. The cells were pelleted and resuspended in lysis buffer (50 mM Tris-HCl, pH 8, 500 mM NaCl, 50 mM imidazole, 20 mM β-mercaptoethanol, 5 mM DTT, and 1% Tween-20 with protease inhibitors) and lysed by sonication. The cell lysate was ultracentrifuged at 15,000 rpm for 2 h and the supernatant was incubated with Ni-NTA resin for 1 h at 4 °C. The resin was washed 3× with wash buffer (50 mM Tris-HCl, pH 8, 500 mM NaCl, 25 mM imidazole, and 5 mM DTT) with 15 min incubations. LgBiT-eIF4E was eluted 2× with 10 mL of elution buffer (10 mM Tris-HCl pH 8, 500 mM NaCl, 100 mM imidazole, and 5 mM DTT). Protein was dialyzed overnight at 4 °C in dialysis buffer (20 mM Tris-HCl pH 7.4, 100 mM NaCl, 2 mM DTT). After dialysis, the protein was quantified via the Bradford assay, aliquoted, and stored at -80 °C.

*SmBiT-HaloTag.* pET28a-SmBiT-HaloTag plasmid was transformed into BL21(DE3) cells. The cells were grown at 37 °C to an $OD_{600}$ of 0.6 and induced with 1 mM IPTG overnight at 37 °C. Cells were pelleted and resuspended in lysis buffer (10 mM Phosphate pH 8, 20 mM imidazole, 3 mM KCl, 150 mM NaCl, 0.1% PMSF, and 1 mM DTT) and lysed by sonication. Lysates were then pelleted at 4,000 rpm for 1 h. The supernatant was incubated with Ni-NTA resin, and the resin was washed 3× with wash buffer (10 mM Tris, pH 8, 50 mM imidazole, 500 mM NaCl, 0.1% PMSF, and 1 mM DTT) with 15 min incubations. SmBiT-HaloTag was eluted from the resin with elution buffer (10 mM Tris, pH 8, 500 mM imidazole, 500 mM NaCl, 0.1% PMSF, and 1 mM DTT) and dialyzed overnight into 20 mM Tris pH 7.8, 100 mM KCl, 0.2 mM EDTA, and 10% glycerol. Protein concentration was measured via the Bradford assay, aliquoted, and stored at -80 °C.

**Fluorescence Polarization Assay.** The protocol was carried out as previously described.[2] Briefly, $EC_{50}$ values were determined by incubating increasing concentration of $His_{10}$-eIF4E (0–10 μM) with 15 nM EDA-$m^7$GTP-5-FAM (Jena Bioscience Cat. No. NU-824-5FM) in FP assay buffer (50 mM HEPES pH 7.2, 100 mM KCl, 0.5 mM EDTA, 1 mM DTT, 0.0025% Tween-20, and 0.05 mg/mL BSA). For competition experiments, 15 nM probe and 100 nM $His_{10}$-eIF4E were incubated with increasing concentrations of RNA (0–800 nM). Samples were gently mixed and incubated at room temperature, shaking at 150 rpm for 40 min. FP measurements were collected with 485 excitation and 528 emission polarization filters.

**SmBiT-HaloTag RNA Labeling**. Purified SmBiT-HaloTag (5 μM) and $m^7$GTP-$PEG_2$-Cl RNA (0–5 μM) were incubated in 20 μL of 100 mM Phosphate buffer (pH 8.0) for 1 hour at 25 °C. The reaction was quenched with 4 μL of 6× Laemmli SDS dye (375mM Tris-HCl/Tris Base, 9% SDS (w/v), 50% Glycerol (v/v), 9% 2-mercaptoethanol (v/v), 0.075% Bromophenol blue (w/v)) and run on a casted 10% Next Gel (Cat. No. M256) for 60 min at 150 V. The gel was stained with Coomassie Brilliant Blue, destained with destaining solution ($H_2O$, methanol, and acetic acid in a ratio of 50/40/10 (v/v/v)) before imaging on a Protein Simple gel imager using the Coomassie Brilliant Blue setting.

**Nano-RaPID, a biochemical version of RiPCA.** In a microcentrifuge tube, LgBiT-eIF4E, SmBiT-HaloTag, and HaloTag ligand labeled-RNA were added to assay buffer (50 mM Tris-HCl pH 7.4, 150 mM NaCl, 5% glycerol, 10 mM BME, 0.05% Tween-20, and freshly added 2 mM

DTT) to 500 nM final concentrations in 50 µL. Samples were gently mixed and incubated at room temperature for 30 min. Nano-Glo® luciferase substrate (Cat. No. N1110) was prepared according to the manufacturer's recommendation and 25 µL was added to each reaction tube. Immediately after, the contents of one tube were added into 4 wells of a white, flat-bottom 384-well assay plate (Cat. No. 3572) (15 µL per well). Chemiluminescence data was collected immediately on a BioTek Cytation3 plate reader.

**RiPCA 2.0.** The RiPCA 2.0 protocol using the TransIT-X2® transfection reagent was performed as previously reported.[3] Briefly, room-temperature Opti-MEM™ was added to a 0.5 mL microcentrifuge tube followed by the DNA, RNA, and Transit-X2 (in this order). Each tube was gently vortexed, centrifuged, and incubated for 20 min at room temperature while Flp-In HEK293 cells were harvested. A solution of 200,000 cells/mL was prepared and 300 µL of this solution was added to each microcentrifuge tube containing the transfection solution to be assayed and mixed by pipetting. Using a multichannel pipette, 100 µL per well (×3) of each condition was plated in a white-bottom, tissue culture-treated 96-well plate (Corning Cat. No. 3917). The plate was incubated in a humidified incubator (37 °C and 5% $CO_2$) for 24 h. After incubation, the media was removed and replaced with 100 µL of room temperature phenol-red free Opti-MEM™ (Cat. No. 11058021) and treated with 25 µL NanoGlo® Live Cell Reagent diluted 1:20 according to the manufacturer's recommendation. Chemiluminescence data was collected immediately on a BioTek Cytation3 plate reader.

**Western Blot.** HEK293T cells, Flp-In HEK293 cells or CRISPR-engineered LgBiT-eIF4E HEK293T cells were grown to 50% confluency in a clear bottom 6-well tissue culture-treated (Cat. No. CC7682-7506). For test expression experiments, cells were transfected with 1 µg of pcDNA3 plasmid in 200 µL of Opti-MEM and mixed with 3 µL of PEI transfection reagent (from 1 mg/mL stock). Transfection complexes were gently vortexed and incubated at 25 °C for 15 min before being added to cells in a drop-wise manner. After 24 h, cells were harvested with 250 µL of RIPA (10 mM Tris-HCl, 150 mM NaCl, 1% Triton, 1% sodium deoxycholate, 0.1% SDS, pH 7.2) containing protease inhibitors. Total protein was quantified using BCA (Cat. No. 23225), and 8 µg of total protein was loaded onto a 4–12% Tris-Glycine gel run at 135 V for 90 min. The gel was transferred to a PVDF membrane in Towbin's buffer (25 mM Tris pH 8.3, 192 mM glycine, 20% v/v methanol). The membrane was blocked in 5% non-fat milk for 1 h at 25 °C before adding the primary antibody overnight at 4 °C. For secondary antibodies, HRP-linked anti-mouse IgG (Cat. No. 7076S) and HRP-linked anti-rabbit IgG (Cat. No. 7074S) were used. For our loading control, actin-HRP (Cat. No. sc-47778) was used.

**CRISPR/Cas9-mediated LgBiT incorporation into eIF4E.** HEK293T cells expressing eIF4E *N*-terminally tagged with LgBiT were generated by CRISPR/Cas9 mediated homology-directed repair. Briefly, a 20-nt sgRNA sequence targeting exon 1 of human eIF4E was designed with the target PAM site upstream of the start codon. gRNA was assembled by incubating 1.0 nmol of Alt-R CRISPR RNA (crRNA) with 1.0 nmol of Alt-R trans-activating crRNA (tracrRNA) in 50 µL of Nuclease-Free Duplex Buffer (Integrated DNA Technologies, IDT) at 95 °C for 5 min and then cooled to room temperature. For LgBiT insertion, a donor template was designed using the Benchling CRISPR tool, purchased from IDT, and resuspended to 500 ng/µL in nuclease-free water. RNP complexes were assembled with Cas9-GFP nuclease (Alt-R™ S.p. Cas9-GFP V3 Cat.

No. 10008100), and cells were electroporated using the Ingenio EZporator Electroporation System (Cat. No. 51000). 72 h post-electroporation, GFP-containing cells were isolated by fluorescence-associated cell sorting (FACS). For viable cell sorting, cells were also treated with a DAPI marker (Cat. No. PI62247), and single cells were sorted into clear round-bottom tissue culture-treated 96-well plates (Cat. No. 3799). Following 2 weeks, wells containing clonal cell colonies were passaged into clear-bottom tissue culture-treated 24-well plates (Cat. No. CC7682-7524) to allow further growth of each colony and later passaged into clear tissue culture-treated 6-well plates to provide enough cells for LgBiT lytic detection. Positive colonies were cultured further, and genomic DNA was isolated for junction PCR amplification of the 5′-region on eIF4E. Clones were additionally analyzed by Western blot, using anti-LgBiT (Promega Cat. No. N7100) and anti-eIF4E (Cell Signaling Cat. No. 9742) antibodies.

**LgBiT Lytic Detection.** LgBiT-edited cells were resuspended to 20,000 cells in 100 μL of growth medium, plated in a white-bottom tissue culture-treated 96-well plate, and cultured for 24 h before treatment. After incubation, the media was removed and replaced with 100 μL of room temperature phenol-red free Opti-MEM™ and treated with an equal volume of NanoGlo HiBiT Lytic Reagent (Promega Cat. No. N3030), consisting of Nano-Glo Lytic Buffer, NanoGlo HiBiT Lytic Substrate, and HiBiT Control Protein (Promega Cat. No. N3010), added according to the manufacturer's recommendation. Cells were incubated by shaking for 10 min at room temperature. Chemiluminescence data was collected immediately on a BioTek Cytation3 plate reader.

**Junction PCR.** Primers were designed to verify LgBiT integration at the correct locus and test for homozygosity. LgBiT-eIF4E HEK293T cells were grown in 35 mm tissue culture-treated dishes (Cat. No. 430165) until 80% confluency, and genomic DNA was extracted using the Qiagen Blood and Cell Culture DNA Mini Kit (Cat. No. 13323).

*Primers for junction site:*

(Outside left homology arm) FWD 5′ GGTGGGGGAGAGACTCCACTTCCCA
(Within LgBiT sequence) REV 5′ GCGCTCGTCGATAATTTTGTTGCCG

*Test for all alleles:*

(Outside left homology arm) FWD 5′ GGTGGGGGAGAGACTCCACTTCCCA
(Outside right homology arm) REV 5′ CCACCCTGTAGACACCTCGCGGTTC

*Full-length site:*

(With left homology arm) FWD 5′ CACGTGGCCAGAAGCTGGCCAATCC
(With right homology arm) REV 5′ AGTCCCCCAGTCAGAAGGAAGACGG

**Confocal Microscopy.**

*Protocol for fixed cells***.** LgBiT-eIF4E HEK293T cells were grown in 12-well plates on poly-L-lysine-coated coverslips. Cells were washed twice with 1X PBS and then fixed with 4% paraformaldehyde for 10 min at room temperature. A 0.1% Triton X-100 diluted in PBS was added to cells and incubated for 10 min at room temperature to permeabilize cells. Cells were blocked for 1 h at room temperature with 3% BSA, and 0.1% Triton X-100 in PBS. Primary antibodies for eIF4E (Cell Signaling Cat. No. 9742) and LgBiT (Promega Cat. No. N7100) were diluted to a final concentration of 2 μg/mL in 1% BSA in PBS and incubated for 2 h at room temperature. Cells were washed twice with 1% BSA in PBS with 10-min incubations at room temperature. Secondary antibodies were diluted to a final 4 μg/mL in 1% BSA in PBS and incubated for 1 h at room temperature, protected from light. Cells were washed twice with 1% BSA in PBS with 10-min incubations at room temperature. Coverslips were mounted to glass slides by adding one drop of the DAPI-mounting solution (Cat. No. P36962) and allowed to dry 24 h before imaging. Images were taken at 60× (with oil) using a Nikon W1-SoRa confocal microscope. Images were processed with NIS-Elements.

*Protocol for Live-cell Imaging.* AlexaFluor488-labeled RNAs were diluted in 50 μL of Opti-MEM™ for reverse transfection using TransIT-X2® and Lipofectamine™ 3000 (Thermo Scientific), following the manufacturer's protocol. While transfection complexes were incubated for 20 min at room temperature, HEK293T cells were diluted to a density of 200,000 cells/mL. For each condition, 200 μL of the cell solution was mixed before plating in an 8-well chambered coverglass (Nunc™ Lab-Tek™ II). The chambered coverglass was incubated in a tissue culture incubator (37 °C and 5% $CO_2$) for 24 h. For nuclear staining, the media was replaced with 2.5 μg/mL Hoescht33342 (Fisher) diluted in Phenol-Red Free DMEM (Supplemented with 10% FBS + L-Glutamine) and returned to the incubator for 30 min. The media was removed and replaced with Phenol-Red Free DMEM media. Fluorescence was visualized at 60× (with oil) using a Nikon W1-SoRa confocal microscope. Images were processed with NIS-Elements.

**Whole Genome Sequencing.** LgBiT-eIF4E HEK293T cells were grown in 35 mm tissue culture-treated dishes (Cat. No. 430165) until 80% confluency, and genomic DNA was extracted using the Qiagen Blood and Cell Culture DNA Mini Kit (Cat. No. 13323). Library prep and next-generation sequencing were carried out in the Advanced Genomics Core at the University of Michigan (UM-AGC). 151bp paired-end sequencing according to the manufacturer's protocol (Illumina NovaSeqXPlus). BCL Convert Conversion Software v4.0 (Illumina) was used to generate de-multiplexed Fastq files.

**WGS data preprocessing, variant calling and visualization.** Fastqs from the UM-AGC were trimmed with bbduk and aligned with BWA-MEM against hg38.[4,5] Base quality score recalibration was done using GATK.[6] Quality score recalibrated BAM files were then ran through delly, manta, and SvABA to generate candidate variants.[7–9] A consensus call was conducted by running the output vcf files through survivor with n-1 threshold.[10] Data QC was performed with fastqc pre- and post- trimming.[11] Alignment QC was done with samtools stats. Coverage following trimming and QC was 33x. gRNA off-targets were deduced using Cas-OFFinder and distance calculations were performed using GenomicRanges.[12,13] Plotting was done with ggplot2.[14]

**Visualization of LgBiT-eIF4E Insertion.** WGS data was prepared as described above up to alignment. During alignment, a custom hg38 genome was used with an LgBiT-eIF4E contig comprised of the LgBiT sequence as well as the 500 nts 5`and 3' of the LgBiT sequence. The resulting BAM file was visualized using IGV.[15]

**Data Availability.** Raw fastqs of the LgBiT-eIF4E HEK293T cells WGS have been deposited into the NCBI's sequence read archive (SRA) and are accessible at BioProject accession, PRJNA1276515. Variant calls and scripts used in analysis have been archived on Zenodo, 10.5281/zenodo.15658775.

**Cap pull-down**. The cap pull-down assay was done as previously described.[16,17] Briefly, LgBiT-eIF4E HEK293T cells were grown in 6 cm tissue culture-treated dishes until 80% confluency.

Cells were then lysed in cap pull-down buffer (50 mM HEPES-KOH pH 7.5, 150 mM KCl, 1 mM EDTA, 2 mM DTT, and 0.1% Tween-20) containing protease inhibitors. The cell lysate was centrifuged at 15,000 rpm for 25 minutes. The supernatant was subsequently incubated for 2 hours at 4 °C with $m^7$GDP-agarose resin. Beads were washed 3× with the cap pull-down buffer, 1× with 1× TBS, and 1× with water. Proteins were eluted by boiling in 2× LDS sample buffer for 10 min at 70 °C, resolved on a 4-12% Bis-Tris gel, and Western Blot protocol was carried out as previously described. Primary antibodies eIF4E (Cell Signaling Cat. No. 9742), 4E-BP1 (Cell Signaling Cat. No. 9644), and eIF4G (Cell Signaling Cat. No. 2498S) were used.

**MNK Inhibitor Treatment.** LgBiT-eIF4E HEK293T cells were grown to 50% confluency in a clear bottom 6-well tissue culture-treated (Cat. No. CC7682-7506). After 18-24 h, cells were treated with 0.5 and 1 μM eFT-508 (Cat. No.) and DMSO control. After 6 h, cells were harvested in 250 μL of RIPA buffer (10 mM Tris-HCl, 150 mM NaCl, 1% Triton, 1% sodium deoxycholate, 0.1% SDS, pH 7.2) containing protease and phosphatase inhibitors. Total protein was quantified using BCA (Cat. No. 23225) and analyzed by Western blot. Primary antibody Phospho-eIF4E (Ser209) (Cell Signaling Cat. No. 9741S) was used. For our loading control, actin-HRP (Cat. No. sc-47778) was used.

**RNA Gel for SmBiT-RNA labeling efficiency.** RNA samples were quantified and prepared at 500 nM with 2× Novex TBE-Urea Sample Buffer (Invitrogen, Cat. No. LC6876) and run on a Novex 15% TBE-Urea gel (Invitrogen, Cat. No. EC68855BOX) at 180 V for 60 min. The gel was incubated in SYBR gold nucleic acid stain (Invitrogen, Cat. No. S11494), incubated shaking for 5 min at 25 °C, and imaged on a BioRad ChemiDoc Gel Imager.

**CRISPR RiPCA Protocol.** For each condition, 37.5 μL of room-temperature Opti-MEM™ was added to a 0.5-mL microcentrifuge tube, followed by 100 nM of SmBiT-labeled RNA and 1 μL of Lipofectamine™ 3000 (in this order). Each tube was gently vortexed, centrifuged, and incubated for at least 15 min at room temperature while LgBiT-eIF4E HEK293T cells were harvested. A solution of 200,000 cells/mL was prepared, and 300 μL of this solution was added to each microcentrifuge tube containing the transfection solution to be assayed and mixed by pipetting. Using a multichannel pipette, 100 μL per well (×3) of each condition was plated in a white-bottom, tissue culture-treated 96-well plate (Corning Cat. No. 3917). The plate was incubated in a humidified incubator (37 °C and 5% $CO_2$) for 24 h. After incubation, the media was removed and replaced with 100 μL of room temperature phenol-red free Opti-MEM™ (Cat. No. 11058021) and treated with 25 μL NanoGlo® Live Cell Reagent diluted 1:20 according to the manufacturer's recommendation. Chemiluminescence data was collected immediately on a BioTek Cytation3 plate reader. For unlabeled RNA competition experiments, the CRISPR RiPCA protocol was followed with the following change. In addition to the SmBiT-labeled RNA added, varying amounts of unlabeled RNA were added to each transfection solution. The transfection solution was scaled 10-fold for inhibitor testing, and 100 nM SmBiT-labeled RNA was added to 375 μL of Opti-MEM™, followed by 10 μL of Lipofectamine™ 3000. After 15 min, the transfection complex was added to 3 mL of 200,000 cells/mL. The solution was mixed, and 100 μL were distributed across 30 wells. The CRISPR RiPCA experiment was plated and, at 18 h, 10 μL of inhibitor in a 5% DMSO intermediate solution was added for a final 0.5% DMSO concentration for each respective concentration, and the assay was read 6 h after inhibitor treatment (24 h total time) as described above.

**Lytic CRISPR RiPCA SmBiT-HaloTag-RNA and SmBiT-RNA Protocol**. LgBiT-eIF4E HEK293T cells were seeded by dispensing 10,000 cells per well in 50 µL in a TC-treated white 384-well plate and cultured 24 h before assay. For inhibitor treatment, 5% DMSO mid-stock compound solutions were prepared in media, and cells were treated in triplicates with 5 µL for a final 0.5% DMSO concentration. After 24 h of treatment, the cell plate was removed from the incubator, and media contents were removed from the cells by dumping them into the sink. The cell plate was gently smacked against a sterile cushioned surface to remove all media from the wells. $K_D$ values were determined by incubating increasing concentrations of $m^7G$-$PEG_8$-SmBiT, GpppG-$PEG_8$-SmBiT, and G-$PEG_8$-SmBiT (0-500 nM) in lysis buffer (50 mM HEPES pH 7.5, 150 mM NaCl, 1 mM DTT, 1 mg/mL BSA, and 0.02% Digitonin). 25 µL of the lysis solution was added in triplicates for each condition. The plate was centrifuged at 800 rpm for 30 s before shaking at 400 rpm for 45 min. The Nano-Glo® Luciferase substrate was prepared according to the manufacturer's protocol, and 6.2 µL per well was added. The plate was centrifuged at 800 rpm for 30 s before shaking at 400 rpm for 2 min. Chemiluminescence data was collected immediately on a BioTek Cytation3 plate reader.

#### B. Supplemental Figures

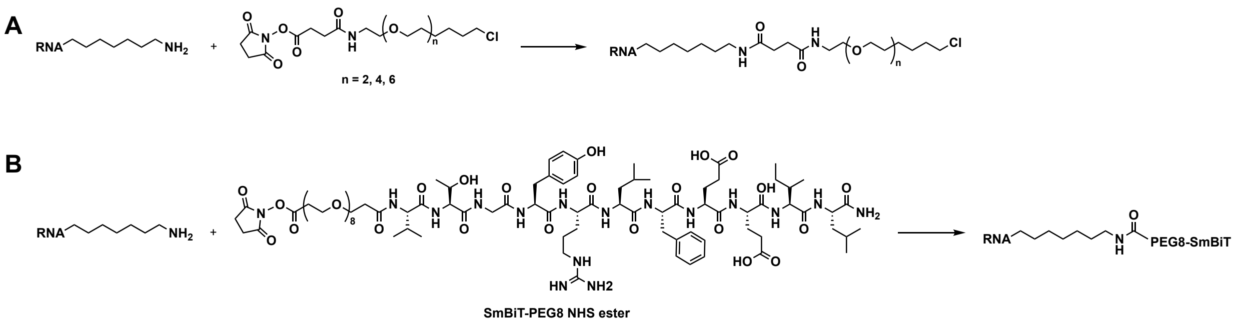


**Figure S1.** RNA bioconjugation reactions. Chemically synthesized amine-containing RNAs were reacted with (A) succinimidyl ester-based HaloTag ligands containing 2, 4, or 6 PEG linker lengths, or (B) a PEG8-labeled SmBiT peptide succinimidyl ester.

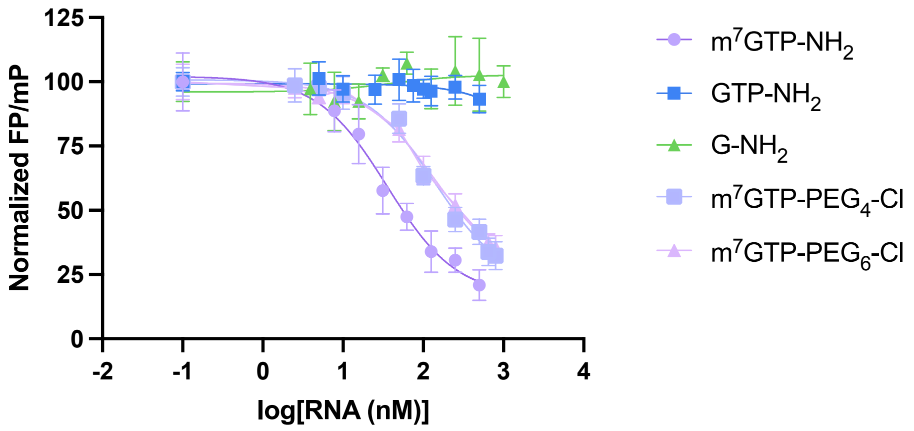


**Figure S2.** Fluorescence polarization assay to measure binding affinity of conjugated RNAs for eIF4E.

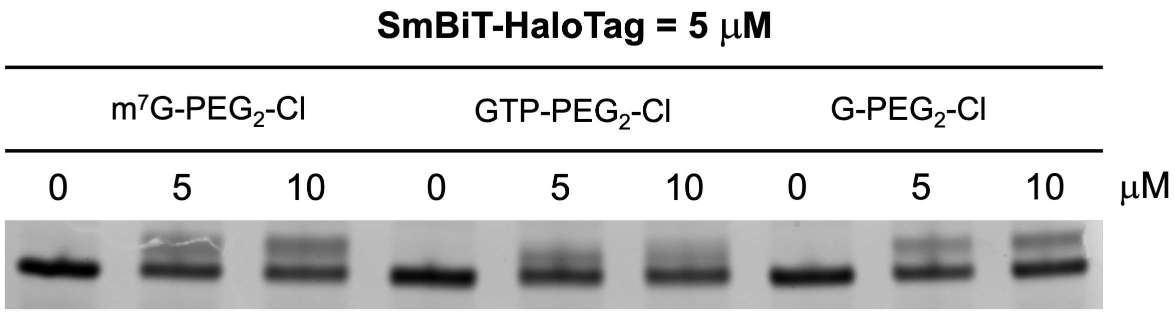


**Figure S3.** EMSA to characterize labeling of RNA substrates with SmBiT-HaloTag protein. Labeling is indicated by the top band shifted to higher molecular weight.

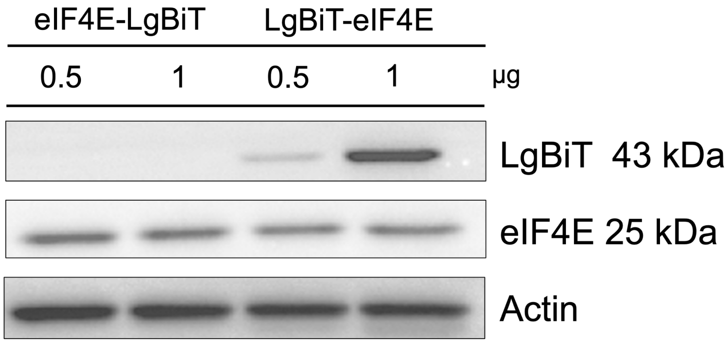


**Figure S4.** Test expression of LgBiT-tagged eIF4E constructs in Flp-In HEK293 cells stably expressing cytoplasmic SmBiT-HaloTag.

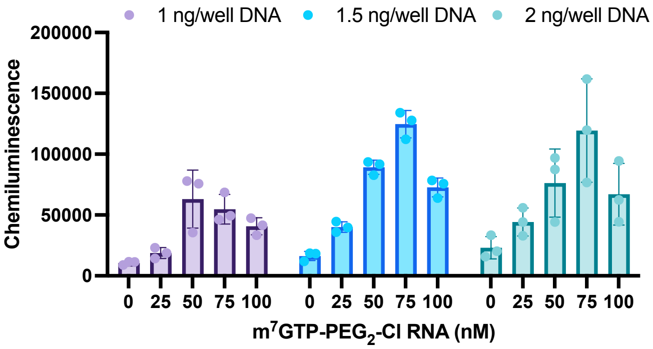


**Figure S5.** Titration of DNA (LgBiT-eIF4E) and RNA concentrations and their impact on RiPCA 2.0 signal.

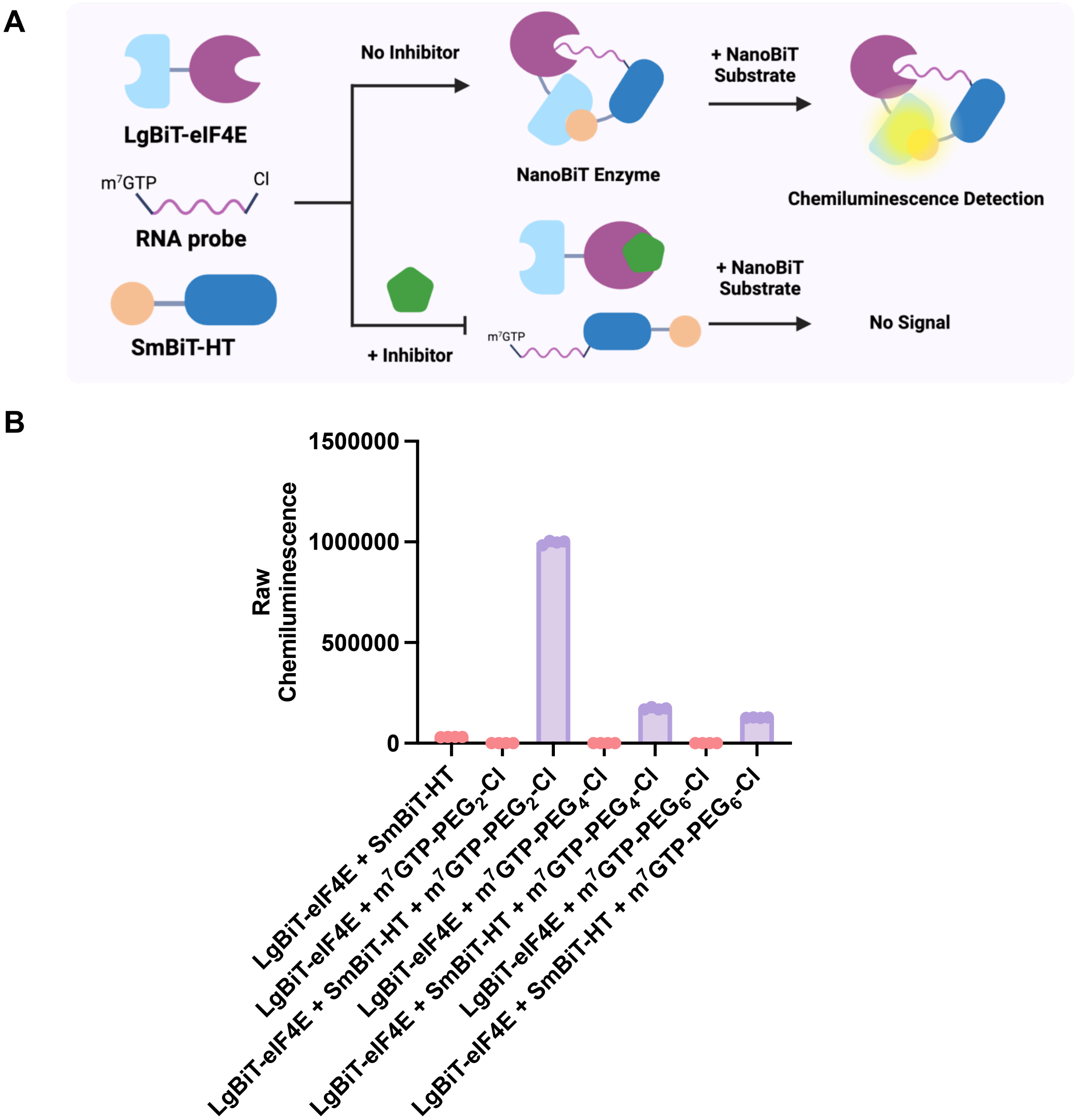


**Figure S6.** Nano-RaPID for ternary complex validation and inhibitor activity. (A) Scheme of Nano-RaPID, a biochemical version of RiPCA utilizing purified proteins. (B) Preferential ternary complex formation with shorter HaloTag ligand PEG linker length m$^7$G RNA.

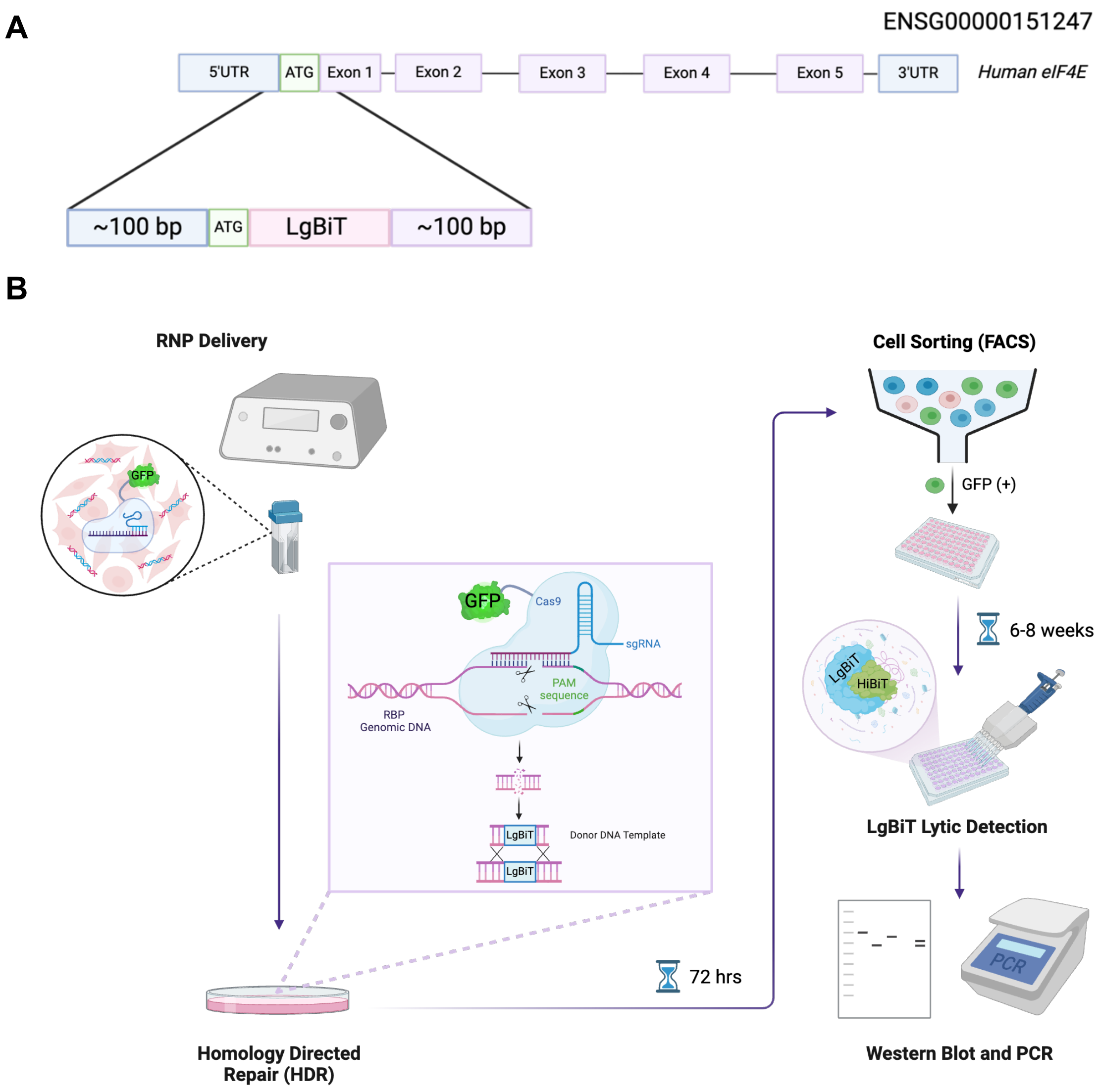


**Figure S7.** CRISPR/Cas9 HDR-mediated LgBiT Knock-In Cell Line. (A) *N*-terminal incorporation of LgBiT onto human eIF4E (Ensembl gene: ENSG00000151247). (B) Scheme for generation of LgBiT knock-in HEK293T cell line.

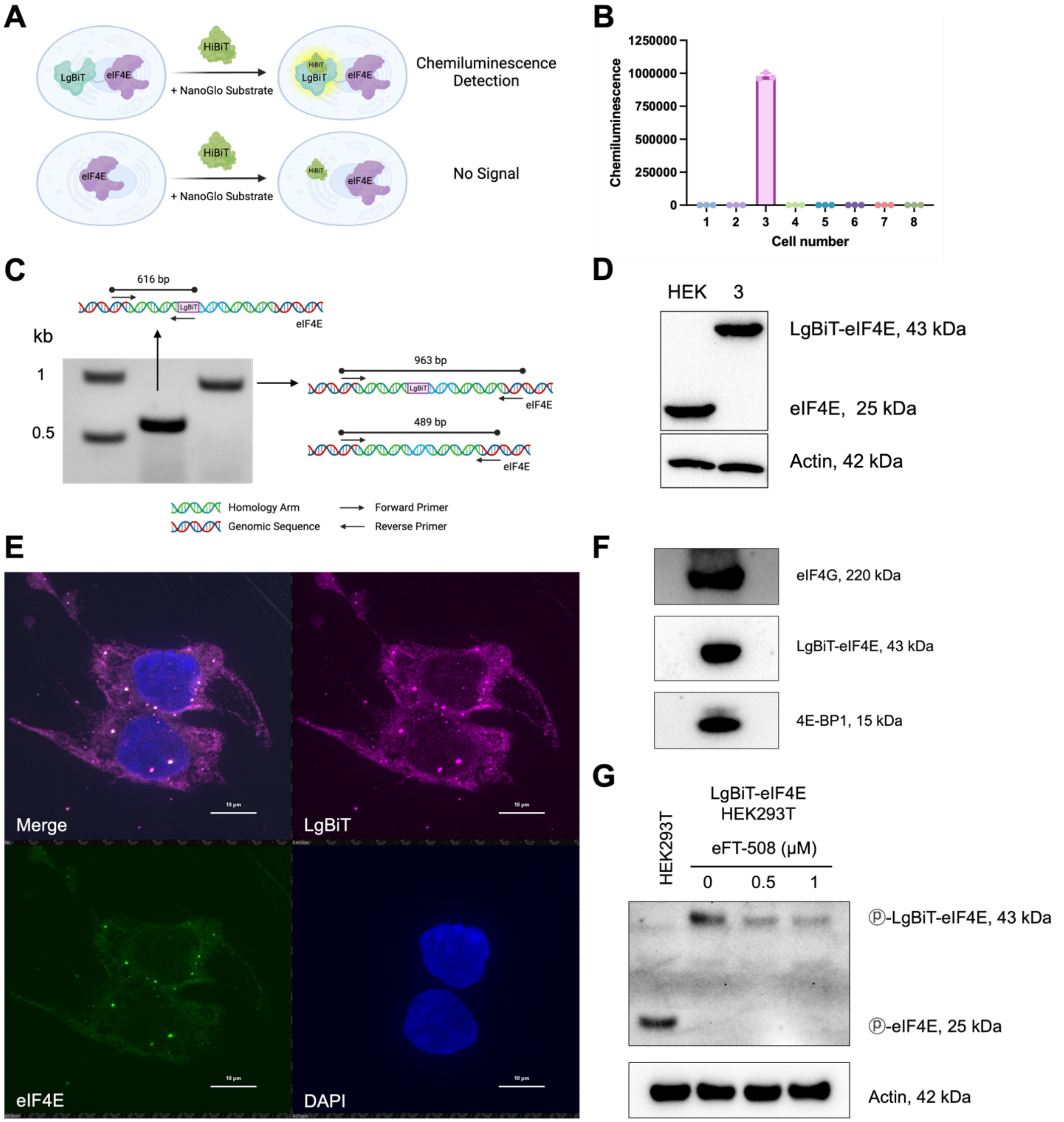


**Figure S8.** Validation of LgBiT-eIF4E HEK293T cells. (A) Scheme for LgBiT lytic detection with HiBiT. (B) Identification of cells with LgBiT incorporation using the HiBiT assay. (C) Junction PCR to confirm LgBiT integration at the correct locus. (D) Western blot confirmation of LgBiT tagging of endogenous eIF4E. (E) Colocalization of eIF4E and LgBiT determined using confocal microscopy. (F) Cap pull-down to assess eIF4E binding to its endogenous protein binding partners, eIF4G and 4E-BP1. (G) Determination of eIF4E phosphorylation capacity.

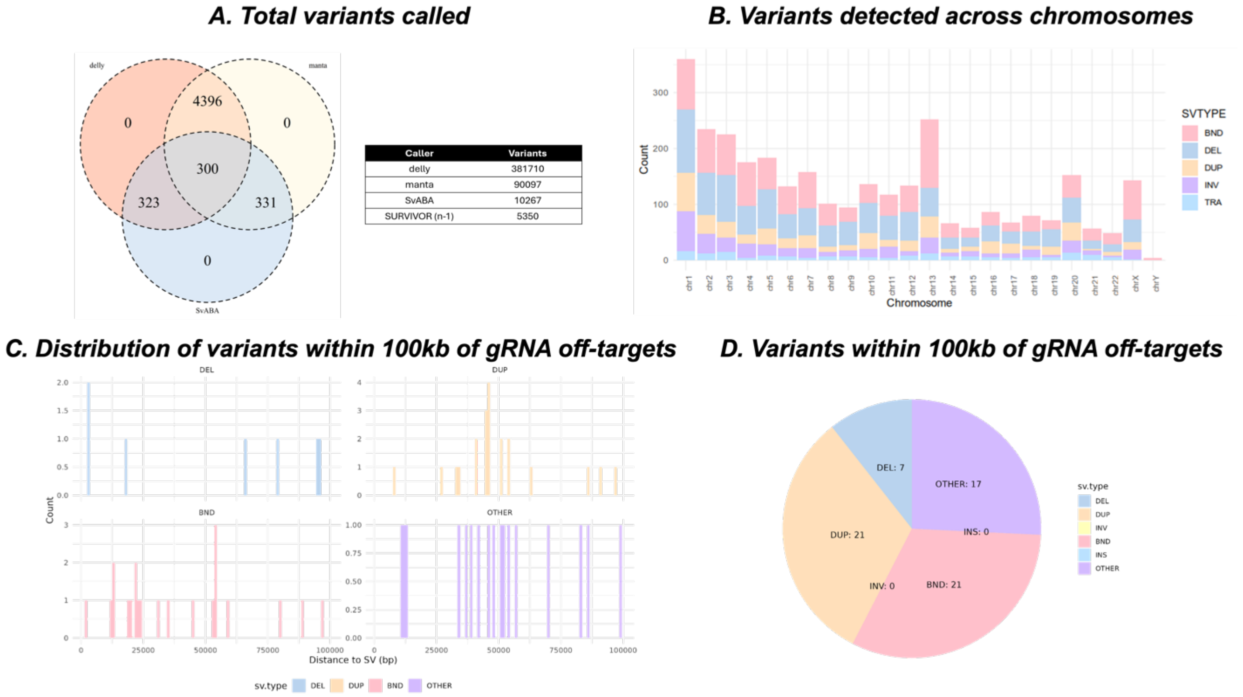


**Figure S9.** (A) Left, total variants called passing n-1 cutoff in survivor. Right, table of total variants in each method. (B) A stacked bar chart of called variants across chromosomes. (C) Distribution of variants within 100kb of gRNA off-targets. (D) Variant types within 100kb of gRNA off-targets.

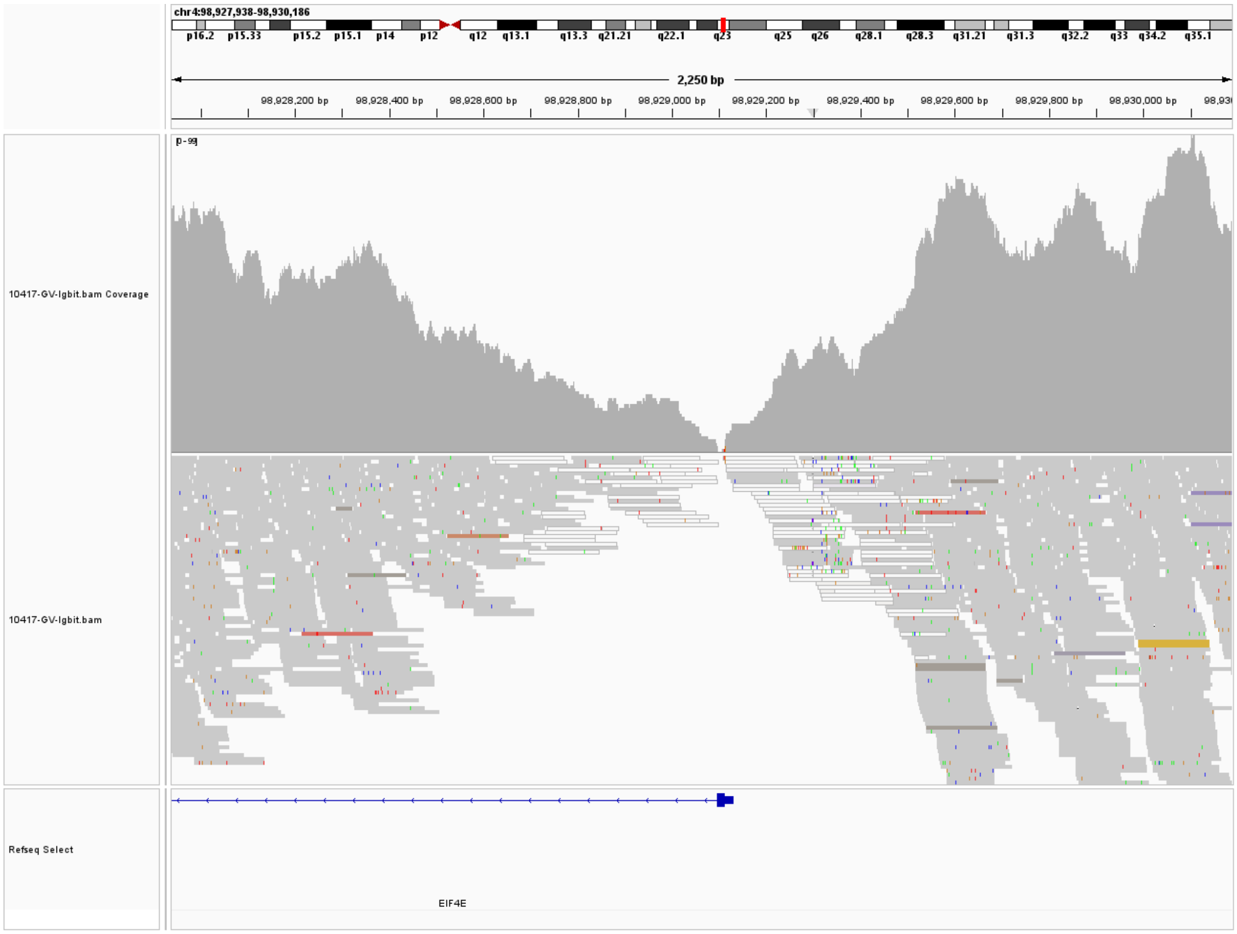


**Figure S10.** IGV screenshot of coverage at eIF4E in alignment to the custom genome showing low coverage to the reference contig near the insertion site. Survivor-based variant calls on our short read data detect no variants at the LgBiT insertion site.

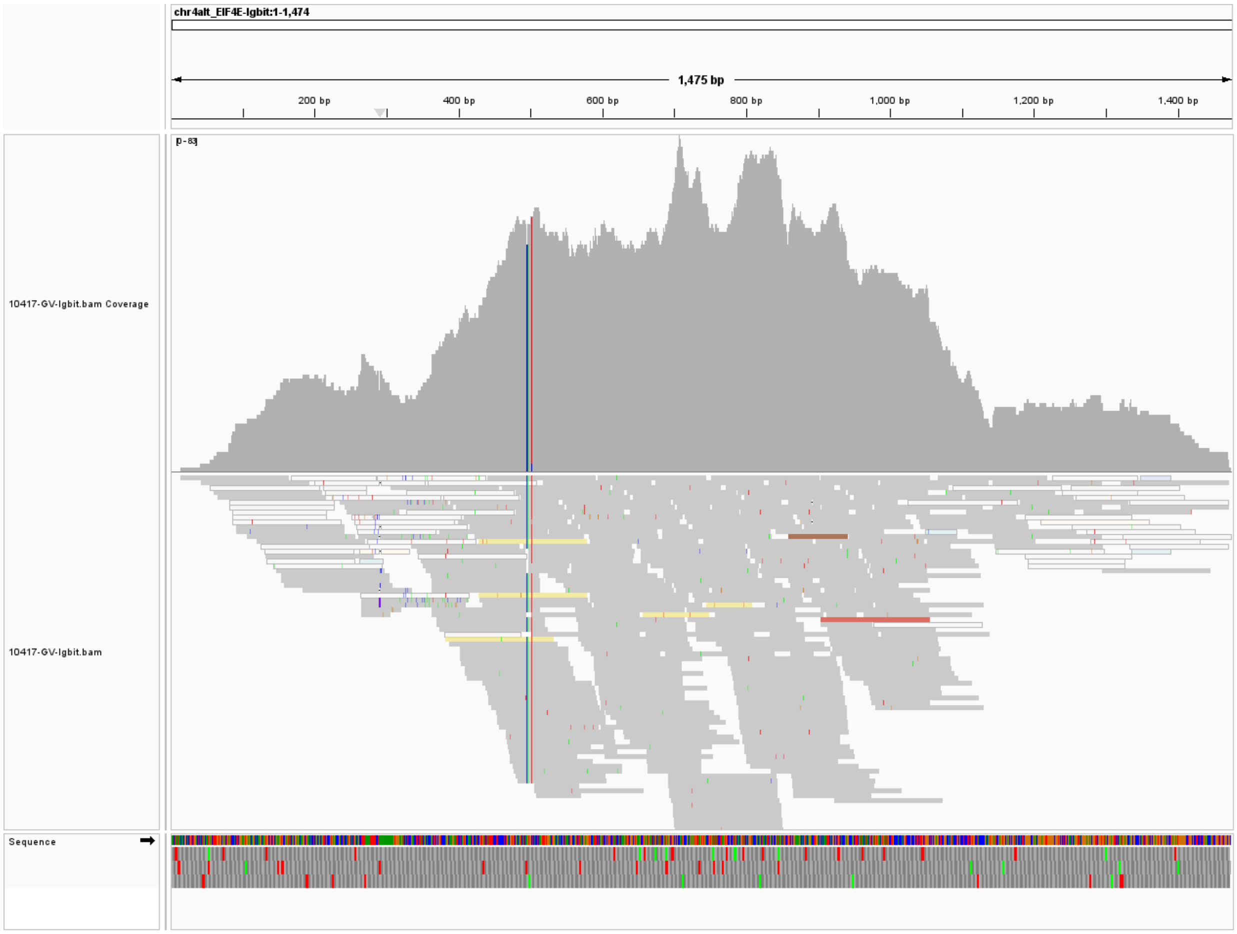


**Figure S11.** IGV screenshot of coverage at LgBiT-eIF4E contig in alignment to the custom genome to visualize LgBiT read coverage.

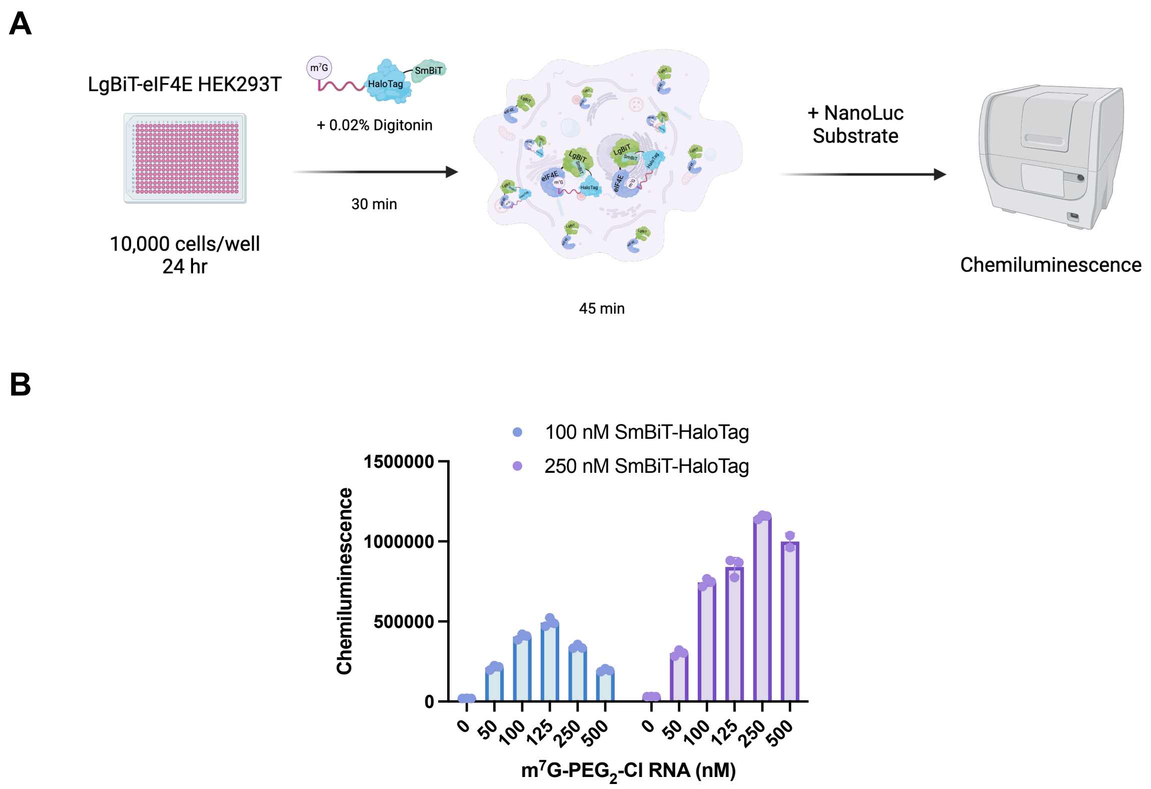


**Figure S12.** Lytic delivery of SmBiT-HaloTag-RNA RNP to LgBiT-eIF4E HEK293T cells. (A) Lytic CRISPR RiPCA. (B) Titration of m$^7$GTP-PEG$_2$-Cl RNA with 100 and 250 nM SmBiT-HaloTag protein in lytic CRISPR RiPCA.

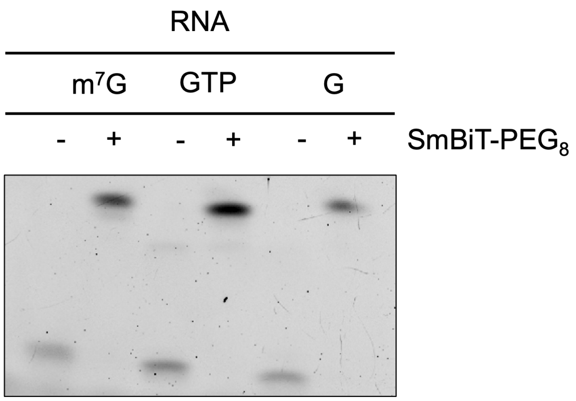


**Figure S13.** Confirmation of the labeling efficiency of RNAs with SmBiT-PEG8 peptide.

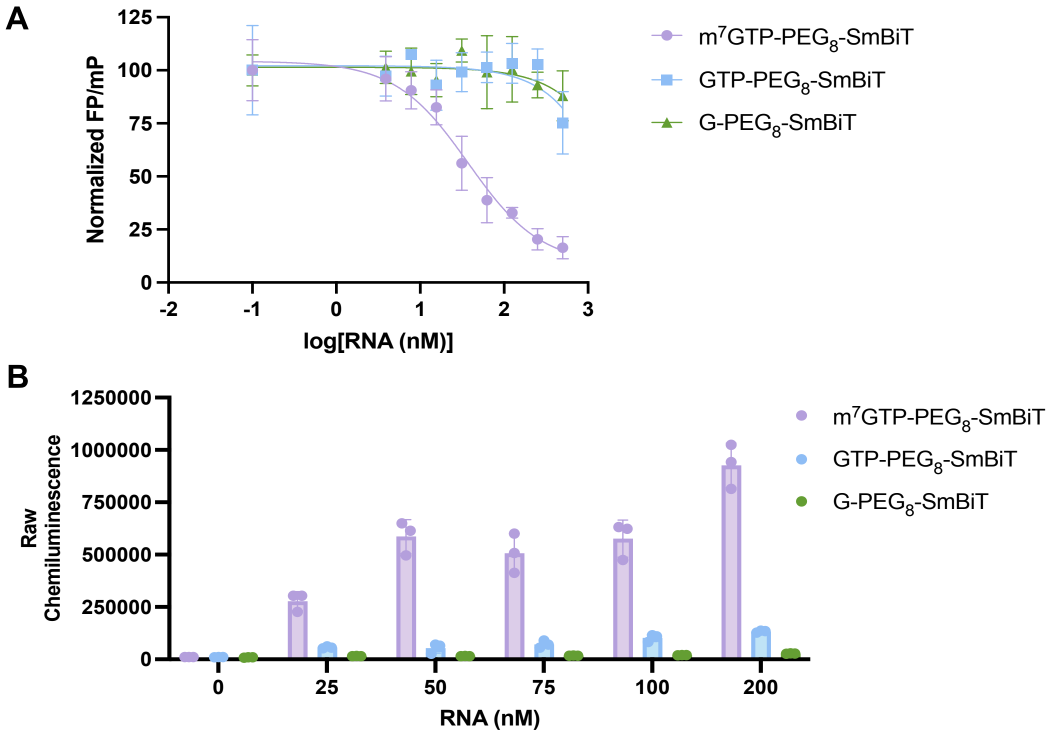


**Figure S14.** Characterization of SmBiT-labeled RNAs. (A) Fluorescence polarization assay to measure the binding affinity of SmBiT-labeled RNAs for eIF4E. (B) Lytic CRISPR RiPCA to demonstrate the specificity of ternary complex formation and signal generation with LgBiT-eIF4E in cell lysate.

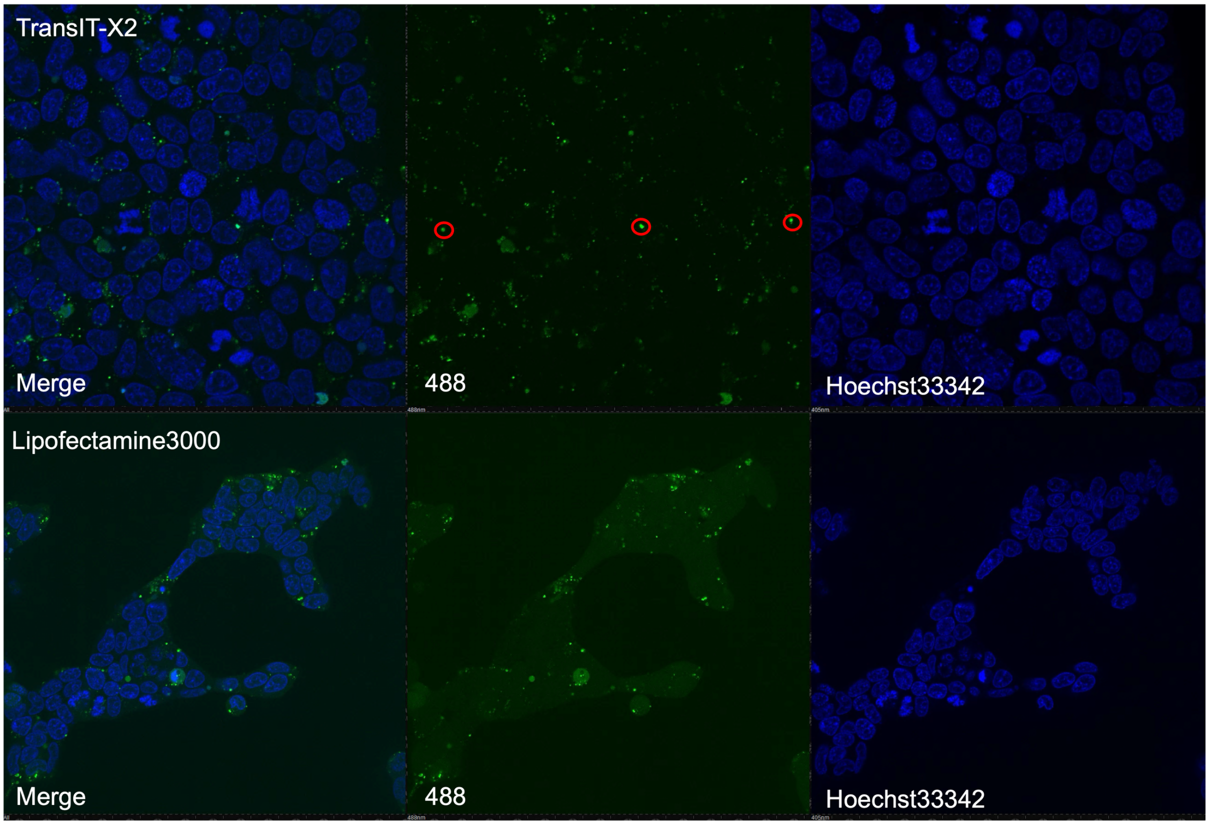


**Figure S15.** Live-cell confocal microscopy of m$^7$G-RNA conjugated to AlexaFluor488 (green) in HEK293T cells with nuclei stained with Hoechst 33342 (blue); circled in red are RNAs likely trapped in endosomes.

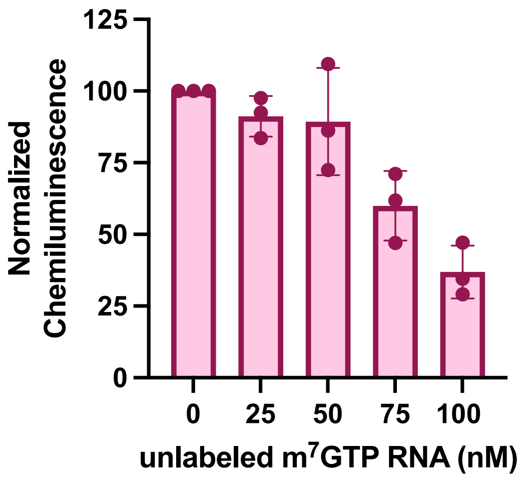


**Figure S16.** Competition of CRISPR RiPCA signal by co-transfection of SmBiT-m$^7$GTP-RNA and non-SmBiT-labeled (unlabeled) m$^7$GTP-RNA.

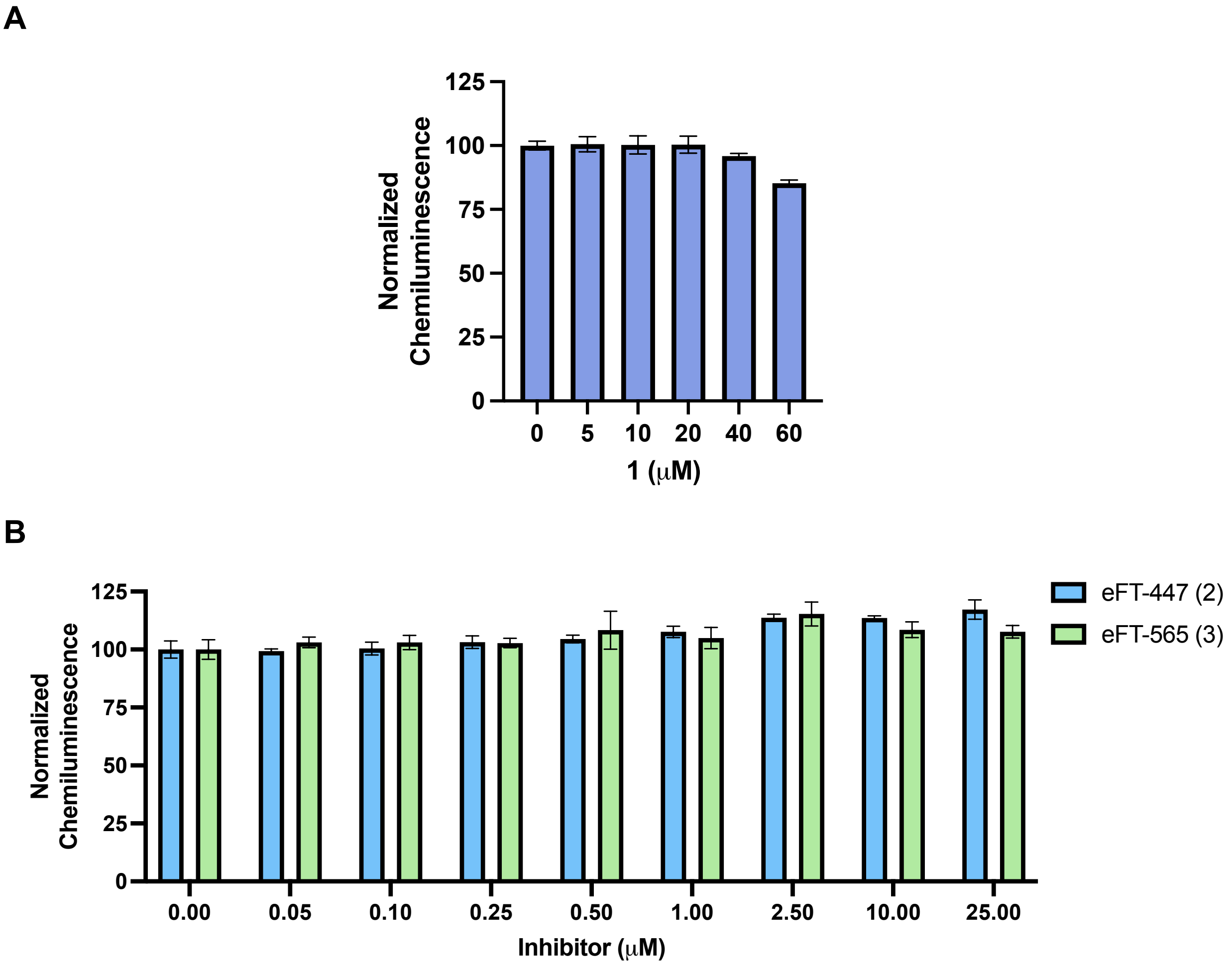


**Figure S17.** CellTiter-Glo® assay to measure cell viability after 6-hour treatment with cap-competitive eIF4E inhibitors (A) **1**, (B) eFT-447 (**2**), and eFT-565 (**3**).

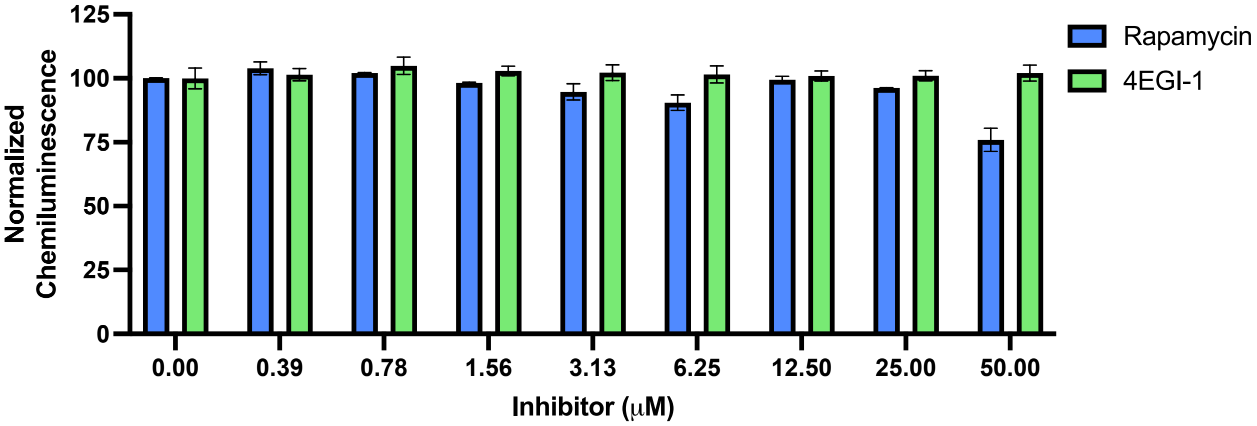


**Figure S18.** CellTiter-Glo® assay to measure cell viability after 6-hour treatment with eIF4E PPI modulators.

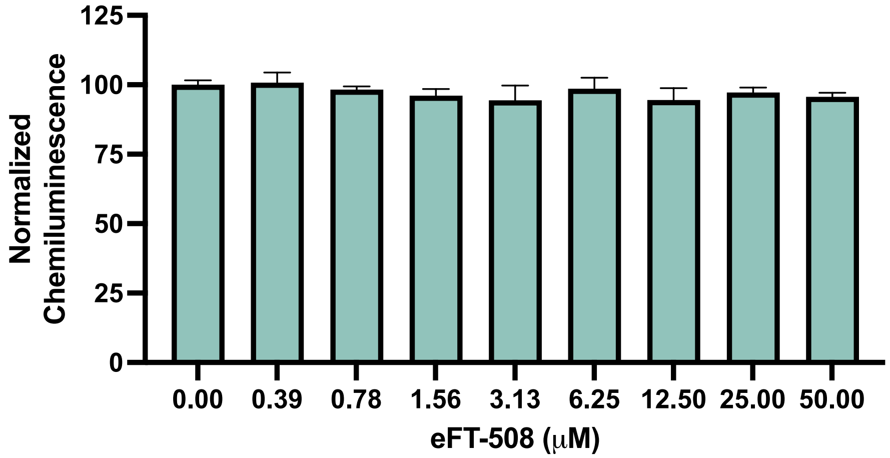


**Figure S19.** CellTiter-Glo® assay to measure cell viability after 6-hour treatment with eIF4E phosphorylation inhibitor.

**Table S1. Sequences and modifications of eIF4E RNA probes.**

| Supplier | 5′ modification | Sequence | Length (nt) | 3′ modification |
|---|---|---|---|---|
| TriLink Biotechnologies | 7-Methylguanosine-triphosphate cap | GCUUCAAGAU | 10 | C7 Amino Linker |
| TriLink Biotechnologies | Guanosine-triphosphate | GCUUCAAGAU | 10 | C7 Amino Linker |
| Horizon Discovery | None | GCUUCAAGAU | 10 | Amino modified C6 |

**Table S2. eIF4E gRNA and LgBiT HDR donor template.** The gRNA sequence shows the 20 nucleotides of the crRNA (IDT), and the HDR (IDT) sequence includes the homology arms (black), LgBiT sequence (purple), and start codon (blue).

| gRNA | HDR donor template |
|---|---|
| 5′ GCGATCAGATCGATCTAAGA | CGTCACGTGGCCAGAAGCTGGCCAATCCGGTTTGA ATCTCATTTTTTTCCTCTTACCCCCCCTTCTGGAGC GGTTGTGCGATCAGATCGATCTACAATG(LgBiT)GC GACTGTCGAACCGGTGAGTATTGCCTTTGGCCCCC ACCCCCACGGGTCCCCGCGCTCCGTCTTCCTTCTG ACTGGGGGACTCCGCGGGACGGCGTTCCC |

**Table S3. Variants within 100kb of gRNA On and Off-targets by Type.**

| sv.id | sv.type | offTarget. chr | offTarget. start | offTarget. end | offTarget. distance |
|---|---|---|---|---|---|
| MantaDEL:156546:0:1:0:0:0 (chr11:58866924) | DEL | chr11 | 58866924 | 58866924 | 2565 |
| MantaDEL:878568:0:1:0:0:0 (chr8:111284635) | DEL | chr8 | 111284635 | 111284635 | 2807 |
| MantaDEL:9:120544:120545:0:0:0 (chr21:10352766) | DEL | chr21 | 10352766 | 10352766 | 17781 |
| MantaDEL:9:28449:242675:0:0:0 (chr1:31357004) | DEL | chr1 | 31357004 | 31357004 | 66027 |
| MantaDEL:532794:0:1:0:0:0 (chr5:100143893) | DEL | chr5 | 100143893 | 100143893 | 78637 |
| MantaDEL:458753:1:2:0:0:0 (chr8:123440851) | DEL | chr8 | 123440851 | 123440851 | 95494 |
| MantaDEL:251311:0:1:0:0:0 (chr16:29249279) | DEL | chr16 | 29249279 | 29249279 | 95549 |
| MantaDUP:TANDEM:862438:0:1:0:0:0 (chr8:61914717) | DUP | chr8 | 61914717 | 61914717 | 7757 |
| MantaDUP:TANDEM:405779:0:1:0:0:0 (chr7:44001123) | DUP | chr7 | 44001123 | 44001123 | 26929 |
| MantaDUP:TANDEM:486122:1:2:0:0:0 (chr8:11963285) | DUP | chr8 | 11963285 | 11963285 | 32579 |

| sv.id | sv.type | offTarget.chr | offTarget.start | offTarget.end | offTarget.distance |
|---|---|---|---|---|---|
| MantaDUP:TANDEM:885319:1:2:0:0:0 (chr8:133900106) | DUP | chr8 | 133900106 | 133900106 | 33570 |
| MantaDUP:TANDEM:758808:0:1:0:0:0 (chr1:32158168) | DUP | chr1 | 32158168 | 32158168 | 40949 |
| MantaDUP:TANDEM:413335:0:1:0:0:0 (chr7:76540265) | DUP | chr7 | 76540265 | 76540265 | 40982 |
| MantaDUP:TANDEM:194207:0:1:0:0:0 (chr10:39203106) | DUP | chr10 | 39203106 | 39203106 | 44924 |
| MantaDUP:TANDEM:194207:0:1:0:0:0 (chr10:39203107) | DUP | chr10 | 39203107 | 39203107 | 44925 |
| MantaDUP:TANDEM:194207:0:1:0:0:0 (chr10:39203108) | DUP | chr10 | 39203108 | 39203108 | 44926 |
| MantaDUP:TANDEM:57394:0:1:0:0:0 (chr13:19249111) | DUP | chr13 | 19249111 | 19249111 | 45868 |
| MantaDUP:TANDEM:9:277581:277583:0:0:0 (chr1:236844093) | DUP | chr1 | 236844093 | 236844093 | 45889 |
| MantaDUP:TANDEM:355628:1:3:0:0:0 (chr5:33326739) | DUP | chr5 | 33326739 | 33326739 | 46197 |
| MantaDUP:TANDEM:355628:1:3:0:0:0 (chr5:33326740) | DUP | chr5 | 33326740 | 33326740 | 46198 |
| MantaDUP:TANDEM:835804:1:2:0:0:0 (chr9:87466481) | DUP | chr9 | 87466481 | 87466481 | 50751 |
| MantaDUP:TANDEM:885319:1:2:0:0:0 (chr8:133917450) | DUP | chr8 | 133917450 | 133917450 | 50914 |
| MantaDUP:TANDEM:95588:0:1:0:0:0 (chr1:225557194) | DUP | chr1 | 225557194 | 225557194 | 54227 |
| MantaDUP:TANDEM:9:179798:179799:0:0:0 (chr4:43765740) | DUP | chr4 | 43765740 | 43765740 | 54388 |
| MantaDUP:TANDEM:269582:1:2:0:0:0 (chr15:43239010) | DUP | chr15 | 43239010 | 43239010 | 63069 |
| MantaDUP:TANDEM:885319:1:2:0:0:0 (chr8:133952110) | DUP | chr8 | 133952110 | 133952110 | 85574 |
| MantaDUP:TANDEM:288026:0:1:0:0:0 (chr15:101578154) | DUP | chr15 | 101578154 | 101578154 | 91335 |
| MantaDUP:TANDEM:451500:1:2:0:0:0 (chr5:21592928) | DUP | chr5 | 21592928 | 21592928 | 96932 |
| MantaBND:76482:5:6:0:0:0:0 (chr20:30852040) | BND | chr20 | 30852040 | 30852040 | 1759 |
| MantaBND:619267:0:1:0:0:0:0 (chr8:70035752) | BND | chr8 | 70035752 | 70035752 | 12120 |
| MantaBND:40650:0:1:0:0:0:0 (chrX:116061728) | BND | chrX | 116061728 | 116061728 | 12615 |

| sv.id | sv.type | offTarget.chr | offTarget.start | offTarget.end | offTarget.distance |
|---|---|---|---|---|---|
| MantaBND:9:120461:120462:0:0:0:0 (chr21:10352766) | BND | chr21 | 10352766 | 10352766 | 12997 |
| MantaBND:57284:0:1:0:0:0:0 (chr21:10352766) | BND | chr21 | 10352766 | 10352766 | 18607 |
| MantaBND:411160:0:1:0:0:0:1 (chr7:67279875) | BND | chr7 | 67279875 | 67279875 | 19862 |
| MantaBND:9:261660:261661:0:0:0:0 (chr1:159480520) | BND | chr1 | 159480520 | 159480520 | 21556 |
| MantaBND:279594:0:1:0:0:0:0 (chr15:77942673) | BND | chr15 | 77942673 | 77942673 | 21845 |
| MantaBND:683025:0:1:0:0:0:1 (chr8:143642131) | BND | chr8 | 143642131 | 143642131 | 23344 |
| MantaBND:482387:0:1:0:0:0:0 (chr8:110752659) | BND | chr8 | 110752659 | 110752659 | 24477 |
| MantaBND:879747:0:1:0:0:0:0 (chr8:114535127) | BND | chr8 | 114535127 | 114535127 | 30805 |
| MantaBND:372572:0:1:0:0:0:0 (chr20:38631495) | BND | chr20 | 38631495 | 38631495 | 35040 |
| MantaBND:287454:0:2:0:0:0:0 (chr15:99826373) | BND | chr15 | 99826373 | 99826373 | 45338 |
| MantaBND:105477:0:1:0:0:0:0 (chr12:42951894) | BND | chr12 | 42951894 | 42951894 | 53399 |
| MantaBND:692477:1:2:0:0:0:1 (chr8:91422579) | BND | chr8 | 91422579 | 91422579 | 53784 |
| MantaBND:692477:1:2:0:0:0:1 (chr8:91422771) | BND | chr8 | 91422771 | 91422771 | 53976 |
| MantaBND:498351:0:3:0:0:0:1 (chr8:74326576) | BND | chr8 | 74326576 | 74326576 | 53982 |
| MantaBND:19647:1:3:0:0:0:1 (chr1:102485594) | BND | chr1 | 102485594 | 102485594 | 58797 |
| MantaBND:110083:0:1:0:0:0:0 (chr15:39782344) | BND | chr15 | 39782344 | 39782344 | 79938 |
| MantaBND:619267:0:1:0:0:0:0 (chr8:70112834) | BND | chr8 | 70112834 | 70112834 | 89202 |
| MantaBND:344436:1:2:0:0:0:1 (chr22:29511422) | BND | chr22 | 29511422 | 29511422 | 97341 |
| 279096586:2 (chr22:11558025) | OTHER | chr22 | 11558025 | 11558025 | 10641 |
| 3152198753:2 (chr8:115410605) | OTHER | chr8 | 115410605 | 115410605 | 11880 |
| 837392215:2 (chr8:30261445) | OTHER | chr8 | 30261445 | 30261445 | 13071 |
| 4044261947:1 (chr8:133900106) | OTHER | chr8 | 133900106 | 133900106 | 33997 |

| sv.id | sv.type | offTarget.chr | offTarget.start | offTarget.end | offTarget.distance |
|---|---|---|---|---|---|
| 2214182814:2 (chr20:5324281) | OTHER | chr20 | 5324281 | 5324281 | 36517 |
| 837392215:2 (chr8:30287778) | OTHER | chr8 | 30287778 | 30287778 | 39404 |
| 837392215:2 (chr8:30290570) | OTHER | chr8 | 30290570 | 30290570 | 42196 |
| 837392215:2 (chr8:30294590) | OTHER | chr8 | 30294590 | 30294590 | 46216 |
| 1674348823:2 (chr15:91553931) | OTHER | chr15 | 91553931 | 91553931 | 48326 |
| 4044261947:1 (chr8:133917450) | OTHER | chr8 | 133917450 | 133917450 | 51341 |
| 4034198010:2 (chr22:32476762) | OTHER | chr22 | 32476762 | 32476762 | 52110 |
| 2332044138:1 (chr8:62996303) | OTHER | chr8 | 62996303 | 62996303 | 54134 |
| 2439647863:2 (chr15:50953899) | OTHER | chr15 | 50953899 | 50953899 | 56796 |
| 888279940:1 (chr8:15501904) | OTHER | chr8 | 15501904 | 15501904 | 70049 |
| 901556159:1 (chr1:241555796) | OTHER | chr1 | 241555796 | 241555796 | 83151 |
| 4044261947:1 (chr8:133952110) | OTHER | chr8 | 133952110 | 133952110 | 86001 |
| 3152198753:2 (chr8:115497887) | OTHER | chr8 | 115497887 | 115497887 | 99162 |

**Table S4. Small molecule inhibitors tested in this study**.

| Small molecule | Target | Reported Activity | Reference |
|---|---|---|---|
| **1** | Prodrug that targets eIF4E's cap-binding site | 4.1 μM (FP) | *ACS Med. Chem. Lett.* **2025**, *16*, 96 |
| eFT-447 (**2**) | Small molecule targeting eIF4E's cap-binding site | 85 nM (FP) | WO/2021/003157 |
| eFT-565 (**3**) | Small molecule targeting eIF4E's cap-binding site | 30 nM (FP) | WO/2021/003157 |
| Rapamycin | Allosteric inhibitor of mTORC1; inhibits 4E-BP1 phosphorylation | 12 nM (SPR) | *JACS* **2005**, *127*, 4715 |
| 4EGI-1 | Allosteric modulator of the eIF4E-eIF4G protein-protein interaction | 25 μM (FP) | *Cell* **2007**, *128*, 257 |
| eFT-508 | Inhibitor of MNK1/2; inhibits eIF4E phosphorylation | 2.4/1 nM (ADP-Glo) | *J. Med. Chem.* **2018**, *61*, 3516 |

#### D. Characterization Data for Compounds 2 and 3

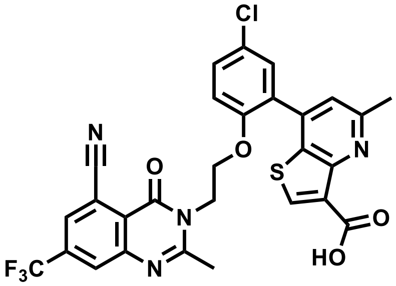

*7-(5-chloro-2-(2-(5-cyano-2-methyl-4-oxo-7-(trifluoromethyl)quinazolin-3(4H)-yl)ethoxy)phenyl)-5-methylthieno[3,2-b]pyridine-3-carboxylic acid (**2**)*. $^1$H NMR (400 MHz, $CDCl_3$) δ 13.83 (s, 1H), 8.32 (s, 1H), 8.05 (s, 1H), 7.98 (s, 1H), 7.46 (dd, J = 8.9, 2.6 Hz, 1H), 7.27 (d, J = 3.9 Hz, 1H), 7.18 (s, 1H), 7.02 (d, J = 8.9 Hz, 1H), 4.43 (t, J = 4.8 Hz, 2H), 4.31 (t, J = 4.9 Hz, 2H), 2.79 (s, 3H), 1.83 (s, 3H). $^{13}$C NMR (101 MHz, $CDCl_3$) δ 161.77, 158.97, 157.44, 156.94, 153.19, 151.41, 148.20, 142.87, 140.31, 140.26, 136.26, 135.92, 131.61, 131.12, 130.12, 130.03, 129.61, 129.57, 129.45, 127.31, 126.99, 125.90, 121.99, 120.86, 120.77, 116.34, 113.77, 112.01, 77.35, 77.03, 76.72, 65.67, 44.90, 24.10, 22.53, 22.51. HRMS (ESI$^+$): 599.079 $[M+1]^+$.

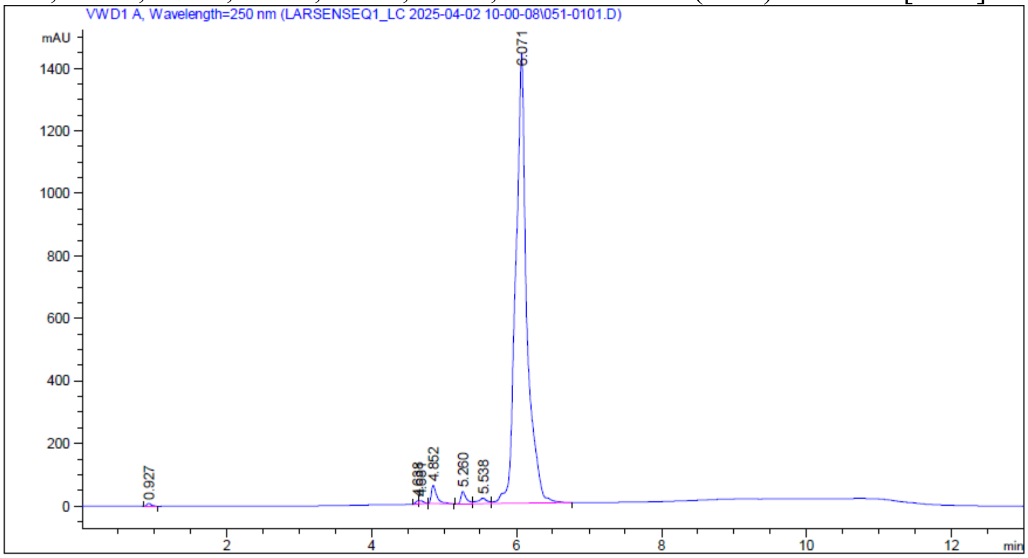


| Peak # | RetTime [min] | Type | Width [min] | Area [mAU*s] | Height [mAU] | Area % |
|---|---|---|---|---|---|---|
| 1 | 0.927 | VB | 0.0649 | 39.99218 | 9.35134 | 0.2478 |
| 2 | 4.638 | BV | 0.0464 | 30.61092 | 10.09453 | 0.1897 |
| 3 | 4.681 | VB | 0.0616 | 41.32732 | 9.91685 | 0.2561 |
| 4 | 4.852 | BB | 0.0759 | 304.64767 | 58.38791 | 1.8880 |
| 5 | 5.260 | BV | 0.0730 | 200.26624 | 39.61590 | 1.2411 |
| 6 | 5.538 | VV | 0.1207 | 160.73929 | 17.91151 | 0.9962 |

| Peak # | RetTime [min] | Type | Width [min] | Area [mAU*s] | Height [mAU] | Area % |
|---|---|---|---|---|---|---|
| 7 | 6.071 | VB | 0.1413 | 1.53581e4 | 1441.01074 | 95.1810 |

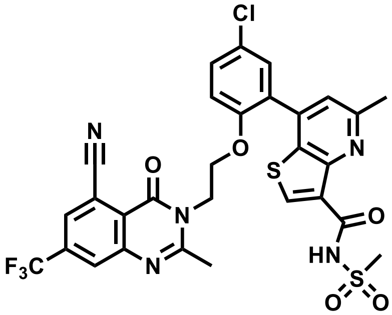

*7-(5-chloro-2-(2-(5-cyano-2-methyl-4-oxo-7-(trifluoromethyl)quinazolin-3(4H)-yl)ethoxy)phenyl)-5-methyl-N-(methylsulfonyl)thieno[3,2-b]pyridine-3-carboxamide (**3**).* $^{1}H$ NMR (400 MHz, $CDCl_3$) δ 13.11 (s, 1H), 8.37 (s, 1H), 8.04 (dt, J = 1.7, 0.8 Hz, 1H), 7.98 (d, J = 1.7 Hz, 1H), 7.45 (ddd, J = 8.9, 2.6, 0.7 Hz, 1H), 7.24 (dd, J = 2.7, 0.7 Hz, 1H), 7.16 (s, 1H), 6.99 (d, J = 8.9 Hz, 1H), 4.42 (t, J = 4.8 Hz, 2H), 4.29 (t, J = 4.8 Hz, 2H), 3.51 (d, J = 0.7 Hz, 3H), 2.79 (s, 3H), 1.77 (s, 3H). $^{13}C$ NMR (101 MHz, $cdcl_3$) δ 159.48, 159.02, 157.48, 157.45, 153.17, 150.33, 148.24, 142.51, 140.58, 140.53, 136.21, 135.87, 132.24, 131.04, 130.12, 129.58, 127.15, 126.95, 126.90, 122.06, 120.75, 120.66, 116.36, 113.33, 112.06, 77.34, 77.03, 76.71, 65.43, 44.92, 42.06, 42.01, 24.48, 24.41, 22.48. HRMS (ESI$^{+}$): 676.076 $[M+1]^{+}$.

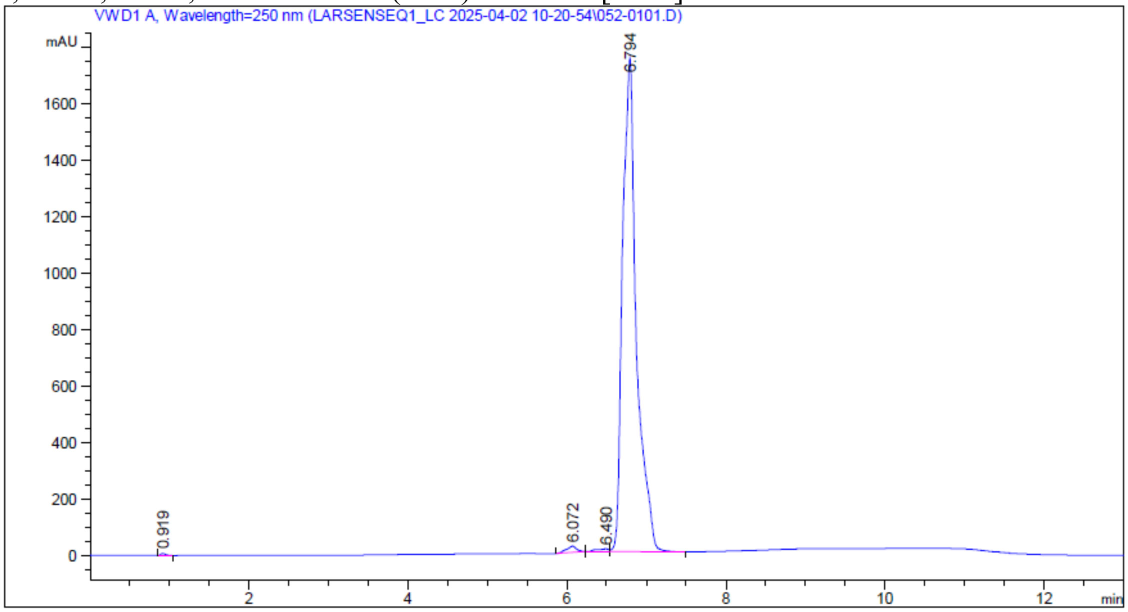


| Peak # | RetTime [min] | Type | Width [min] | Area [mAU*s] | Height [mAU] | Area % |
|---|---|---|---|---|---|---|
| 1 | 0.919 | VB | 0.0679 | 33.02919 | 7.28365 | 0.1534 |
| 2 | 6.072 | BB | 0.1214 | 201.86620 | 22.78627 | 0.9376 |
| 3 | 6.490 | BV | 0.1429 | 110.60599 | 10.17002 | 0.5138 |
| 4 | 6.794 | VB | 0.1602 | 2.11835e4 | 1750.43359 | 98.3952 |

HRMS spectrum of **2**

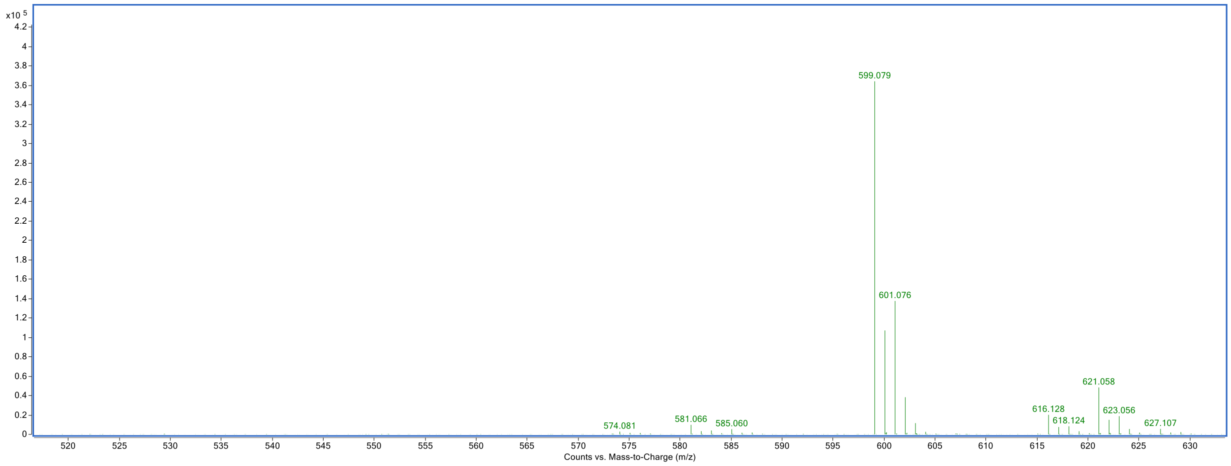


HRMS spectrum of **3**

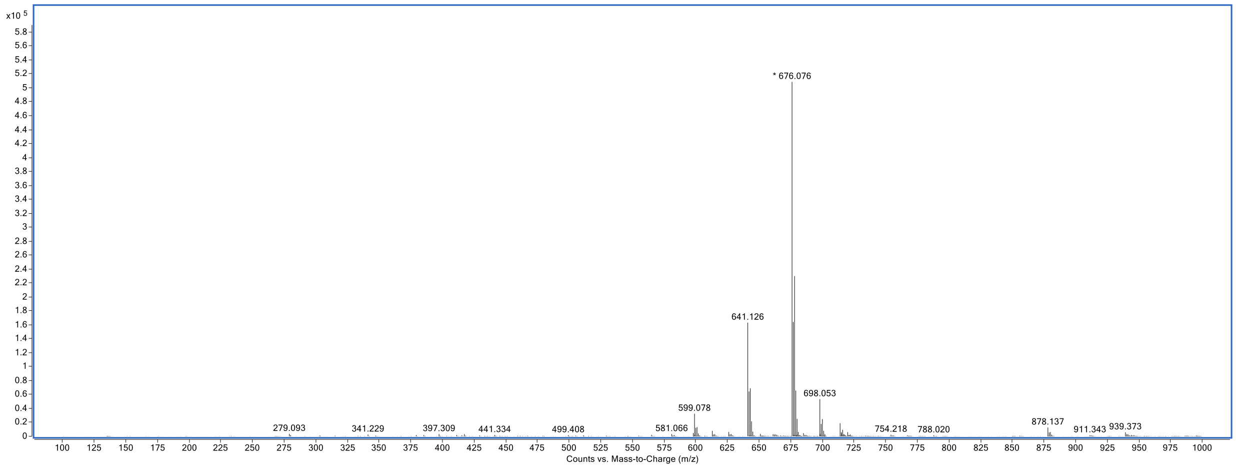


**Table S5.** Analytical purity determined by RP-HPLC

| Inhibitor | Purity (%) |
|---|---|
| **2** | 95.1 |
| **3** | 98.4 |

### NMR Spectra for Compounds 2 and 3

#### Compound 2

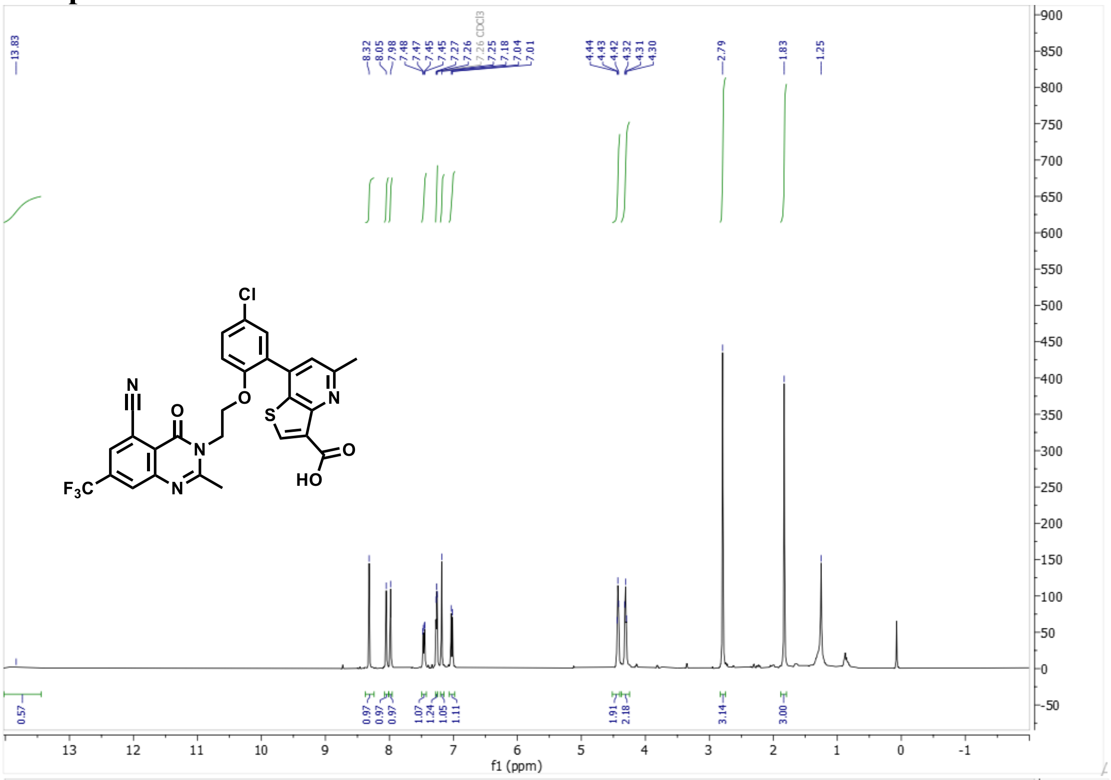

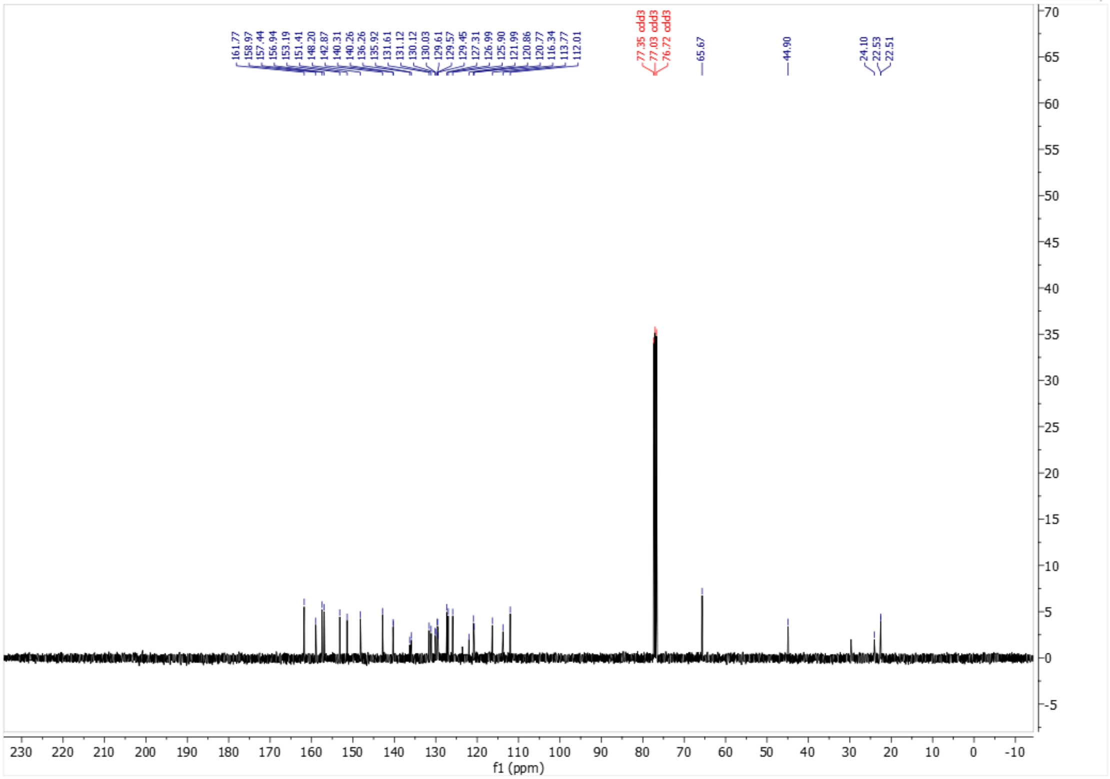

#### Compound 3

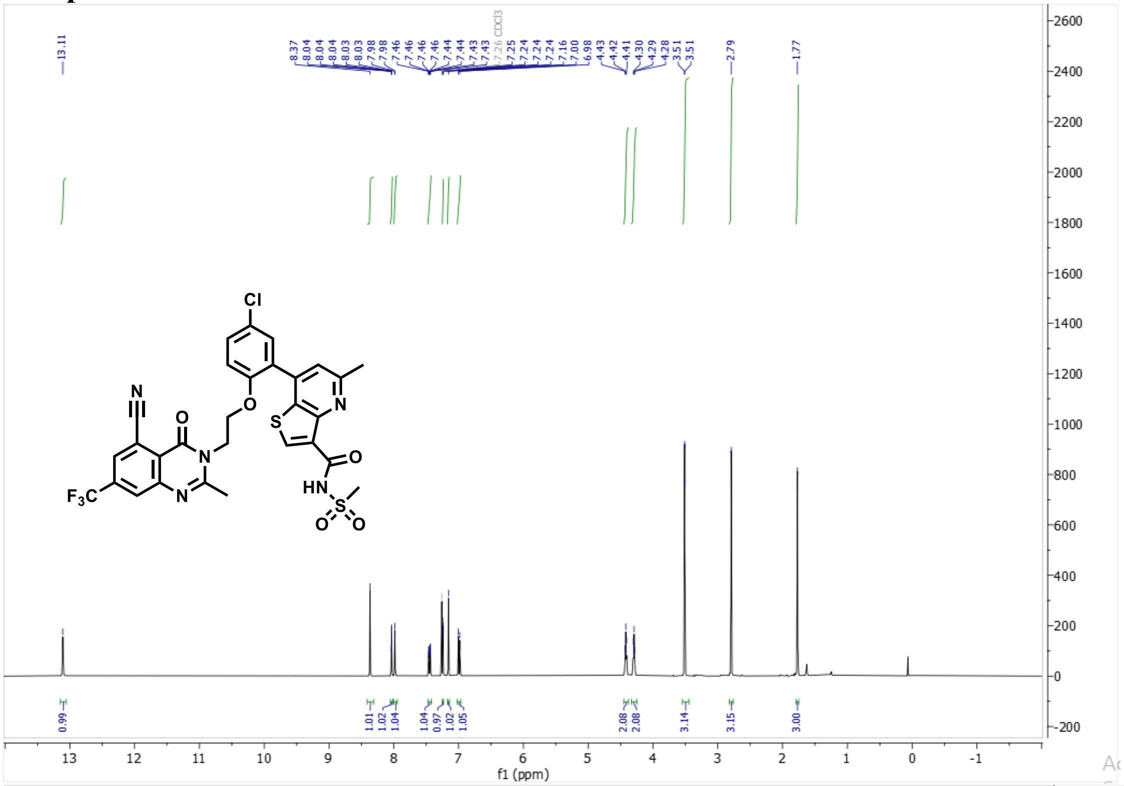

13.11
8.37
8.04
8.04
8.04
8.03
8.03
7.98
7.98
7.46
7.46
7.46
7.46
7.44
7.44
7.43
7.43
7.26 CDCl3
7.25
7.24
7.24
7.24
7.24
7.16
7.00
6.98
4.43
4.42
4.41
4.30
4.29
4.28
3.51
3.51
2.79
1.77
Cl
N
O
O
N
S
N
O
F3C
N
HN
S
O
O
0.99
1.01
1.02
1.04
1.04
0.97
1.02
1.05
2.08
2.08
3.14
3.15
3.00
f1 (ppm)


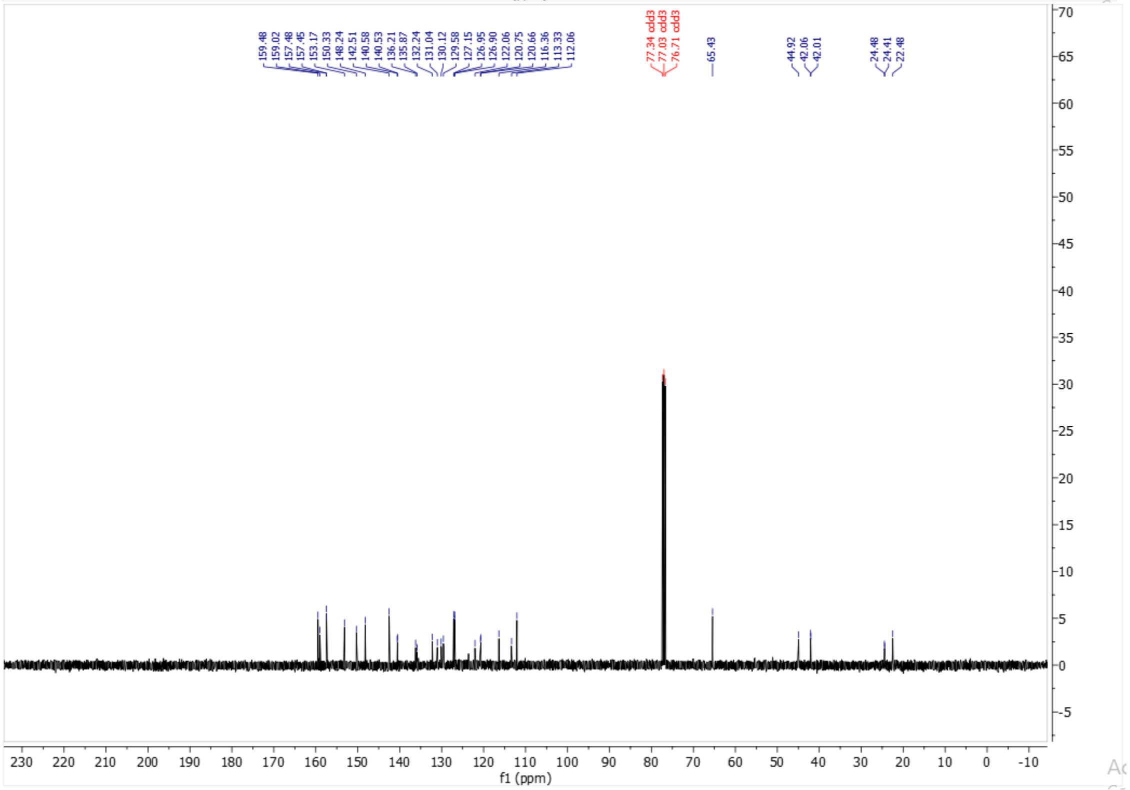

159.48
159.02
157.48
157.45
153.17
150.33
148.24
142.51
140.58
140.53
136.21
135.87
132.24
131.04
130.12
129.58
127.15
126.95
126.90
122.06
120.75
120.66
116.36
113.33
112.06
77.34 cdcl3
77.03 cdcl3
76.71 cdcl3
65.43
44.92
42.06
42.01
24.48
24.41
22.48
f1 (ppm)

#### D. Characterization Data for Compounds 2 and 3

**General Chemistry Materials and Methods.** All purchased solvents and reagents were used without further purification. Reactions were monitored by thin-layer chromatography (TLC) carried out on 20 cm Analtech silica gel plates (LOT# 28923) using UV-light (254 nm). Flash chromatography was performed using Redisep® Silver Flash columns on a Combiflash Nextgen 300+. Inhibitor **2** was purified by C18 reverse-phase preparative HPLC with solvent A (0.1% TFA in $H_2O$) and solvent B (0.1% TFA in $CH_3CN$) as eluents. General Method A: 50-70% B over 20 min at 60 mL /min. NMR spectra were performed on a 400 MHz Varian instrument calibrated using tetramethylsilane (TMS) as an internal reference. Chemical shifts ($\delta$ values) are reported in parts per million (ppm) and are referenced to the deuterated residual solvent peak. High resolution Mass spectrometry (HRMS) was performed using an Agilent Q-TOF HPLC-MS spectrometer using ESI ionization with an accuracy of 2 ppm. All compounds were found to be >95% pure by HPLC analysis.

#### Synthetic Methods

**Scheme 1.** Synthesis of **2**

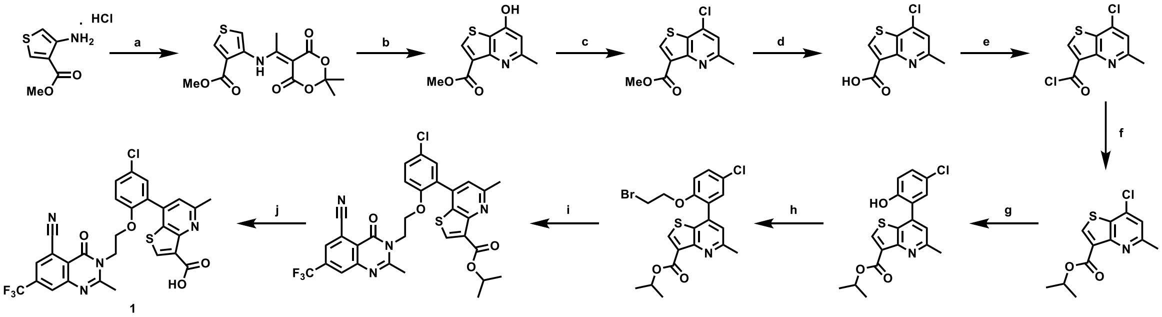

Reagents and conditions: (a) methyl 4-aminothiophene-3-carboxylate hydrochloride, Meldrum's acid, triethylorthoacetate, 90 °C; (b) Dowtherm A, 235 °C; (c) $POCl_3$, DCE, 90 °C; (d) THF/MeOH/$H_2O$, LiOH · $H_2O$, 23 °C; (e) $SOCl_2$, DCM, 23 °C; (f) IPA, 23 °C; (g) (5-chloro-2-hydroxyphenyl)boronic acid, Pd(dppf)$Cl_2$, $K_2CO_3$, 1,4-dioxane/$H_2O$, 90 °C; (h) 1,2-dibromoethane, $K_2CO_3$, acetone, 45 °C; (i) 2-methyl-4-oxo-7-(trifluoromethyl)-3,4-dihydroquinazoline-5-carbonitrile, $K_2CO_3$, DMF, 60 °C; (j) $SnMe_3OH$, DCE, 90 °C.

**2**, step a: *methyl 4-((1-(2,2-dimethyl-4,6-dioxo-1,3-dioxan-5-ylidene)ethyl)amino)thiophene-3-carboxylate*. 2,2-dimethyl-1,3-dioxane-4,6-dione (8.19 g, 56.8 mmol, 1.1 eq) was added to an oven-dried round-bottom-flask followed by triethylorthoacetate (0.69 M, 75 mL). A condenser was attached, and the mixture was stirred at 90 °C for 2 h under $N_2$. Then, methyl 4-aminothiophene-3-carboxylate hydrochloride (10.00 g, 51.6 mmol, 1.0 eq) was added portion wise under $N_2$ at 90 °C and heating continued overnight. The reaction mixture was cooled to room temperature, water added and extracted with EtOAc (50 mL x 3). The organic layer was dried over anhydrous $Na_2SO_4$, filtered and concentrated *in vacuo* to afford a viscous yellow oil which was purified by flash column chromatography (Rf: 0.35, 0-80% EtOAc in hexanes) to provide the product as off-white solids (5.55 g, 33%). $^1$H NMR (400 MHz, DMSO-$d_6$) δ 12.69 (s, 1H), 8.48 (d, $J$ = 3.4 Hz, 1H), 7.79 (dd, $J$ = 3.4, 0.8 Hz, 1H), 3.77 (s, 3H), 2.53 (s, 3H), 1.66 (s, 6H).

**2**, step b: *methyl 7-hydroxy-5-methylthieno[3,2-b]pyridine-3-carboxylate*. Methyl 4-((1-(2,2-dimethyl-4,6-dioxo-1,3-dioxan-5-ylidene)ethyl)amino)thiophene-3-carboxylate (5.55 g, 17.1 mmol) was added to an oven-dried pressure vessel followed by Dowtherm A (0.34 M, 50 mL). The mixture was sparged with $N_2$ for 5 min before sealing the tube and heating at 235 °C overnight. The reaction mixture was cooled to room temperature and 100 mL $Et_2O$ was added. The precipitated solids were filtered and washed with $Et_2O$ (25 mL x 5) to afford the product as light-brown solids (2.18 g, 57%). $^1$H NMR (400 MHz, DMSO-$d_6$) δ 10.98 (s, 1H), 8.79 (s, 1H), 6.00 (dd, $J$ = 1.7, 0.8 Hz, 1H), 3.91 (s, 3H), 2.43 (d, $J$ = 0.7 Hz, 3H).

**2**, step c: *methyl 7-chloro-5-methylthieno[3,2-b]pyridine-3-carboxylate*. Methyl 7-hydroxy-5-methylthieno[3,2-b]pyridine-3-carboxylate (2.176 g, 9.75 mmol, 1.0 eq) was added to an oven-dried round bottom flask followed by anhydrous DCE (0.23 M, 40 mL). Then, $POCl_3$ (2.73 mL, 29.3 mmol, 3.0 eq) was added followed by a catalytic amount of DMF. The mixture was warmed to 90 °C and stirred under $N_2$ overnight. The reaction mixture was concentrated, diluted with ice cold water (100 mL), and basified with 15% NaOH to pH 10 followed by extraction with DCM (50 mL x 3). The organic layer was dried over anhydrous $Na_2SO_4$, filtered and concentrated *in vacuo* to afford the crude product which was purified by flash column chromatography ($R_f$: 0.4, 0-

100% EtOAc in hexanes) to afford the product as white solids (1.80 g, 76%). $^{1}H$ NMR (400 MHz, DMSO-$d_6$) δ 8.96 (d, $J$ = 0.4 Hz, 1H), 7.62 (s, 1H), 3.87 (s, 3H), 2.64 (s, 3H).

**2**, step d: *7-chloro-5-methylthieno[3,2-b]pyridine-3-carboxylic acid*. Methyl 7-chloro-5-methylthieno[3,2-b]pyridine-3-carboxylate (1.80 g, 7.47 mmol, 1.0 eq) was added to an oven-dried round bottom flask followed by a 3:3:1 solvent mixture (0.35 M, 21 mL) of MeOH/$H_2O$/THF, respectively. Then, LiOH · $H_2O$ (626.6 mg, 14.934 mmol, 2.0 eq) was added and the resulting mixture stirred at room temperature overnight. The solids were filtered, the filtrate was concentrated and combined with the solids. The combined solids were acidified using saturated citric acid to pH 1 and filtered. The resulting solids were washed with MeOH (15 mL x 3), then $Et_2O$ (25 mL x 3) to afford the product as off-white solids (1.39 g, 82%). $^{1}H$ NMR (400 MHz, DMSO-$d_6$) δ 13.21 (s, 1H), 8.98 (s, 1H), 7.67 (s, 1H), 2.67 (s, 3H).

**2**, step e: *7-chloro-5-methylthieno[3,2-b]pyridine-3-carbonyl chloride*. 7-chloro-5-methylthieno[3,2-b]pyridine-3-carboxylic acid (1.39 g, 6.11 mmol, 1.0 eq) was added to an oven-dried round bottom flask followed by anhydrous DCM (40 mL). The mixture was cooled to 0 °C, then oxalyl chloride (0.80 mL, 9.16 mmol, 1.5 eq) was added dropwise. A catalytic amount of DMF was added and the resulting mixture was stirred at room temperature for 2 h. The mixture was concentrated *in vacuo* to afford off-white solids and used directly without further purification (assumed quantitative yield). $^{1}H$ NMR (400 MHz, DMSO-$d_6$) δ 8.99 (s, 1H), 7.69 (s, 1H), 2.68 (s, 4H).

**2**, step f: *isopropyl 7-chloro-5-methylthieno[3,2-b]pyridine-3-carboxylate*. To the round bottom flask containing 7-chloro-5-methylthieno[3,2-b]pyridine-3-carbonyl chloride was added isopropanol (0.12 M, 50 mL) and stirred at room temperature overnight. The reaction mixture was concentrated *in vacuo*, diluted with 25 mL DCM, then neutralized with saturated $NaHCO_3$. The aqueous layer was extracted with DCM (25 mL x 3), the combined organic layer was dried over anhydrous $Na_2SO_4$, filtered and concentrated *in vacuo* to afford a yellow oil which was purified by flash column chromatography (Rf: 0.4, 0-50% EtOAc in hexanes) to afford the product as white solids (1.45 g, 88%). $^1$H NMR (400 MHz, DMSO-$d_6$) δ 8.89 (d, $J$ = 0.4 Hz, 1H), 7.61 (s, 1H), 5.15 (p, $J$ = 6.2 Hz, 1H), 2.64 (s, 3H), 1.34 (d, $J$ = 6.2 Hz, 6H).

**2**, step g: *isopropyl 7-(5-chloro-2-hydroxyphenyl)-5-methylthieno[3,2-b]pyridine-3-carboxylate*. Isopropyl 7-chloro-5-methylthieno[3,2-b]pyridine-3-carboxylate (700.0 mg, 2.59 mmol, 1.0 eq), (5-chloro-2-hydroxyphenyl)boronic acid (536.8 mg, 3.11 mmol, 1.2 eq), and $K_2CO_3$ (717.3 mg, 5.19 mmol, 2.0 eq) were added to an oven-dried pressure tube followed by anhydrous dioxane (0.17 M, 10 mL) and DI $H_2O$ (5 mL). The mixture was sparged with $N_2$ for 5 min before adding Pd(dppf)$Cl_2$ (94.9 mg, 2.59 mmol, 0.05 eq), sealing it and heating to 100 °C overnight. The reaction mixture was cooled to room temperature, diluted with EtOAc, and washed with water. The aqueous layer was extracted with EtOAc (25 mL x 3), the organic layer washed with brine, dried over anhydrous $Na_2SO_4$, then filtered and concentrated *in vacuo* to afford a brown oil which was purified via flash column chromatography ($R_f$: 0.6, 0-100% EtOAc in hexanes) to afford the product as white solids (700.0 mg, 75%). $^1$H NMR (400 MHz, DMSO-$d_6$) δ 10.28 (s, 1H), 8.77 (s, 1H), 7.41 – 7.37 (m, 2H), 7.34 (s, 1H), 7.08 – 7.01 (m, 1H), 5.16 (p, $J$ = 6.2 Hz, 1H), 2.66 (s, 3H), 1.35 (d, $J$ = 6.3 Hz, 6H).

**2**, step h: *isopropyl 7-(2-(2-bromoethoxy)-5-chlorophenyl)-5-methylthieno[3,2-b]pyridine-3-carboxylate*. Isopropyl 7-(5-chloro-2-hydroxyphenyl)-5-methylthieno[3,2-b]pyridine-3-carboxylate (700.0 mg, 1.93 mmol, 1.0 eq), $K_2CO_3$ (935.7 mg, 6.77 mmol, 3.5 eq), and anhydrous acetone (0.18 M, 10 mL) were added to an oven dried round bottom flask. Then, dibromoethane (0.836 mL, 9.67 mmol, 5.0 eq) was added and the reaction heated to 45 °C under $N_2$ for 2 days. The reaction mixture was cooled to room temperature, filtered and washed with acetone (15 mL x 5), and the filtrate concentrated to afford a pale-yellow oil which was purified by flash column chromatography ($R_f$: 0.7, 0-100% EtOAc in hexanes) to afford the product as a yellow foam (645.0 mg, 71%). $^1$H NMR (400 MHz, DMSO-d$_6$) δ 8.77 (s, 1H), 7.59 – 7.54 (m, 2H), 7.42 (s, 1H), 7.30 (d, $J$ = 8.7 Hz, 1H), 5.17 (p, $J$ = 6.3 Hz, 1H), 4.37 (dd, $J$ = 6.1, 4.7 Hz, 2H), 3.66 – 3.59 (m, 2H), 2.67 (s, 3H), 1.35 (d, $J$ = 6.2 Hz, 6H).

**2**, step i: *isopropyl 7-(5-chloro-2-(2-(5-cyano-2-methyl-4-oxo-7-(trifluoromethyl)quinazolin-3(4H)-yl)ethoxy)phenyl)-5-methylthieno[3,2-b]pyridine-3-carboxylate*. Isopropyl 7-(2-(2-bromoethoxy)-5-chlorophenyl)-5-methylthieno[3,2-b]pyridine-3-carboxylate (159.0 mg, 0.340 mmol, 1.0 eq), 2-methyl-4-oxo-7-(trifluoromethyl)-3,4-dihydroquinazoline-5-carbonitrile (86.0 mg, 0.340 mmol, 1.0 eq), $K_2CO_3$ (141.0 mg, 1.02 mmol, 3.0 eq), and anhydrous DMF (0.07 M, 5 mL) were added to an oven-dried round bottom flask. A condenser was attached and the mixture was heated overnight at 60 °C. The reaction mixture was cooled to room temperature and diluted with EtOAc, then transferred to a separatory funnel. Water was added, the layers mixed and separated, and the aqueous layer extracted with EtOAc (15 mL x 3). The organic layer was dried over anhydrous $Na_2SO_4$, filtered and concentrated to afford a yellow oil which was purified by flash column chromatography ($R_f$: 0.4, 0-100% EtOAc in hexanes) to afford the product as white solids (163.0 mg, 75%). $^1$H NMR (400 MHz, DMSO-$d_6$) δ 8.37 (d, $J$ = 1.7 Hz, 1H), 8.23 (s, 1H), 8.00 (d, $J$ = 1.6 Hz, 1H), 7.55 (dd, $J$ = 8.9, 2.7 Hz, 1H), 7.35 (d, $J$ = 2.7 Hz, 1H), 7.31 (d, $J$ = 9.0 Hz, 1H), 7.23 (s, 1H), 5.19 (p, $J$ = 6.2 Hz, 1H), 4.38 (t, $J$ = 4.9 Hz, 2H), 4.25 (d, $J$ = 4.9 Hz, 2H), 2.62 (s, 3H), 1.77 (s, 3H), 1.38 (d, $J$ = 6.3 Hz, 6H).

**2**, step j: *7-(5-chloro-2-(2-(5-cyano-2-methyl-4-oxo-7-(trifluoromethyl)quinazolin-3(4H)-yl)ethoxy)phenyl)-5-methylthieno[3,2-b]pyridine-3-carboxylic acid*. Isopropyl 7-(5-chloro-2-(2-(5-cyano-2-methyl-4-oxo-7-(trifluoromethyl)quinazolin-3(4H)-yl)ethoxy)phenyl)-5-methylthieno[3,2-b]pyridine-3-carboxylate (163.0 mg, 0.254 mmol, 1.0 eq) was added to a round bottom flask followed by anhydrous DCE (0.05 M, 5 mL). Then, trimethyltin hydroxide (689.6 mg, 3.81 mmol, 15.0 eq) was added and the mixture stirred under $N_2$ overnight at 90 °C. The reaction mixture was cooled to room temperature, diluted with DCM, then filtered through celite and washed with DCM. The filtrate was concentrated to afford a pale-yellow oil which was purified by flash column chromatography ($R_f$: 0.4, 0-10% MeOH in DCM) to afford **1** as white solids (107.1 mg, 66%).

**Scheme 2.** Synthesis of **3**

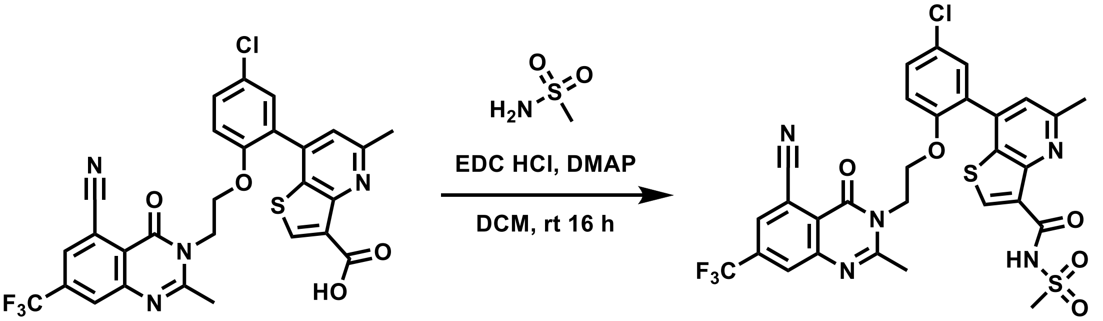


**2**

*7-(5-chloro-2-(2-(5-cyano-2-methyl-4-oxo-7-(trifluoromethyl)quinazolin-3(4H)-yl)ethoxy)phenyl)-5-methyl-N-(methylsulfonyl)thieno[3,2-b]pyridine-3-carboxamide*. To a round bottom flask was added **1** (50 mg, 0.835 mmol, 1.0 eq), methanesulfonamide (19.9 mg, 0.209 mmol, 2.5 eq), DMAP (25.5 mg, 0.209 mmol, 2.5 eq), and anhydrous DCM (0.03 M, 3 mL). Then, EDC · HCl (32.0 mg, 0.167 mmol, 2.0 eq) was added and the mixture stirred overnight at room temperature under $N_2$. The reaction mixture was concentrated and directly used for flash column chromatography to afford the product as pale-yellow solids, which was purified further using preparative RP-HPLC Method A to afford **2** as white solids (27.0 mg, 48%).

#### $^1$H NMR Spectra of Intermediates

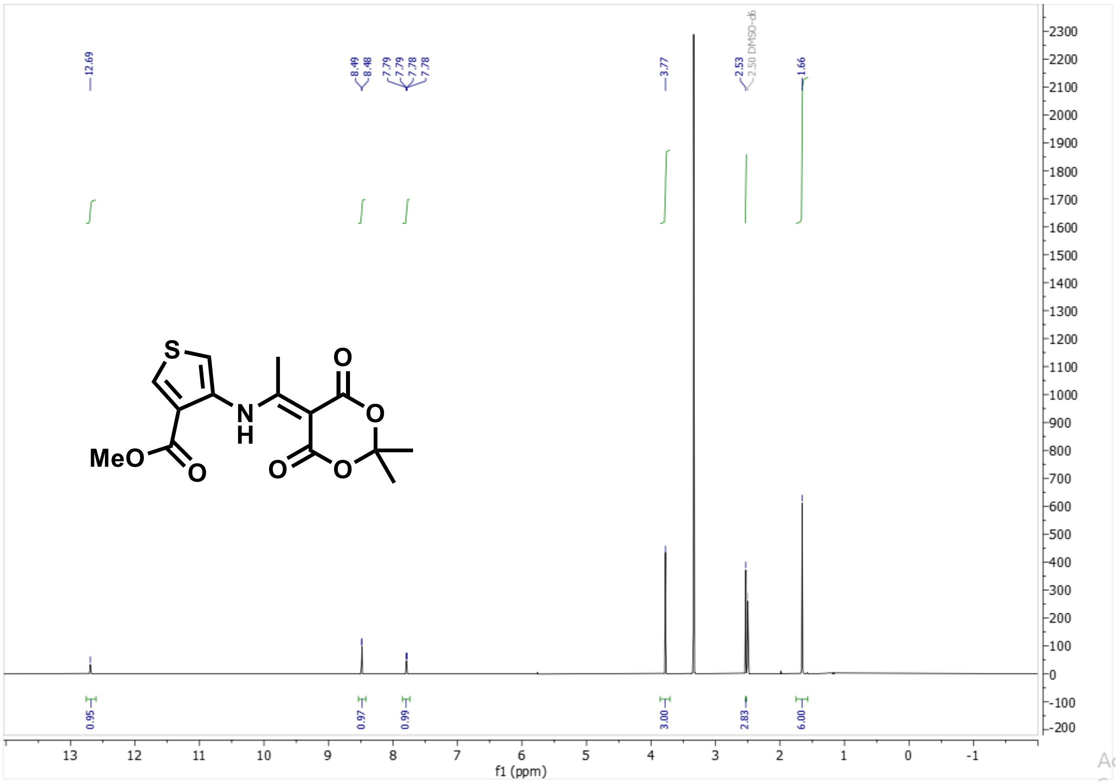


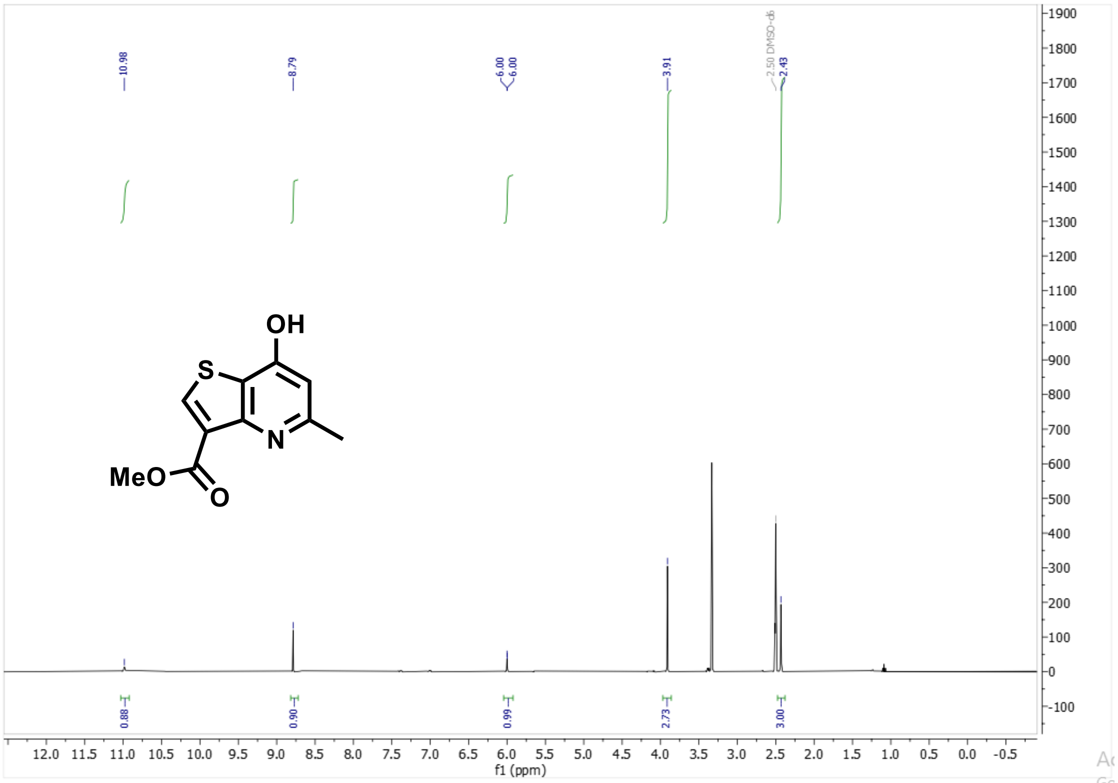

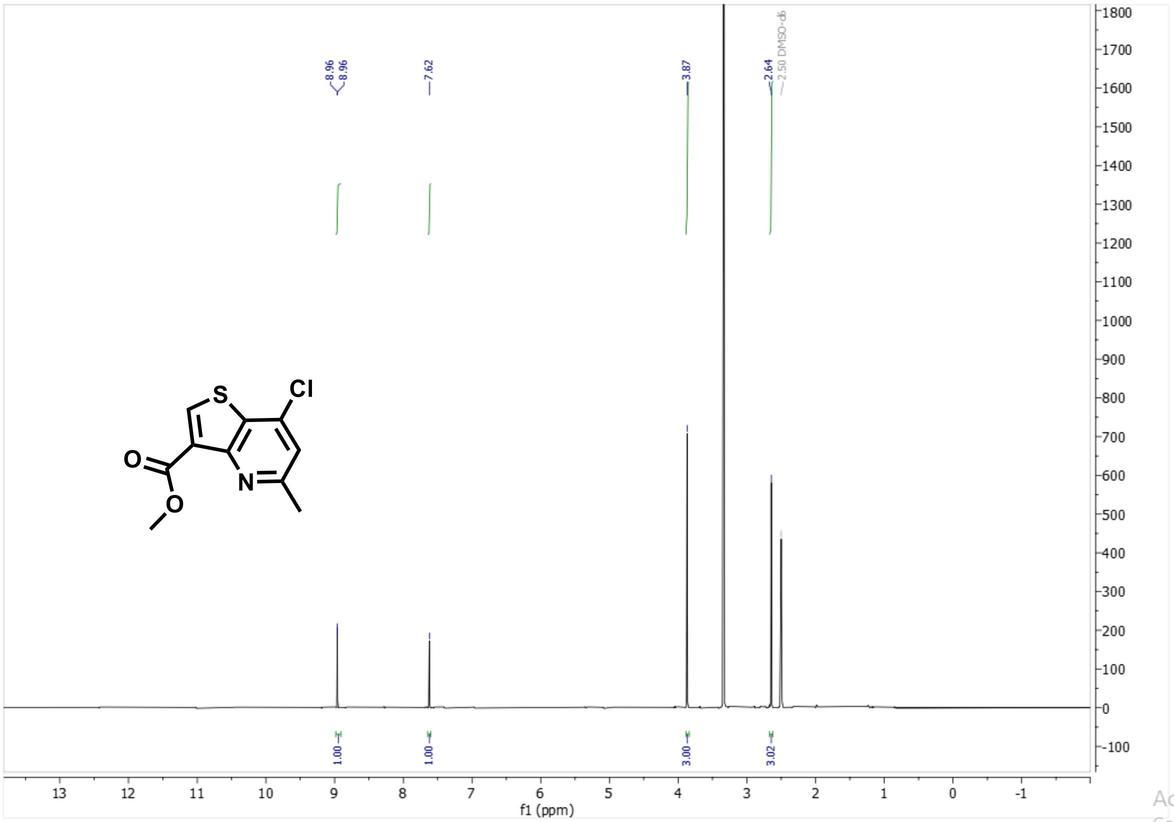
8.96
8.96
7.62
3.87
2.64
2.50 DMSO-d6
1.00
1.00
3.00
3.02
f1 (ppm)
Cl
S
O
O
N

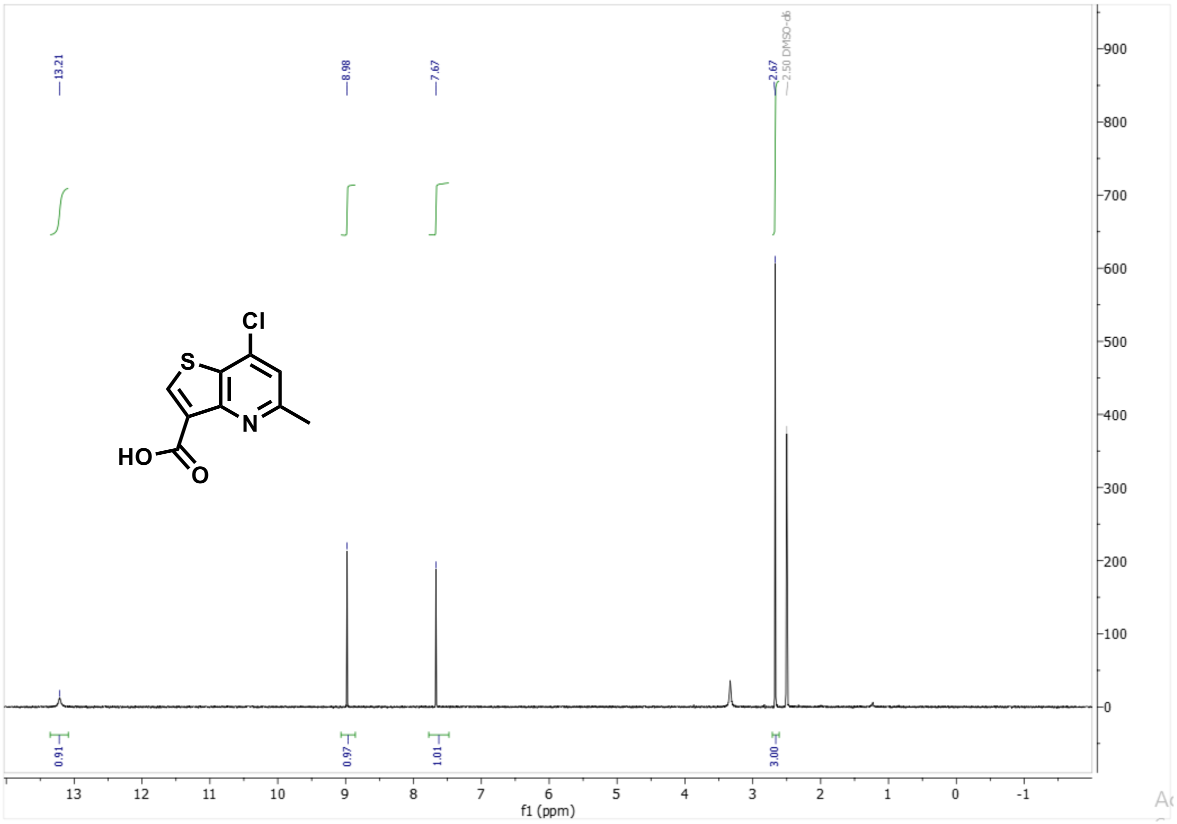
13.21
8.98
7.67
2.67
2.50 DMSO-d6
0.91
0.97
1.01
3.00
f1 (ppm)
Cl
S
N
HO
O

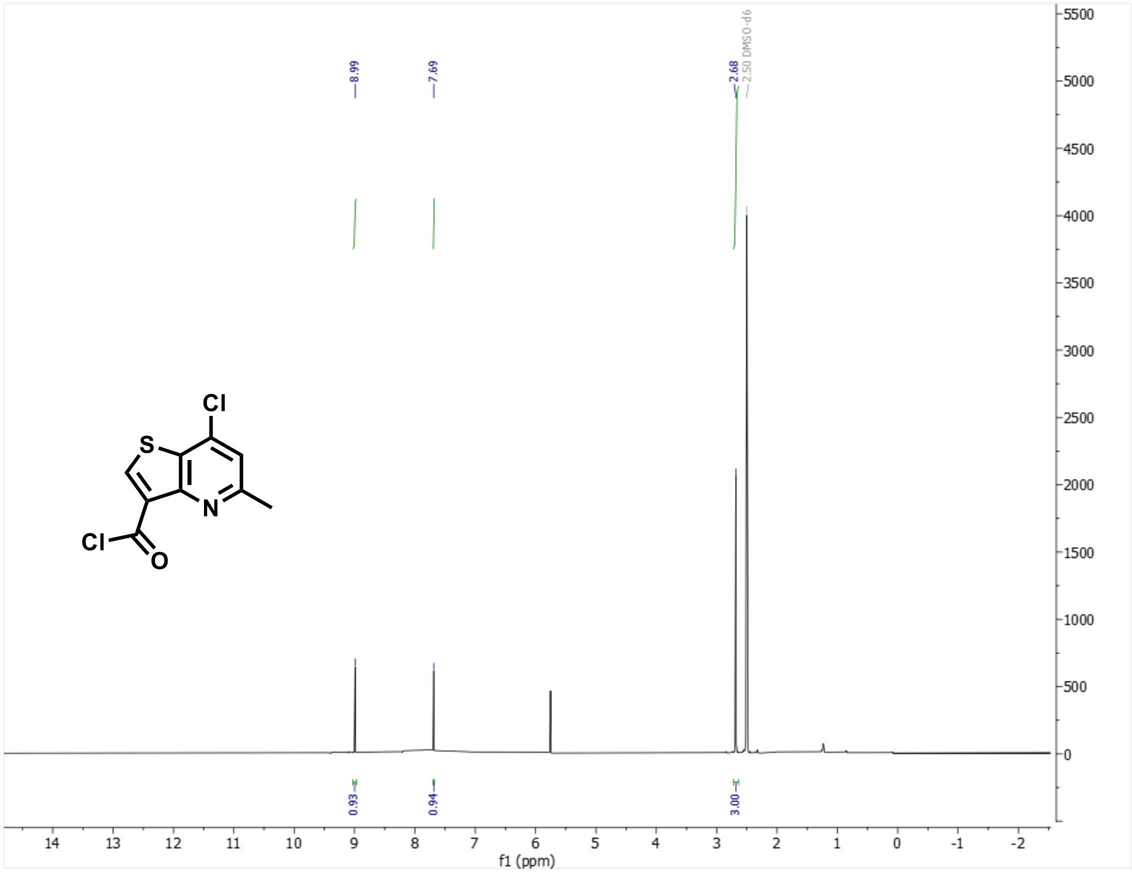

8.99
7.69
2.68
2.50 DMSO-d6
Cl
S
N
Cl
O
0.93
0.94
3.00
f1 (ppm)


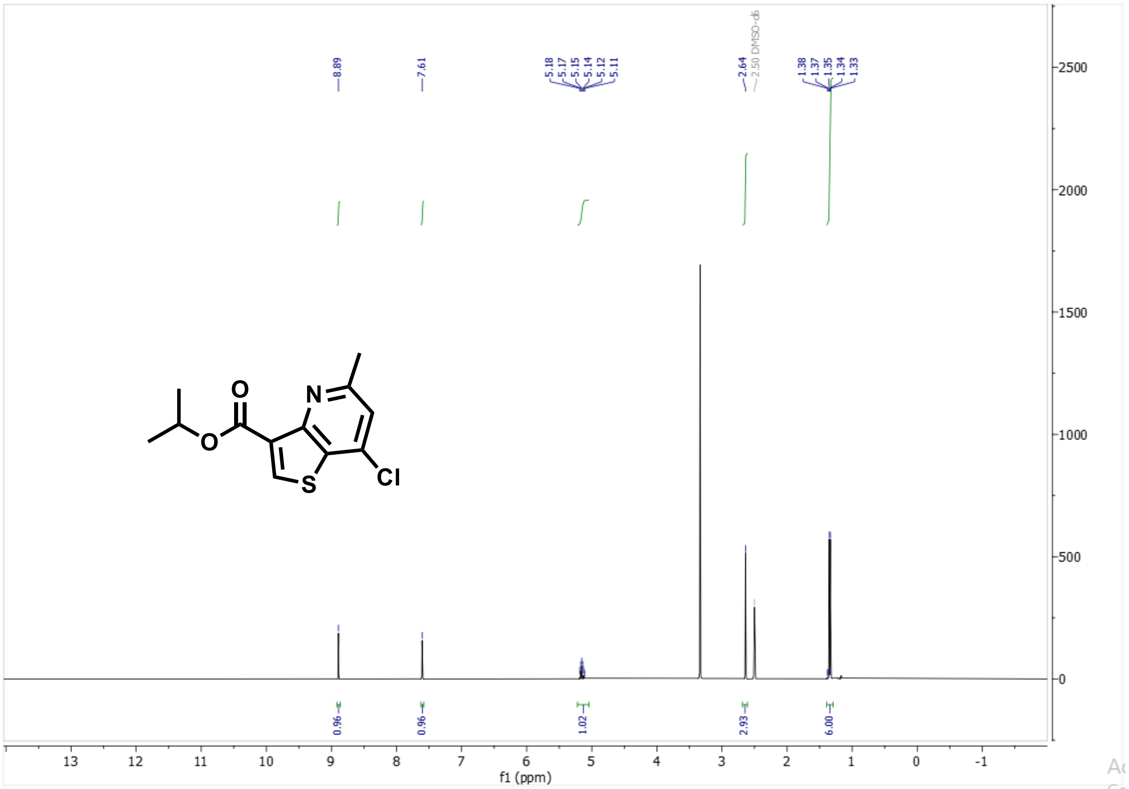

8.89
7.61
5.18
5.17
5.15
5.14
5.12
5.11
2.64
2.50 DMSO-d6
1.38
1.37
1.35
1.34
1.33
O
N
O
S
Cl
0.96
0.96
1.02
2.93
6.00
f1 (ppm)

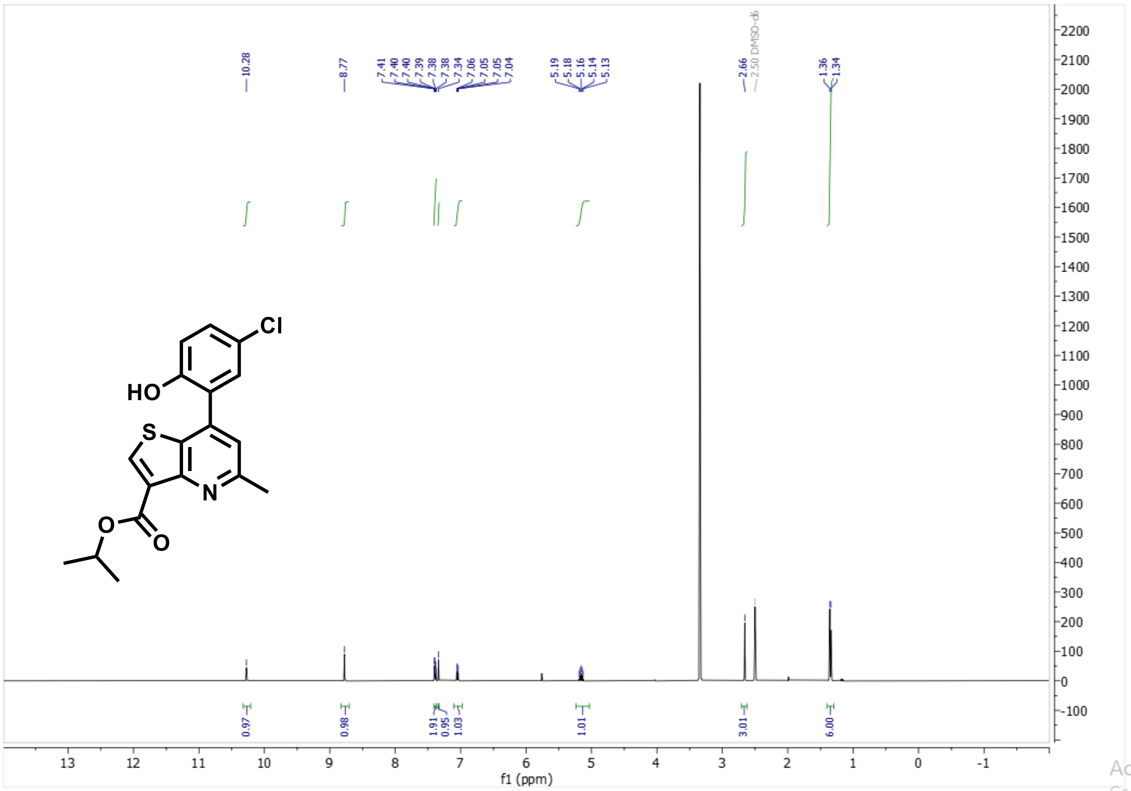
Cl
HO
S
N
O
O
10.28
8.77
7.41
7.40
7.40
7.39
7.38
7.38
7.34
7.06
7.05
7.05
7.04
5.19
5.18
5.16
5.14
5.13
2.66
2.50 DMSO-d6
1.36
1.34
0.97
0.98
1.91
0.95
1.03
1.01
3.01
6.00
f1 (ppm)

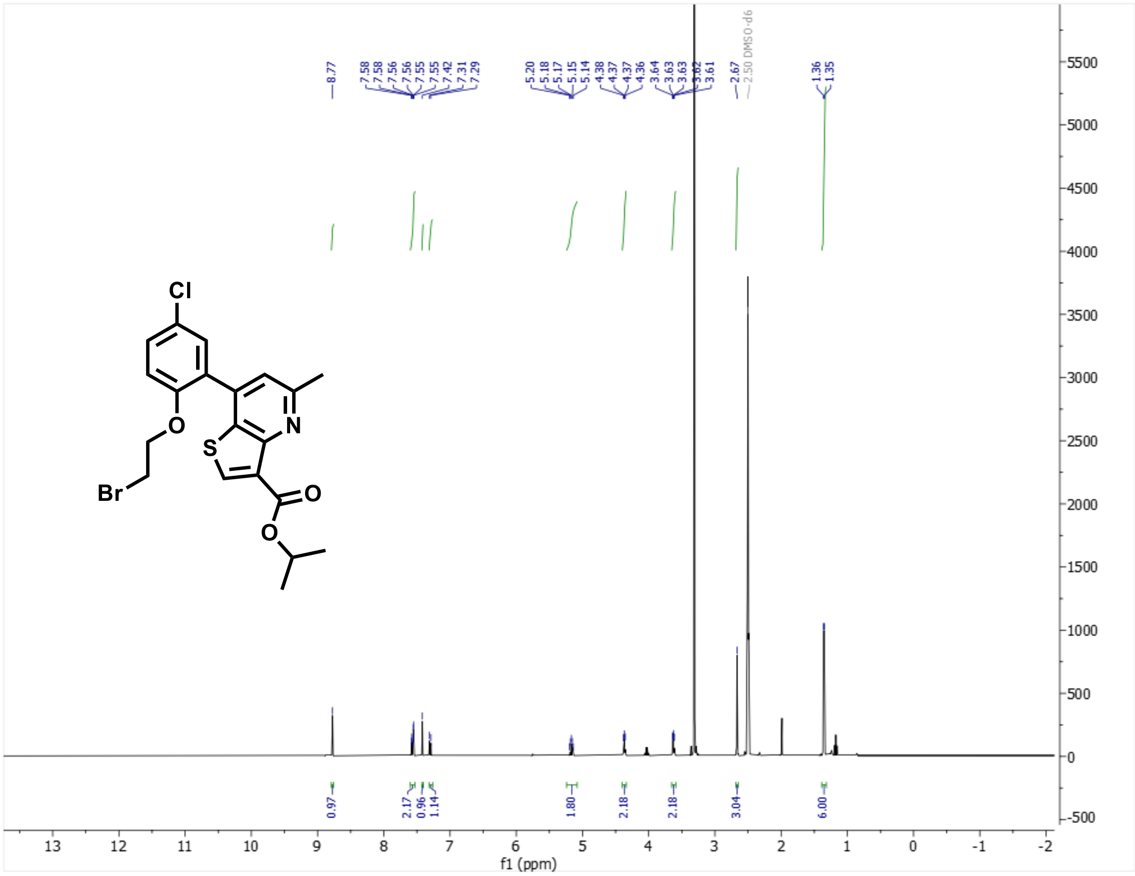
Cl
O
Br
S
N
O
O
8.77
7.58
7.58
7.56
7.56
7.55
7.55
7.42
7.31
7.29
5.20
5.18
5.17
5.15
5.14
4.38
4.37
4.37
4.36
3.64
3.63
3.63
3.62
3.61
2.67
2.50 DMSO-d6
1.36
1.35
0.97
2.17
0.96
1.14
1.80
2.18
2.18
3.04
6.00
f1 (ppm)

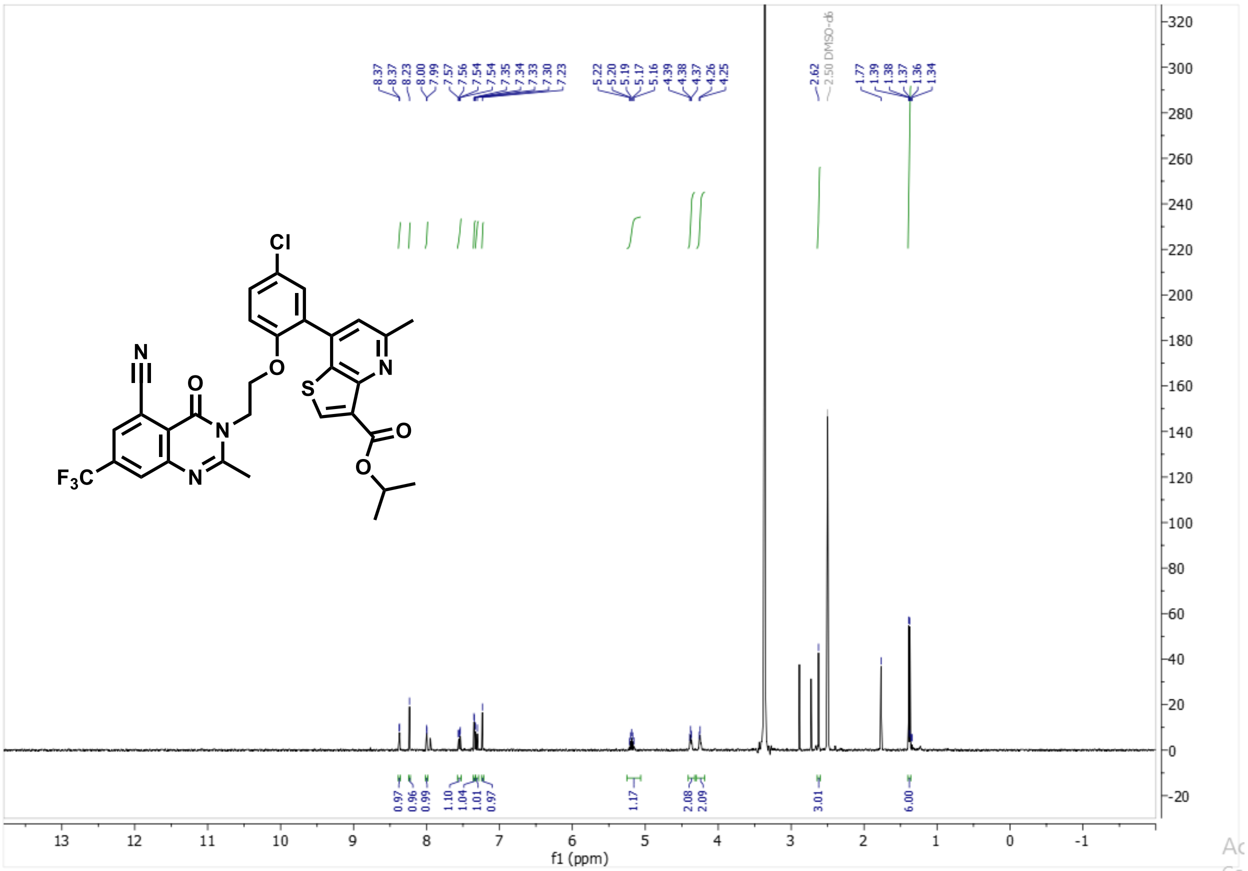
